# Whole brain mapping of orexin receptor mRNA expression visualized by branched *in situ* hybridization chain reaction

**DOI:** 10.1101/2023.10.22.563416

**Authors:** Yousuke Tsuneoka, Hiromasa Funato

**Affiliations:** Department of Anatomy, Faculty of Medicine, Toho University, Tokyo, Japan; International Institutes for Integrative Sleep Medicine (WPI-IIIS), University of Tsukuba, Ibaraki, Japan

**Keywords:** orexin, receptor, sleep, *in situ* hybridization chain reaction, whole-brain, dopamine, acetylcholine, serotonin, histamine, fluorescent in situ hybridization

## Abstract

Orexins, which are produced within neurons of the lateral hypothalamic area, play a pivotal role in the regulation of various behaviors, including sleep/wakefulness, reward behavior, and energy metabolism, via orexin receptor type 1 (OX1R) and type 2 (OX2R). Despite the advanced understanding of orexinergic regulation of behavior at the circuit level, the precise distribution of orexin receptors in the brain remains unknown. Here, we develop a new branched *in situ* hybridization chain reaction (bHCR) technique to visualize multiple target mRNAs in a semiquantitative manner, combined with immunohistochemistry, which provided comprehensive distribution of orexin receptor mRNA and neuron subtypes expressing orexin receptors in mouse brains. Only a limited number of cells expressing both *Ox1r* and *Ox2r* were observed in specific brain regions, such as the dorsal raphe nucleus and ventromedial hypothalamic nucleus. In many brain regions, *Ox1r*-expressing cells and *Ox2r*-expressing cells belong to different cell types, such as glutamatergic and GABAergic neurons. Moreover, our findings demonstrated considerable heterogeneity in *Ox1r*- or *Ox2r*-expressing populations of serotonergic, dopaminergic, noradrenergic, cholinergic, and histaminergic neurons. The majority of orexin neurons did not express orexin receptors. This study provides valuable insights into the mechanism underlying the physiological and behavioral regulation mediated by the orexin system, as well as the development of therapeutic agents targeting orexin receptors.

**Significance statement:** The neuropeptide orexin regulates sleep and other behaviors through its receptors, OX1R and OX2R, which are targets for the development of therapeutic agents for sleep and related disorders. However, the cellular distribution of orexin receptors in the brain is only partially known. We applied a newly developed branched *in situ* hybridization chain reaction (bHCR) technique and conducted a whole-brain mapping of orexin receptor mRNA expression in the brain with neuron subtype markers. Few cells expressed both OX1R and OX2R, and OX1R and OX2R were expressed in the different neuronal subtypes in many brain regions. This study fills an important gap in understanding and modulating the orexin system.

## 1 Introduction

The orexin system, which consists of the neuropeptides, orexin A and orexin B, and their receptors, orexin receptor type 1 (OX1R) and type 2 (OX2R), serves as a core neural circuit for sleep and wakefulness (1, 2). In addition, orexinergic neurons are involved in multiple aspects of behaviors and metabolism, such as motivation, addiction, stress response, food intake, energy metabolism, glucose metabolism, sympathetic nervous regulation, and pain sensation (3–8).

Orexins, also known as hypocretins, are highly conserved neuropeptides among vertebrates and are exclusively produced in specific neurons of the lateral hypothalamic area (LHA) (9, 10). Orexin A and B are processed from the common precursor peptide, prepro-orexin. Orexin A is a 33 amino acid peptide with two intrachain disulfide bonds formed between cysteine residues. Orexin B is a 28 amino acid peptide with an amidated carboxyl-terminus. OX1R preferentially binds to orexin A, whereas OX2R binds to orexin A and B with similar high affinity (10). OX1R and OX2R are G protein-coupled receptors (GPCRs) and have distinct roles in sleep/wake regulation and energy metabolism (11–14). Orexin receptors were thought to be coupled primarily to Gq and partially to Gi. However, recent comprehensive analysis indicates that both OX1R and OX2R couple promiscuously to Gq, Gs, and Gi (15–17).

Orexinergic neurons send their axons throughout the central nervous system. Optogenetic and chemical genetic analysis identified target regions of orexinergic neuron projections that regulate sleep/wakefulness (18–20), aggression (21), fear (22), and glucose metabolism (14). However, in contrast to the advanced circuit-level understanding of the orexin system, the detailed distribution of OX1R and OX2R in the brain has not been shown. The rough distribution of orexin receptors was reported in rat brains by radiolabeled probes approximately 20 years ago and is still the most comprehensive information available (23, 24). Cell resolution information of OX1R and OX2R has been available only for brain regions that abundantly express orexin receptors, such as the amygdala, tuberomammillary nucleus, raphe nuclei, locus coeruleus, and laterodorsal tegmental nucleus (14, 25–28). At this time, the Allen Brain Atlas fails to show positive signals for orexin receptors and reliable antibodies for OX1R and OX2R that work for immunohistochemistry are not available.

To obtain whole-brain information on cells expressing orexin receptors in a semiquantitative manner, we developed a new branched *in situ* hybridization chain reaction (HCR) technique (Fig. 1A, S1), by combining short hairpin HCR with split-initiator probe (29), and hyperbranched HCR systems (30). This branched HCR (bHCR) can visualize not only multiple target mRNAs but also proteins combined with immunohistochemistry because bHCR does not require proteinase K treatment, which reduces the immunogenicity of proteins. Using bHCR combined with immunohistochemistry, we characterized the distribution and transmitters of cells expressing orexin receptors throughout the brain. This study also provides information on orexin receptor expression in monoaminergic and cholinergic neurons. We discuss orexin receptor expression in terms of different behavioral modalities such as sleep/wakefulness, reward behavior, feeding, energy metabolism, and the autonomic nervous system.

**Fig. 1.**
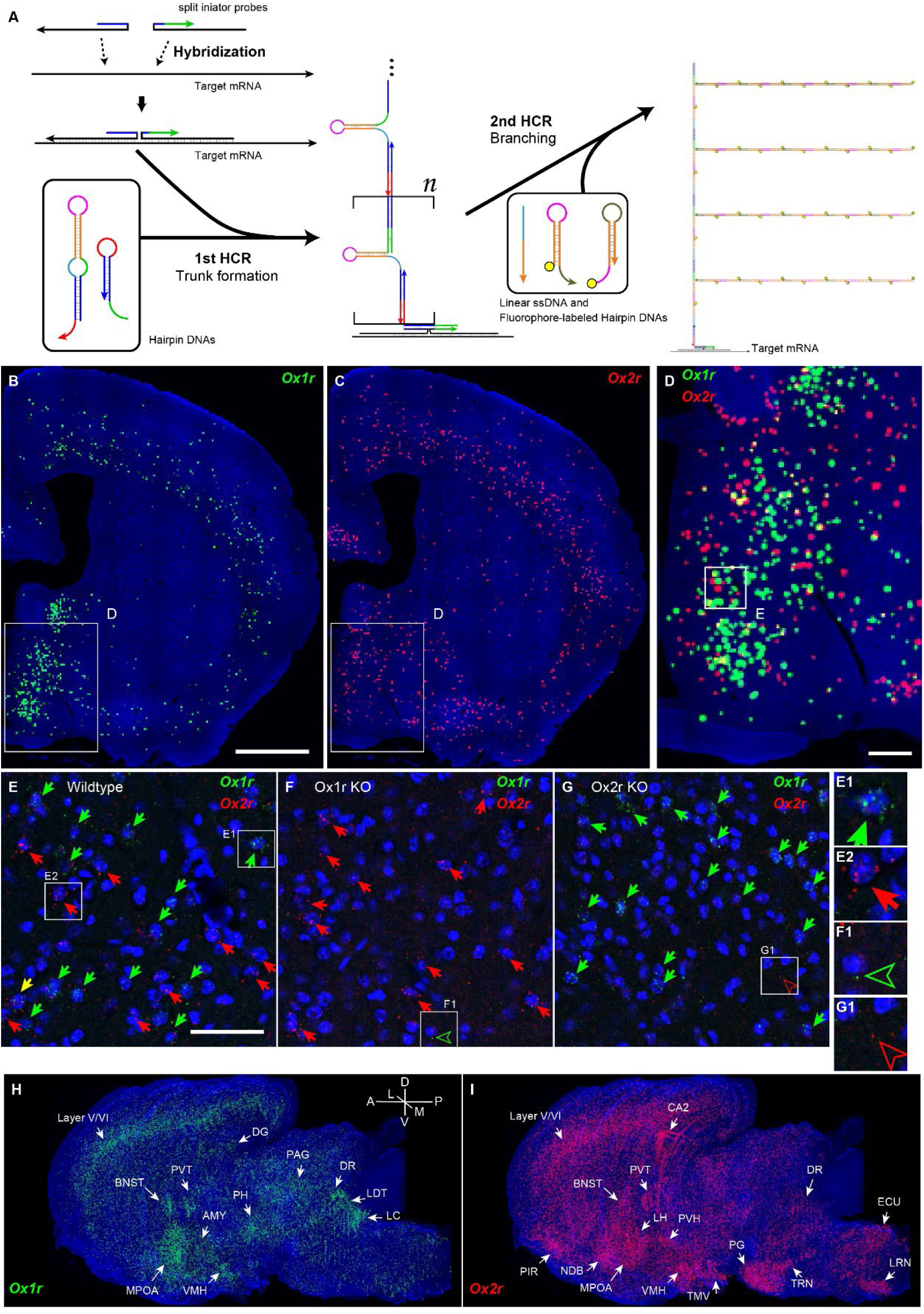
Outline of branched HCR (bHCR) amplification and its specificity. (A) A pair of split-initiator probes formed an initiator sequence to start the 1st HCR when the probes hybridized with the target mRNAs in the correct position. In the 1st HCR, the long hairpin DNA and the short hairpin DNA formed a “trunk” on the mRNA. Subsequently, the remaining hairpin domains of the trunk were hybridized with linear ssDNA to expose initiator sequences to start the polymerization of fluorophore-labeled hairpin DNAs (2nd HCR). For details, please see Supplemental Figure S1. (B-D) Reconstructed distribution of *Ox1r* and *Ox2r* expressing cells around bregma 0 level. The receptor-expressing cells are indicated by green (*Ox1r*) or red (*Ox2r*) circles. The large circles denote the cells with high expression. The rectangles in B and C indicate the region shown in D, and the rectangle in D indicates the magnified view in E. (E-G) Representative microphotographs of *Ox1r-* and *Ox2r-*stained sections of the MPOA of wildtype, *Ox1r* KO and *Ox2r* KO mice. Cell nuclei are shown in blue. Arrows indicate the receptor-expressing cells. The outlined arrowheads in F and G show a single granule of noise. Scale bars: 1 mm (B), 200 µm (D), and 50 µm (E). (H-I) Reconstructed 3-dimensional images showing receptor expression of *Ox1r* (H) and *Ox2r* (I). The regions showing notable expression are indicated by arrows.

## 2 Results

To obtain whole-brain information on orexin receptor expression at a cellular resolution in a semiquantitative manner, we developed bHCR, which consists of two steps of HCRs (Fig. 1A, S1). In the first step, the hybridization of a pair of split-initiator probes to target mRNAs triggers HCRs of double-stem hairpin and short hairpin, which forms a “hairpin-containing DNA trunk” (29, 31). In the second step, assist oligo DNAs bind to double-stem hairpins of the DNA trunk to stretch self-hybridization of double-stem hairpins, which allows double-stem hairpins to hybridize to fluorophore-labeled hairpin DNAs. Then, HCRs further occur on the trunk to form fluorescently tagged branches.

bHCR visualized *Ox1r* and *Ox2r* signals throughout the brain in a region-specific manner (Fig. 1B-D, H, I). Since our HCR system can visualize a single mRNA molecule (29), the number of fluorescent granules produced by HCRs is thought to roughly correlate with the expression level of the target mRNAs. The locus coeruleus (LC) and ventral part of the tuberomammillary nucleus (TMV) contain cells expressing a large number of fluorescent granules for *Ox1r* and *Ox2r*, respectively (Fig. 1H, I, also see Fig. 6E, H) as previously reported (24, 26). However, the majority of orexin receptor-positive cells exhibited fewer than five fluorescent granules around the nucleus, reflecting their low expression (Fig. 1E). The specificity of the fluorescent granules for *Ox1r* and *Ox2r* was verified using *Ox1r-* and *Ox2r-*deficient brains (Fig. 1E-G). A few fluorescent granules were sparsely and evenly distributed in all brain regions, which may be artifacts or background signals (Fig. 1E-G). Therefore, we considered a cell with two or more fluorescent granules to be a positive cell. Depending on the number of fluorescence granules, positive cells were further classified into low or high expression. We drew small circles for cells with low expression and large circles for cells with high expression on original microscopic images (Fig. 1B, C, D; Fig. S2). The reconstructed 3-D images of cells expressing *Ox1r* and *Ox2r* showed that the density of orexin receptor-positive cells largely varied among brain regions (Fig. 1H, I). Importantly, many brain regions had *Ox1r*-positive cells and *Ox2r*-positive cells, but *Ox1r*-and *Ox2r*-double positive cells were rarely observed (Fig. 1 B-D), except for some brain regions, such as the paraventricular thalamus (PVT), VMH and dorsal raphe nucleus (DRN), where a large number of double-positive cells were found (see below).

To make a whole-brain map of *Ox1r-* and *Ox2r-*expressing cells, we used a complete series of 40-μm thick coronal sections from a single male mouse. For every three sections, bHCR for *Ox1r* and *Ox2r*, bHCR for *Vgat, Vglut1*, and *Vglut2*, and bHCR for *Chat* with immunostaining for calbindin and TH were performed (Fig. S2). In addition, to characterize *Ox1r*- or *Ox2r*-expressing cells, bHCR for *Ox1r* or *Ox1r* was combined with bHCR for *Vgat*, *Vglut2*, or Chat, followed by immunostaining of either tyrosine hydroxylase (TH), histidine decarboxylase (HDC), 5-HT or orexin. Regional expression of the orexin receptor, coexpression of glutamatergic or GABAergic markers (*Vglut2* or *Vgat*), and distribution of area markers (*Vglut1, Vglut2*, *Vgat*, *Chat*, TH, and Calbindin) are summarized in Supplementary Tables S1,S2.

### 2.1 Cerebrum

In the isocortex, a large number of *Ox1r*-positive cells and *Ox2r*-positive cells were observed in layers V and VI (Fig, 2A-D, Fig. S2). *Ox1r*- or *Ox2r*-positive cells included glutamatergic cells and GABAergic cells. A small number of *Ox1r*- and *Ox2r*-double positive cells were found in layer VI (Fig 2A-D, Fig. S2). In layers II/III, a moderate number of *Ox2r*-positive cells were observed, which were *Vgat*-positive cells, while a small number of *Ox1r*-positive cells were observed, which were *Vgat*-negative cells (Table S1). Few orexin receptor-expressing cells were found in layers I and IV.

**Fig. 2.**
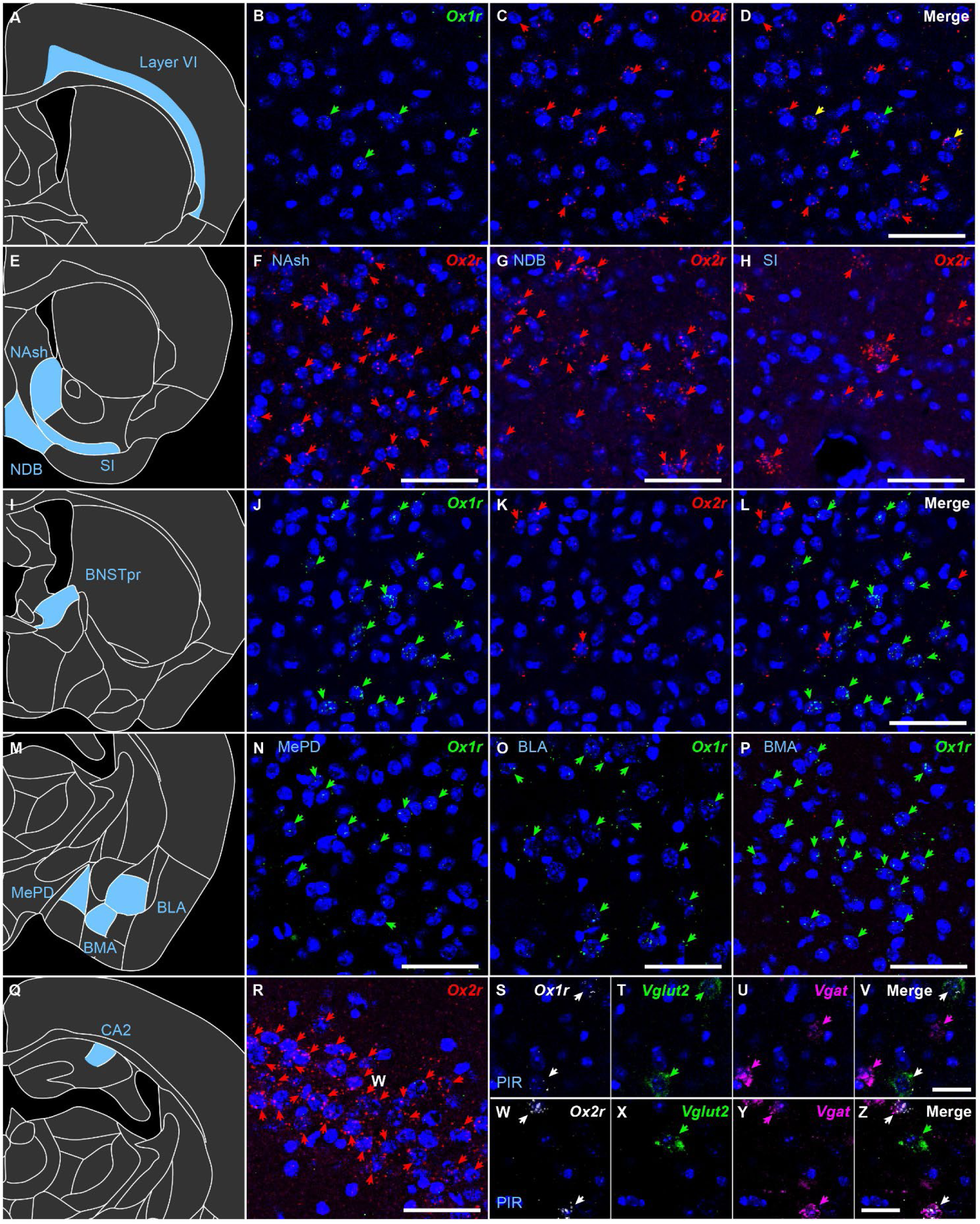
Representative expression of *Ox1r* and *Ox2r* in the cerebrum. (A-D) *Ox1r* and *Ox2r* expression in isocortex layer VI. (E-H) *Ox2r* expression in the cell region of the nucleus accumbens (F), the nucleus of the diagonal band (G), and the substantia innominata (H). (I-L) *Ox1r* and *Ox2r* expression in the BNST principal nucleus. (M-P) Ox1r expression in the medial amygdala, posterodorsal (N), basolateral amygdala (O), and basomedial amygdala (P). (Q, R) *Ox2r* expression in the CA2 region of the hippocampus. Green, red, and yellow arrows denote *Ox1r*, *Ox2r,* and both receptors-expressing cells. (S-Z) *Ox1r*, *Ox2r*, *Vglut2,* and *Vgat* expression in the piriform cortex (PIR). Scale bars: 50 μm (A-R), 25 μm (V, Z).

In the anterior olfactory area, such as the anterior olfactory nucleus (AON), dorsal taenia tecta (DTT), ventral taenia tecta (VTT), and olfactory tubercle (OT), *Ox2r*-positive cells were abundant and *Ox1r*-positive cells were scarce (Fig. S2). The majority of *Ox2r*-positive cells were *Vgat*-positive. In the piriform area, anterior to the bregma level, abundant *Ox2r*-positive cells and few *Ox1r*-positive cells were observed, but in the piriform area posterior to the bregma level, the number of *Ox2r*-positive cells decreased, and the number of *Ox1r*-positive cells increased (Fig. 2S-Z, S2 Panel 43-46). In the anterior cortical amygdala (ACo) and nucleus of the lateral olfactory tract (LOT), *Ox1r* was expressed in *Vglut2*-positive neurons. In the posterolateral cortical amygdala (PLCo), *Ox1r*-positive cells were rare. Similar to the anterior olfactory area, more than half of *Ox2r*-positive cells were *Vgat*-positive in the cortical amygdala and adjacent nuclei (Fig. 2P).

In general, the striatum and related regions contained many *Ox2r*-positive cells with strong expression and fewer *Ox1r*-positive cells. A small population of GABAergic neurons in the caudoputamen (CPu) expressed *Ox2r*. In the CPu, there were scattered *Vgat*- and *Chat*-double-positive large cells, one-third of which strongly expressed *Ox2r*. The nucleus accumbens expressed *Ox2r* but not *Ox1r*. The nucleus accumbens shell (Nash) was one of the most densely populated regions in the brain with cells expressing *Ox2r* (Fig. S2 Panel 13-28). The diagonal band (NDB), magnocellular nucleus (MN), and globus pallidus (GP) were enriched in *Vgat*-positive cells highly expressing *Ox2r* (Fig. 2 E-H). Regarding the septal area, *Ox2r* expression was found mainly in *Vgat*-positive cells in the medial septum (MS), while *Ox2r* expression was found in *Vglu2*-positive cells in the triangular septal nucleus (TRS), septofimbrial nucleus (SF), and lateral septum (LS).

For the extended amygdala, the principal nucleus of the BNST (BNSTpr) contained *Ox1r*-expressing cells, which are *Vgat*-positive neurons, while cells expressing *Ox2r* were mainly positive for *Vglut2* (Fig. 2. I-L). In the BNST nuclei, except for BNSTpr, *Ox1r*- and *Ox2r*-positive cells were *Vgat*-positive. The posterodorsal and posteroventral medial amygdala (MePD, MePV) were rich in *Ox1r*-positive cells, and they were mainly *Vglut2*-positive (Fig. 2 M-P). The central amygdala (CEA) expressed *Ox2r* but not *Ox1r*, and *Ox2r*-positive cells were *Vgat*-positive. The anterodorsal and anteroventral medial amygdala (MeAD, MeAV) contained *Ox1r*-positive cells and *Ox2r*-positive cells, which were mainly *Vgat*-positive.

In the endopiriform nucleus (Epi) and claustrum (CLA), the expression levels of *Ox1r* and *Ox2r* were relatively low. Among the lateral, basolateral, and basomedial amygdala, area boundaries can be easily divided by *Vglut1* and *Vglut2* expression (Fig. S2 Panel 74-80). In the basolateral amygdala (BLA) and basomedial amygdala (BMA), a dense distribution of *Ox1r*-positive glutamatergic cells was observed (Fig. 2 M-P), while a relatively small number of *Ox2r*- and *Vgat*-positive cells were distributed in the posterior part of the basolateral amygdala (BLP), the posterior part of the basomedial amygdala (BMP), lateral amygdala (LA) and amygdala hippocampal area (Ahi).

In the hippocampus, there was a high density of *Ox2r*-positive cells in the CA2 region that were *Vgat*-negative and appeared to be pyramidal cells and a moderate density of *Ox2r*-positive cells in the dentate gyrus, CA1, CA3, and subiculum, which were *Vgat*-positive (Fig. 2 Q,R). *Ox1r* was expressed in *Vgat*-negative cells, which were localized in the polymorphic cell layer of the dentate gyrus and CA3 region. In the posterior part of the CA1, CA3, and subiculum, *Ox2r*-positive cells were *Vgat*-negative.

### 2.2 Thalamus

In the thalamus, *Ox1r* and *Ox2r* were moderately expressed in the midline regions, such as the paraventricular nucleus of the thalamus (PVT) and centromedian nucleus (CM), but low in other thalamic regions. Since the thalamus is composed of excitatory neurons, *Ox1r* expression was also found in *Vglut2*-positive cells of the anteromedial nucleus (AM), paraxiphoid nucleus (PaXi), and subparafascicular nucleus (SPF). *Ox2r*-positive cells were more frequently found around midline cells, especially in the parataenial nucleus (PT) and xiphoid nucleus (Xi). In the posterior group, *Ox2r*-positive cells were found in the subparafascicular area (SPA), SPF, and perireunensis nucleus (PR). The anterior part of the PVT, which is one of the major target areas of orexin neurons to promote wakefulness (32), moderately expressed *Ox1r* and *Ox2r* and contained many *Ox1r*- and *Ox2r*-double-positive cells (Fig. 3A-D). Regarding GABAergic areas surrounding the thalamus, *Ox1r* and *Ox2r* were not expressed in the reticular thalamus (RT) but were expressed in *Vgat*-positive cells of the intergeniculate leaflet (IGL) and pregeniculate nucleus (PrG) (Fig. 3 E,F). The medial and lateral habenula expressed *Ox2r* but not *Ox1r*.

**Fig. 3.**
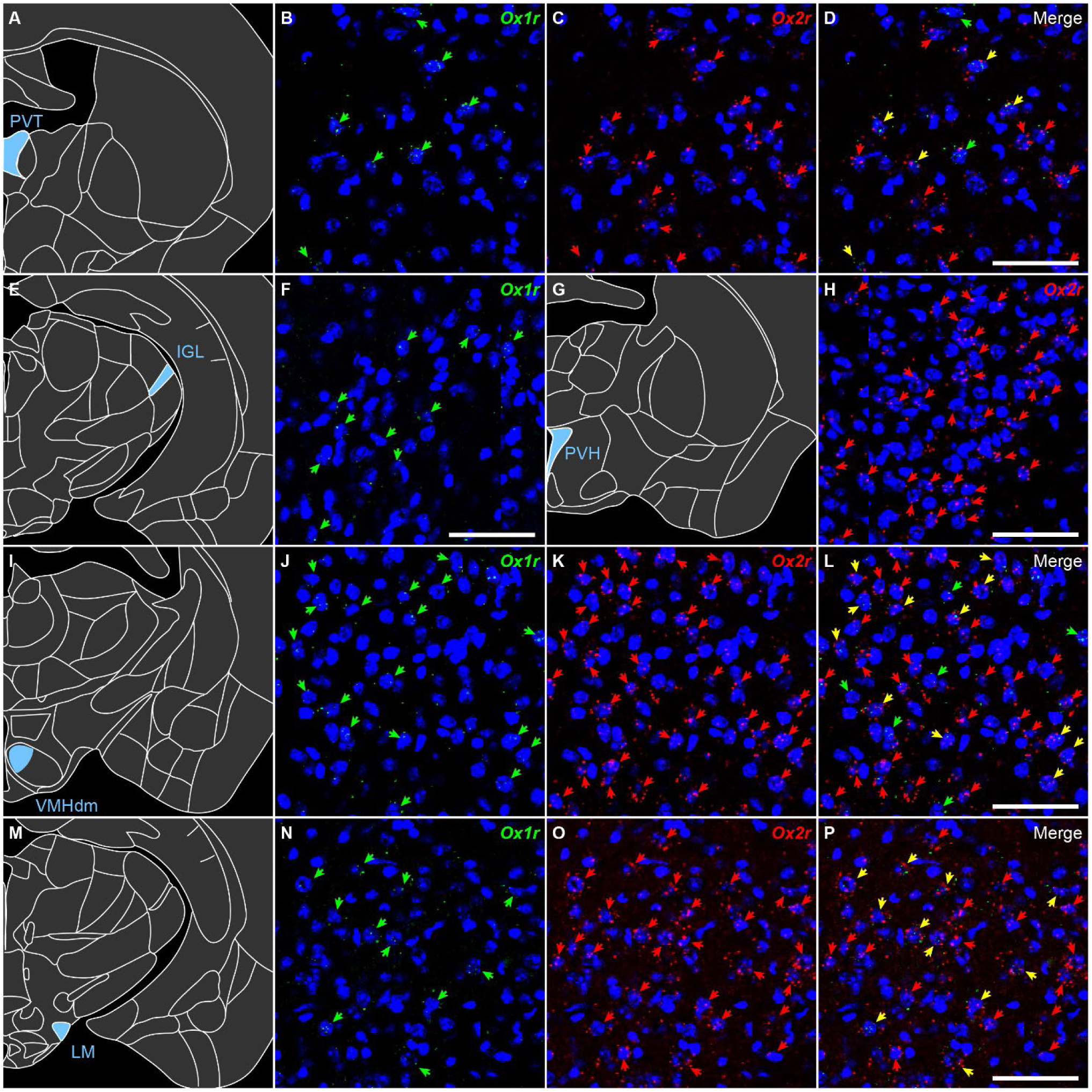
Representative expression of *Ox1r* and *Ox2r* in the diencephalon. (A-D) *Ox1r* and *Ox2r* expression in the periventricular thalamus. (E, F) *Ox1r* expression in the intergeniculate leaflet. (G, H) *Ox2r* expression in the paraventricular hypothalamic nucleus. (I-L) *Ox1r* and *Ox2r* expression in the VMH, dorsomedial. (M-P) *Ox1r* and *Ox2r* expression in the lateral mammillary nucleus. Green, red, and yellow arrows denote *Ox1r-*, *Ox2r-,* and both receptor-expressing cells. Scale bars: 50 μm.

### 2.3 Hypothalamus

Consistent with the vicinity of orexin neurons localized in the LHA and dense orexin fiber projections in the hypothalamus, almost all hypothalamic areas expressed either *Ox1r* or *Ox2r*, except for the suprachiasmatic nucleus (SCN) and vascular organ of the lamina terminalis (VOLT). Importantly, only a few orexin A-positive neurons were positive for either *Ox1r* or *Ox2r* (5 of 101 cells).

In the preoptic region, the MPOA was one of the densest areas of receptor-expressing cells in the brain, although the expression intensity per cell was not high (Fig. 1 B-D). *Ox1r*-positive cells were found in *Vgat*-positive cells but not in *Vglut2*-positive cells. *Ox1r*-positive cells were distributed in the relatively posterior part of the MPOA. In contrast, *Ox2r*-positive cells were found in both *Vgat*- and *Vglut2*-positive cells and distributed in the relatively anterior part of the MPOA. In the median preoptic nucleus (MnPO) and lateral preoptic areas (LPOA), the expression was relatively low.

In the periventricular area, notable expression of *Ox2r* was found, although the expression intensity per cell was not high. In the paraventricular nucleus (PVN), *Ox2r* was expressed in many *Vglut2*-positive cells but not in *Vgat*-positive cells (Fig. 3G, H). The arcuate nucleus (ARC), which is composed of GABAergic neurons, expressed *Ox2r* but not *Ox1r*. In the dorsomedial nucleus (DMH), most of the *Ox2r*-positive cells were *Vgat*-positive.

*Ox1r*- and *Ox2r*-positive cells were abundant in the ventromedial nucleus (VMH), which is composed of excitatory neurons. In the dorsomedial part of the VMH, positive cell density was highest, followed by the ventrolateral part. Double-positive cells for *Ox1r* and *Ox2r* were frequently found in the dorsomedial part (Fig. 3I-L). In the LHA, a moderate number of receptor-expressing cells were observed in *Vgat*-positive and *Vglut2*-positive cells. In the posterior hypothalamus (PH), subthalamic nucleus (STN), and parasubthalamic nucleus (PST), many *Vglut2*-positive cells expressed *Ox2r*, but *Vgat*-positive cells did not express either *Ox1r* or *Ox2r*.

In the hypothalamus, cells strongly expressing *Ox2r* were most abundant in the premammillary and mammillary areas. In the dorsal premammillary nucleus (PMD) and ventral premammillary nucleus (PMV), strong and dense expression of *Ox2r*-positive cells was observed, whereas weakly expressed *Ox1r* mRNAs were observed in a moderate number of cells. The dorsal tuberomammillary nucleus (TMD) and ventral tuberomammillary nucleus (TMV) were highly dense in strongly *Ox2r*-expressing cells but did not express *Ox1r*. The lateral mammillary nucleus (LM), the lateral cluster of *Vglut2*-positive neurons in the mammillary body, was abundant in *Ox1r-* and *Ox2r-*double-positive cells, in addition to single-positive cells (Fig. 3M-P). In the supramammillary nucleus (SuM), which contains cells expressing both *Vglut2* and *Vgat* (33), a moderate number of *Ox2r*-positive cells were observed.

### 2.4 Midbrain

In the pretectal region, the anterior pretectal nucleus (APN) and medial pretectal nucleus (MPT) express the orexin receptor. In the APN, a large number of *Vglut2*- and *Vgat*-positive neurons expressed *Ox2r*, and a few of them expressed *Ox1r*.

In the ventral dopaminergic region, a moderate number of *Ox1r*- or *Ox2r*-expressing cells were found, but the expression per cell was low for both *Ox1r* and *Ox2r*. Among these regions, *Ox1r* expression was greater. In the retrorubral area (RR), substantia nigra pars compacta (SNc), and substantia nigra pars reticulata (SNr), approximately half of the receptor-expressing neurons were *Vgat*-positive, and the remaining neurons were *Vgat*- and *Vglut2*-negative and were presumed to be dopaminergic cells (Fig. S2 panel 108-120, also see “Monoamine neurons” section). In the ventral tegmental area (VTA), cells expressing either *Ox1r* or *Ox2r* were *Vgat*-positive cells, *Vglut2*-positive cells, or *Vgat*- and *Vglut2*-double-negative cells.

The periaqueductal gray (PAG) and adjacent nuclei were densely populated with *Ox1r*-positive cells and sparsely populated with *Ox2r*-positive cells. In the PAG, especially the lateral and ventrolateral subdivisions, approximately half of the neurons were *Ox1r*-positive. Most of the *Ox1r*-positive neurons were *Vgat*-positive and a few were *Vglut2*-positive. Conversely, *Ox2r*-positive neurons in the PAG were predominantly *Vglut2*-positive neurons. The highest expression of *Ox2r* per cell was observed in *Vglut2*-positive neurons of the medial accessory oculomotor nucleus (MA3, or Edinger-Westphal nucleus).

In the midbrain raphe nuclei, *Ox1r* and *Ox2r* were highly expressed. Whereas most *Ox1r*-positive neurons in the central linear nucleus (CLi) were *Vgat*-positive cells, most *Ox1r*-positive neurons in the rostral linear nucleus (RLi) were *Vglut2*-positive cells. The DRN is one of the regions with the highest per-cell expression of both *Ox1r* and *Ox2r*. The vast majority of receptor-expressing cells were negative for both *Vgat* and *Vglut2* and were probably serotonergic cells (also see “Monoamine neurons” section).

In the other midbrain nuclei, notable expression of *Ox1r* was found in the nucleus sagulum (SAG) and pedunculopontine tegmental nucleus (PPT). The SAG was a small cluster of *Vglut2*-positive cells, and almost all cells in the SAG showed low expression of *Ox1r*. In the PPT, most *Vglut2*-negative neurons, presumed to be cholinergic neurons, expressed *Ox1r* at the highest intensity in the brain. In contrast, *Ox2r*-positive cells in the PPT were *Vglut2*-positive. In the nucleus of the brachium of the inferior colliculus (NB) and parabigeminal nucleus (PBG), which are composed of *Vglut2*-positive neurons, cells expressing a high level of *Ox2r* were densely localized. Moderate expression of *Ox2r* was found in the superior colliculus (SC) and red nucleus (RN). In the motor-related SC (SCm), *Ox2r* expression was found in both *Vgat*-positive and *Vglut2-*positive cells, whereas *Ox2r* expression was found only in *Vgat*-positive cells in the sensory-related SC (SCs).

### 2.5 Pons

In the locus coeruleus (LC) and laterodorsal tegmental nucleus (LDT), there were numerous *Ox1r*-expressing cells with high *Ox1r* expression and no *Ox2r*-expressing cells. Nearly all *Ox1r*-positive cells in the LC and LDT were negative for both *Vgat2* and *Vgat*, which are assumed to be adrenergic and cholinergic neurons, respectively. In the sublaterodorsal tegmental area (SLD), a moderate number of *Vglut2*-positive cells were *Ox1r*- or *Ox2r-*positive, and a small number of *Vglut2*-positive cells expressed both *Ox1r* and *Ox2r*. In Barrington’s nucleus (BAR), which is composed of mostly *Vglut2*-positive neurons, *Ox1r*-positive cells, *Ox2r*-positive cells, and *Ox1r*- and *Ox2r*-double positive cells were observed (Fig. 4A-D). The pontine gray (PG) and tegmental reticular nucleus (TRN) were abundant in *Ox2r*-positive neurons, which were *Vglut2*-positive (Fig. 4 E-H). In addition, the PG and TRN also intensely expressed *Vglut1* (Fig. S2 Panel 125-146). The principal sensory nucleus of the trigeminal nerve (PSV) expressed only *Ox2r*. In the ventrolateral PSV (PSVvl), *Ox2r*-positive neurons were *Vglut2*-positive, while in the dorsomedial PSV (PSVdm), *Ox2r*-positive neurons were mainly *Vgat*-positive.

**Fig. 4.**
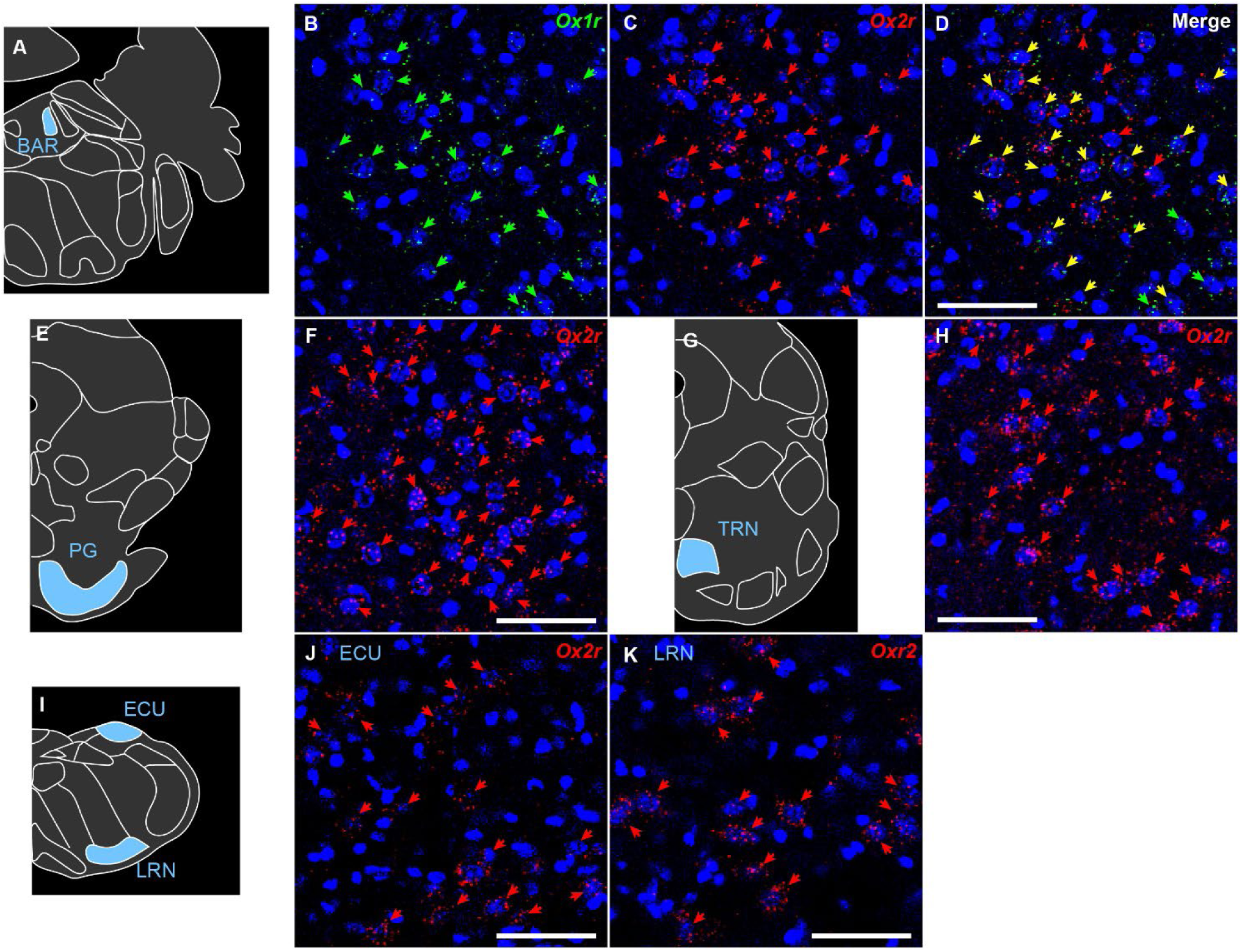
Representative expression of *Ox1r* and *Ox2r* in the brainstem. (A-D) *Ox1r* and *Ox2r* expression in Barrington’s nucleus. (E-F) *Ox2r* expression in the pontine gray. (G-H) *Ox2r* expression in the tegmental reticular nucleus. (I-K) *Ox2r* expression in the external cuneate nucleus (J) and the lateral reticular nucleus (K). Green, red, and yellow arrows denote *Ox1r-*, *Ox2r-,* and both receptor-expressing cells. Scale bars: 50 μm. All sections every 120 μm from single mouse were quantified.

### 2.6 Medulla and cerebellum

In the medulla, *Ox1r* expression was generally low and sparse. A relatively dense population of *Ox1r*-positive cells was observed in the dorsal motor nucleus of the vagus nerve (10N), compact part of the nucleus ambiguous (AMBc), medial vestibular nucleus (MVE), lateral part of the paragigantocellular reticular nucleus (PGRNl), nucleus raphe obscurus (RO), nucleus raphe pallidus (Rpa) and spinal vestibular nucleus (SPIV), although the expression per cell was low. In contrast, a dense population of *Ox2r*-expressing cells with a high expression level was observed in many nuclei in the medulla. In the cochlear nuclei, almost all neurons in the granular region (GRC) expressed *Ox2r*, while the ventral cochlear nucleus (VCO) did not express *Ox2r* and the dorsal cochlear nucleus (DCO) moderately expressed *Ox2r*. Cholinergic motor nuclei also showed moderate expression of *Ox2r* (Also see “Cholinergic neuron” section). Moderate *Ox2r* expression was found in *Vgat*-positive cells and *Vglut2*-positive cells in the cuneate nucleus (CU), intermediate reticular nucleus (IRN), magnocellular reticular nucleus (MARN), dorsal part of the medullary reticular nucleus (MdV), nucleus of the solitary tract (NTS), parvocellular reticular nucleus (PARN), dorsomedial part of the spinal nucleus of the trigeminal nerve (SPVd) and oral part of the SPV (SPVo). The external cuneate nucleus (ECU) contained numerous cells that were negative for both *Vgat* and *Vglut2* and highly expressed *Ox2r*. In the linear nucleus (LIN), lateral reticular nucleus (LRN), ventral part of the nucleus prepositus (PRPv), and nucleus X, a dense population of highly *Ox2r*-expressing cells was observed (Fig. 4I-K), and the *Ox2r*-expressing cells were *Vglut2*-positive (Fig. S2 panel 200-221).

### 2.7 Monoamine neurons

In serotonergic neurons, high expression levels of *Ox1r* and/or *Ox2r* were observed, but the proportions of *Ox1r* and *Ox2r* expression differed among cell groups (Fig S3). In the anterior DRN, referred to as the B7 group, where relatively large 5-HT-positive cells were distributed, numerous neurons expressed either *Ox1r* or *Ox2r*, or both (Fig. 5 A-D). The dominant cell types in the B7 group were *Ox1r*-positive (*Ox1r*: 36.6%; *Ox2r*: 18.8%; *Ox1r* + *Ox2r*: 22.6%, Fig. 5K). In the posterior DRN, referred to as B4/6 groups, where relatively small cells were clustered in the midline, the expression level per cell was weaker than in other serotonergic nuclei, and the proportion of orexin receptor-expressing cells was the smallest among serotonergic cell groups (*Ox1r*: 11.0%; *Ox2r*: 20.1%; *Ox1r* + *Ox2r*: 9.1%, Fig. 5E-H, K). In the MRN, called B5/8/9 cell groups, there was a large number of *Ox1r*-positive neurons, whereas the number of *Ox2r*-positive neurons was relatively small (*Ox1r*: 45.8%; *Ox2r*: 6.3%; *Ox1r* + *Ox2r*: 10.4%, Fig. 5K). The ventral raphe nucleus (VRN), called B1/2/3 cell groups, was a relatively small cluster of serotonergic neurons located in the ventral medulla, and the majority of receptor-expressing cells was *Ox1r*-positive (*Ox1r*: 62.5%; *Ox2r*: 1.3%; *Ox1r* + *Ox2r*: 2.5%, Fig. 5I-K).

**Fig. 5.**
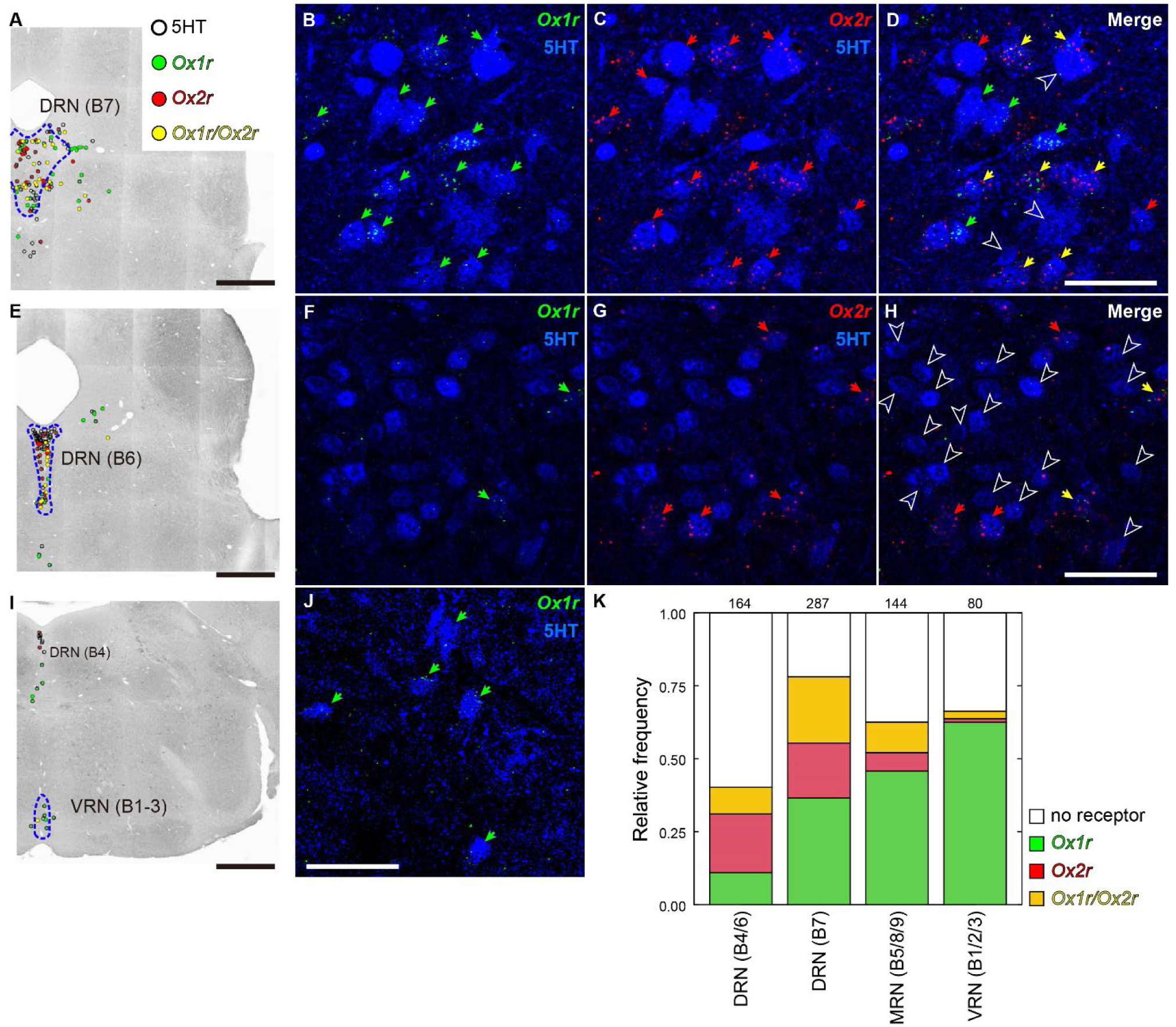
Orexin receptor expression of serotonergic neurons in the brainstem. (A, E, I) Reconstructed image showing *Ox1r-* and *Ox2r-*positive serotonergic neurons. Open circle, green circle, red circle, and yellow circle denote no receptor, *Ox1r-, Ox2r-,* and both receptor-positive cells, respectively. Scale bars: 500 μm. (B-D) *Ox1r-* and *Ox2r-*expression and immunofluorescence of 5HT in the B7 region of the dorsal raphe nucleus. (F-H) Immunofluorescence of 5HT in the B6 region of the dorsal raphe nucleus. (J) *Ox1r-*expression and immunofluorescence of 5HT in the ventral raphe nucleus. 5HT immunofluorescence is shown in blue. Green, red, and yellow arrows denote *Ox1r-*, *Ox2r-,* and both receptor-expressing cells, respectively. Outlined arrowheads denote the no-receptor expressing 5HT-ir neurons. Scale bars: 50 μm. (K) Regional differences in the expression of receptor subtypes in serotonergic neurons. The numbers above the bar plot indicate the total number of cells. All sections every 120 μm from single mouse were quantified.

Dopaminergic cell groups A11-15, in addition to periventricular nuclei, PVD and PVH, were distributed in the hypothalamus. In these cell groups except A11, a small proportion of dopaminergic cells were *Ox2r*-positive, and very few cells were *Ox1r*-positive (*Ox1r*: 1.0%, *Ox2r*: 8.3% in total, Fig. S4). A11 dopaminergic cells were scattered around the dorsal border of the posterior hypothalamus, and approximately half of these cells were *Ox2r*-positive (51.7%, Fig. 6A,B). In contrast to the hypothalamic dopaminergic neurons, midbrain dopaminergic neurons called A8-10 groups contained a relatively high proportion of *Ox1r*-positive cells. Although the distributions of *Ox1r*- or *Ox2r-*positive cells were not uniform within each nucleus (Fig. S4), similar proportions of *Ox1r*-positive cells were found in A8 RR, A9 SNc, and A10 VTA neurons (RR: 13.8%, SNc, 14.8%, VTA: 16.6%, Fig. 6C, D and G). Dopaminergic neurons in the midbrain were also distributed in the DRN, anterior to the B7 serotonergic cell clusters. In these DRN dopaminergic cells, receptor-positive neurons were never found.

**Fig. 6.**
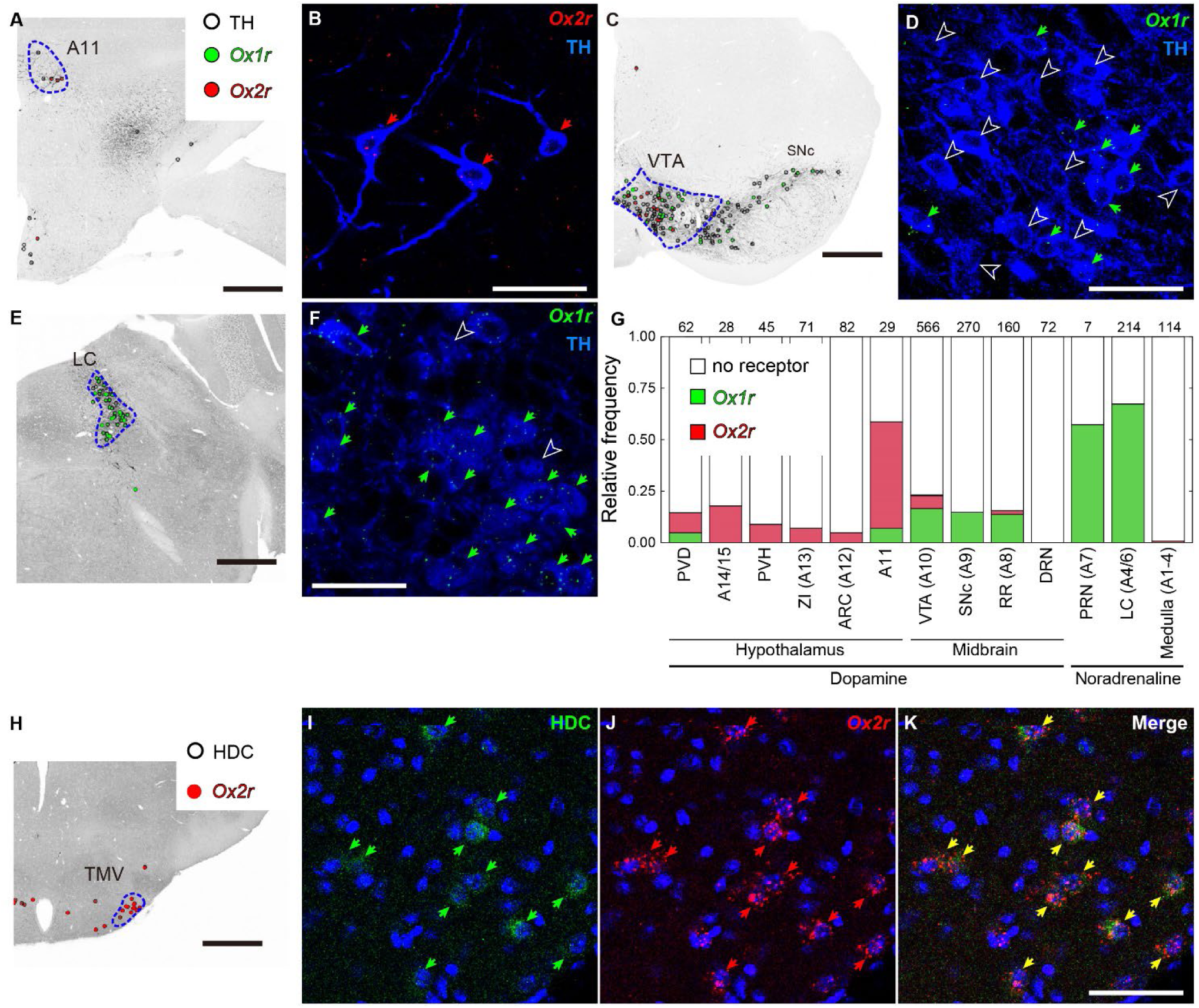
Orexin receptor expression of dopaminergic, noradrenergic, and histaminergic neurons. (A, C, E) Reconstructed image showing *Ox1r*- and *Ox2r-*positive cells in TH-positive neurons. Open circle, green circle, and red circle denote no receptor, *Ox1r,* and *Ox2r* positive cells, respectively. Scale bars: 500 μm. (B) *Ox2r-*expression and immunofluorescence of TH in the A11 region of the hypothalamus. (D) *Ox1r*-expression and immunofluorescence of TH in the ventral tegmental area. (F) *Ox1r*-expression and immunofluorescence of TH in the locus coeruleus. TH immunofluorescence is shown in blue. The green and red arrows denote the *Ox1r-* and *Ox2r-*expressing cells, respectively. Outlined arrowheads denote the no-receptor expressing TH-ir neurons. Scale bars: 50 μm. (G) Regional differences in the expression of receptor subtypes in TH-ir neurons. The numbers above the bar plot indicate the total number of cells. (H) Reconstructed image showing *Ox2r-*positive histaminergic neurons. A total of 97.4% of histidine decarboxylase (HDC)-ir neurons were positive for *Ox2r*. Scale bar: 500 μm. (I) Immunofluorescence of histidine decarboxylase (green). (J) mRNA of *Ox2r* (red). (K) Merged image of (I) and (J). Cell nuclei are shown in blue. Green, red, and yellow arrows denote the histidine decarboxylase-, *Ox2r-* and both expressing cells, respectively. Scale bar: 50 μm. All sections every 120 μm from single mouse were quantified.

Noradrenergic neurons in the pons, called A4/A6 (LC) and A7 groups, showed strong and dense expression of *Ox1r* (Fig. 6E,F) but no expression of *Ox2r* (LC *Ox1r*: 67.3%, A7 *Ox1r*: 57.1%). These features were well contrasted with the lack of expression of receptors of TH-ir neurons in the medulla, which are presumed to be adrenergic and/or noradrenergic neurons (Fig. 6G, S4).

Histaminergic neurons showed a scattered distribution throughout the hypothalamus but clustered in the TMV (Fig. 6H). Strong expression of *Ox2r* was observed in almost all histaminergic neurons (97.3%, Fig. 6H-K, S5), while no expression of *Ox1r* was found.

### 2.8 Cholinergic neurons

In the cerebral nuclei, a moderate proportion of cholinergic neurons expressed either *Ox1r* or *Ox2r* (Fig. 7). A total of 15.5% of *Chat*-positive neurons in the MS and NDB were *Ox1r*-positive, whereas 3.4% of those were *Ox2r*-positive. In the Cpu, there were scattered large *Chat*-positive neurons, approximately one-third of which were *Ox2r*-positive (34.9%). Although *Chat*-positive neurons were also found in layers II/III of the isocortex and MHb, almost all of them were receptor-negative (layers II/III: 99.3%, MHb: 99.7%).

**Fig. 7.**
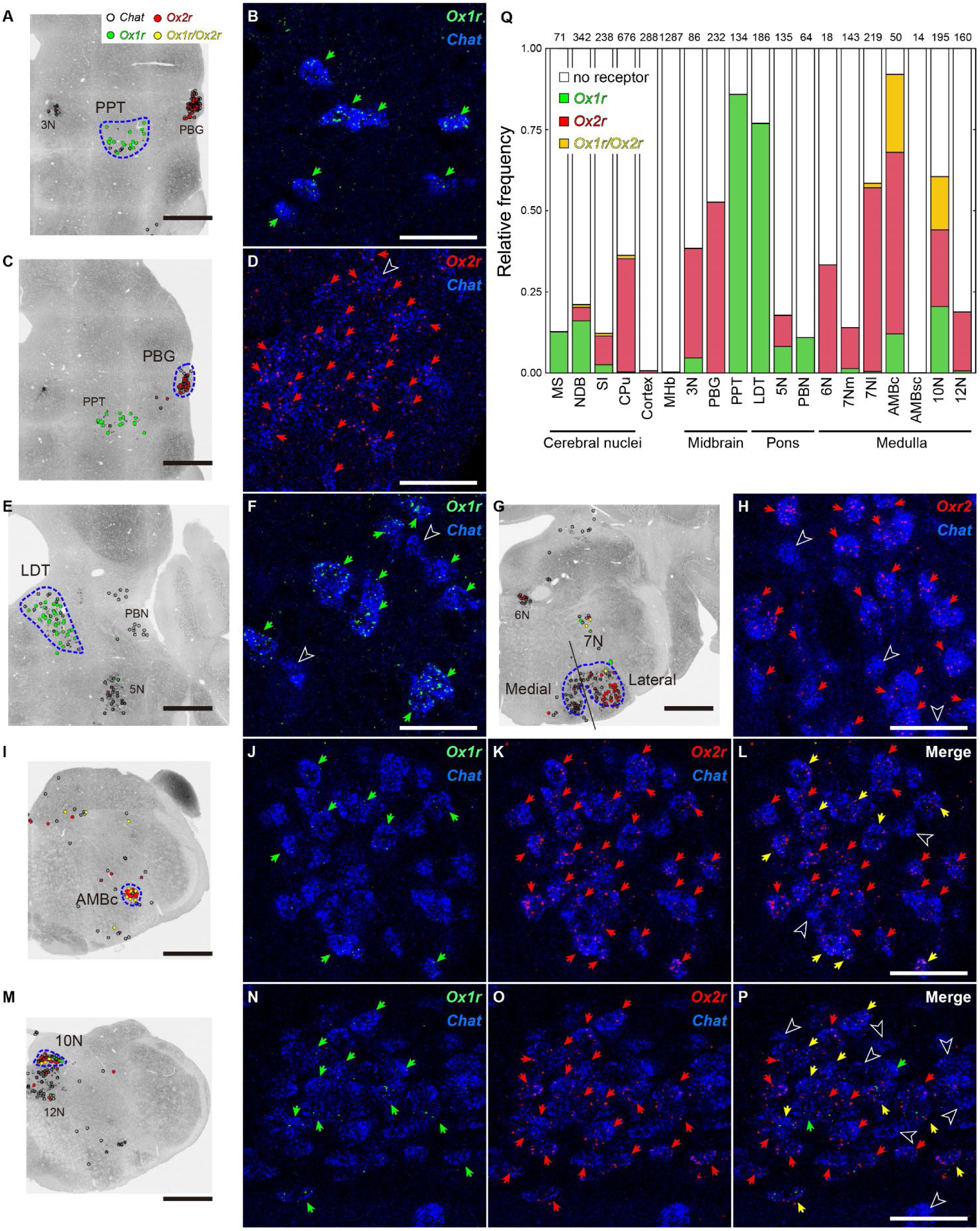
Orexin receptor expression in cholinergic neurons. (A, C, E, G, I, M) Reconstructed image showing *Ox1r*- and *Ox2r*-positive cholinergic neurons. Open circle, green circle, red circle, and yellow circle denote no receptor, *Ox1r-*, *Ox2r-,* and both receptor-positive cells, respectively. Scale bars: 500 μm. (B) *Ox1r* and *Chat* expression in the pedunculopontine nucleus. (D) *Ox2r* and *Chat* expression in the parabigeminal nucleus. (F) *Ox1r* and *Cha*t expression in the laterodorsal tegmental nucleus. (H) *Ox2r* and *Chat* expression in the lateral part of the facial motor nucleus (7N). (J-L) *Ox1r*-, *Ox2r*- and *Chat* expression in the ambiguus nucleus, compact. (N-P) *Ox1r-*, *Ox2r-* and *Chat* expression in the dorsal motor nucleus of the vagus nerve (10N). *Chat* mRNA is shown in blue. Green, red, and yellow arrows denote *Ox1r*-, *Ox2r*- and both receptor-expressing cells, respectively. Outlined arrowheads denote the no-receptor expressing *Chat-*positive neurons. Scale bars: 50 μm. (Q) Regional differences in the expression of receptor subtypes in cholinergic neurons. The numbers above the bar plot indicate the total number of cells. All sections every 120 μm from single mouse were quantified.

In the brainstem, very high expression of *Ox1r* and/or *Ox2r* was found. The 3N has been known to be a small cluster of *Chat*-positive neurons, and *Ox2r*-positive neurons were dominant in the 3N (*Ox1r*: 4.7%, *Ox2r*: 33.7%). In the PBG, there was a dense cluster of small neurons that expressed both *Chat* and *Vglut2* mRNAs. These neurons also expressed *Ox2r* in a high proportion (52.6%), but the expression level per cell was relatively low (Fig. 7C,D). In contrast to these *Ox2r*-positive nuclei in the midbrain, almost all *Chat*-positive neurons in the PPT showed highly intense expression of *Ox1r* (Fig. 7A,B). Similar to PPT *Chat* neurons, *Chat*-positive neurons in the LDT also show strong expression of *Ox1r* with a high proportion (PPT: 85.8%, LDT: 76.9%, Fig. 7E,F). The 5N and PBN in the pons showed a small percentage of receptor-positive cells (*Ox1r*: 8.1%, *Ox2r*: 9.6% in 5N, *Ox1r*: 10.9% in PBN, Fig. S6). In the 6N and 7N, there was dominant expression of *Ox2r* with high intensity, and a subregional difference in *Ox2r* expression was found between the medial and lateral 7N (*Ox2r* in 6N: 33.3%, 7N medial: 12.6%, 7N lateral: 56.6%, Fig. 7G.H).

Subregional differences in receptor expression were also found in the AMB. The AMBc was a relatively large cluster of *Chat*-positive cells located in the anterior portion of the AMB, and almost all neurons expressed *Ox1r* and/or *Ox2r* (*Ox1r:* 12.0%, *Ox2r*: 56.0%, *Ox1r,* and *Ox2r*: 24.0%, Fig 7I-L). In contrast, the *Chat*-positive neurons in the AMBsc, the posterior small cell cluster of AMB, showed no expression of the receptor mRNAs. Similar to AMBc, the proportion of *Ox1r*- and *Ox2r*-double-positive cells was high in the 10N (*Ox1r:* 20.5%, *Ox2r*: 23.6%, *Ox1r* and *Ox2r*: 16.4%, Fig 7M-P). The expression profile in the 12N was different from that in the 10N, and 18.1% of the *Chat*-positive neurons in the 12N expressed *Ox2r*.

## 3 Discussion

In the present study, the bHCR system was used to visualized *Ox1r* and *Ox2r* mRNA, combined with other neuron type markers, which provides comprehensive information on the distribution of orexin receptor mRNA and neuron types expressing orexin receptors in mouse brains. We showed that many brain regions express both *Ox1r* and *Ox2r* mRNAs, but only a few cells express both *Ox1r* and *Ox2r* mRNAs. Double receptor-positive cells are observed in certain brain regions, such as the DRN, LM, BAR AMBc, VMH, and 10N. Since orexin receptors are promiscuous GPCRs (16), how downstream signaling of double receptor-positive cells differs from that of single receptor-positive cells needs to be determined individually. The subcellular localization of OX1R and OX2R may be different.

We also showed that *Ox1r*-expressing cells and *Ox2r*-expressing cells in the same region tend to be classified into different subpopulations, such as glutamatergic and GABAergic neurons. For example, *Ox1r* and *Ox2r* were expressed predominantly in *Vgat*-negative cells and *Vgat*-positive cells, respectively, in isocortex layer II/III, olfactory areas, and the cortical subplate in the cerebrum, as shown in the BLA (28). In the cerebrum, although *Vglut2* was not expressed in almost all of the excitatory neurons, almost all of the *Vgat*-negative neurons expressed *Vglut1* (Fig. S2). Therefore, we assumed that *Vgat*-negative neurons were excitatory neurons in the cerebrum.

In the thalamus and more caudal brain regions, the majority of excitatory neurons express *Vglut2*. *Vglut1* is also expressed in certain neural groups along with *Vglut2*, such as PG, TRN, PRPv, nucleus X, LRN, and LIN. The ECU and GRC express only *Vglut1*. Interestingly, these *Vglut1*-positive regions strongly expressed *Ox2r* and are *Barhl1-*lineage cells (34). *Barhl1* is a homeobox transcription factor and is expressed in a subset of developing and mature glutamatergic cells in the hindbrain and the cerebellum. Exceptionally, the VCO, which expresses *Vglut1* but is not the *Barhl1*-lineage (34), did not express *Ox2r*. These findings imply that *Barhl1*-lineage cells constitute a certain subpopulation of hindbrain excitatory neurons that are characterized by *Vglut1* and strong *Ox2r* expression.

Regarding the wake-promoting action of the orexin system (2), each orexin receptor exhibits a distinct distribution in many wake-active or wake-promoting neurons. *Ox1r* is abundantly expressed in the DRN, LC, PPT, and LDT, mildly expressed in the NDB, VTA, and parabrachial nucleus, and not expressed in the histaminergic neurons of the TMV. *Ox2r* is abundantly expressed in the NDB, histaminergic TMV neurons, DRN, and PPT, mildly expressed in the VTA, and parabrachial nucleus, and not expressed in the LC or LDT. These observations are generally consistent with those of previous studies (14, 25–28, 35, 36). Furthermore, the cellular resolution analysis showed that the DRN expresses both *Ox1r* and *Ox2r* but serotonergic cells expressing both receptors are a minor population of DRN neurons. Consistently, single-cell RNA-seq of Pet1-lineage DR neurons showed that cell clusters expressing *Ox1r* are largely different from cell clusters expressing *Ox2r* (37). *Ox1r*-expressing clusters express Gad2, whereas *Ox2r*-expressing clusters express *Vglut3*. In addition, orexin neurons themselves rarely expressed orexin receptors, which suggests that orexin neurons do not augment or suppress their activity via autoreceptor feedback.

The expression of the orexin receptor in wake-promoting neurons supports the role of orexins in wakefulness but orexin receptors are also expressed in NREM sleep-promoting neurons (2). *Ox2r* is abundantly expressed in the MPOA, including the ventrolateral POA, and in the ventrolateral periaqueductal gray (vlPAG), subthalamic nucleus, and nucleus of the solitary tract, caudoputamen, and nucleus accumbens. Given that orexin neurons are generally inactive during NREM sleep (38, 39), orexin receptor signaling in these sleep-promoting neurons may be involved more in other behaviors during wakefulness than in sleep regulation.

Glutamatergic neurons in the SLD and cholinergic neurons in the PPT and LDT play an important role in the regulation of REM sleep (2). Glutamatergic neurons in the SLD expressed *Ox1r* and/or *Ox2r*, consistent with a previous study (19). The orexinergic projection to SLD glutamatergic neurons stabilized REM sleep, and activation of these neurons promoted a REM sleep-related symptom, cataplexy, in the orexin-deficient mice (19, 40). In the PPT and LDT, a vast majority of cholinergic neurons only express *Ox1r*. Whereas the LDT did not express *Ox2r*, the PPT contained abundant *Ox2r*-expressing cells that were glutamatergic, not cholinergic. Thus, the expression of the orexin receptor differs depending on brain region and cell type.

Regarding the metabolic effects of the orexin system, orexin receptors are abundantly expressed in brain regions regulating energy metabolism, such as the medullary raphe, DMH, and ARC. More than 50% of serotonergic neurons in the medullary raphe, including the raphe pallidus nucleus (B1), raphe obscurus nucleus (B2), and raphe magnus nucleus (B3), express *Ox1r* but not *Ox2r*, which serve as sympathetic premotor neurons to enhance thermogenesis by brown adipose tissue (BAT)(41, 42). The DMH is one of the major upstream regions of medullary raphe neurons, and the excitatory projection from the DMH to the medullary raphe enhances sympathetic outflows. However, *Ox2r*-expressing DMH cells were mainly GABAergic. The role of DMH GABAergic neurons in the regulation of sympathetic and other functions remains unclear. In the ARC, GABAergic neurons express *Ox2r* but not *Ox1r*. Orexin activates ARC GABAergic neurons (43) and suppresses POMC neurons, a subpopulation of ARC GABAergic neurons (44). Since the loss of either *Ox1r* or *Ox2r* only partially mimics the obesity-prone phenotype of orexin-deficient mice (13), it is suggested that both *Ox1r* and *Ox2r* may function independently to render mice less prone to obesity.

The orexin system is also involved in reward behavior and addiction (7, 45). Pharmacological studies using orexin and orexin receptor antagonists have repeatedly reported the role of OX1R in the VTA in reward-related behavior (46–50). However, the expression level of *Ox1r* per cell was very low, and less than 20% of dopaminergic cells expressed *Ox1r*. It may be possible that orexin receptors are functional even if the number of mRNA copies per cell is very low (51). In addition, orexin may change the presynaptic function of VTA dopaminergic cells (52). The nucleus accumbens contained a dense population of *Ox2r*-expressing cells. Orexin modulates risk-avoidance and reward behaviors through nucleus accumbens neurons (53, 54). OX2R in the nucleus accumbens modulates anxiety (55) and nociception (56).

High expression of *Ox2r* with variable *Ox1r* expression is observed in cholinergic motor neurons in the brain stem, such as the lateral facial nucleus (7N) that regulates muscles for facial expression, and the ambiguus nucleus compact part (AMBc) that innervates the pharyngeal constrictor and the cervical esophageal muscles (57–59). In contrast, the medial 7N and ambiguus nucleus semicompact part (AMBsc), which are not associated with motor regulation (57, 58, 60, 61), did not show *Ox2r* expression. *Ox1r* and *Ox2r* are expressed in cholinergic neurons of the dorsal motor nucleus of the vagus nerve (10N), which serves parasympathetic functions in many thoracic and abdominal organs. These findings suggest that orexin receptor signaling, mainly via OX2R, works to enhance motor action and parasympathetic activity.

In this study, we visualized orexin receptor mRNA in the brains of male mice. However, female mice could exhibit a different distribution of orexin receptors than males, especially in the hypothalamus. In female rats, *Ox1r* expression in the hypothalamus changed with the estrous cycle, but *Ox2r* expression was stable (62). Consistently, the regions expressing sex steroid hormone receptors, such as the MPOA, BNSTpr, VMH, MeA, and PAG, also abundantly expressed *Ox1r*. Sexual differences in orexin receptor expression will be the focus of future studies.

The bHCR system allows us to detect target molecules with much higher sensitivity (30, 63–66), and this is the first report to apply branched HCR to the *in situ* detection of multiple targets. The time required for obtaining confocal images was reduced due to the ability to perform highly sensitive detection by branched *in situ* HCR, allowing us to inspect many brain sections within a shorter time. Split-initiator probes also suppressed noise signals in the branched *in situ* HCR, as confirmed by staining sections of *Ox1r* and *Ox2r*-deficient mice. Very few cells expressing both receptors indicate that the signals detected in this study are highly specific.

In summary, the detailed distribution of orexin receptors helps interpret the consequences of optogenetic manipulation of orexin circuits and provides valuable insight into the development of orexin receptor agonists and antagonists for human sleep and related diseases.

## 4 Materials and methods

### 4.1 Animals

All animal procedures were conducted in accordance with the Guidelines for Animal Experiments of Toho University and were approved by the Institutional Animal Care and Use Committee of Toho University (Approval Protocol ID #21-53-405). Breeding pairs of C57BL/6J mice were obtained from Japan SLC and CLEA Japan. Mice were raised in our breeding colony under controlled conditions (12 h light/dark cycle, lights on at 8:00 A.M., 23 ± 2 °C, 55 ± 5% humidity, and ad libitum access to water and food). We also used male *Ox1r*-deficient mice (13) and male *Ox2r*-deficient mice (12).

Twelve-to 15-week-old male mice were anesthetized with sodium pentobarbital (50 mg/kg, i.p.) and then transcardially perfused with 4% paraformaldehyde (PFA) in phosphate-buffered saline (PBS) two-four hours after the start of the light phase. The brains were postfixed in 4% PFA at 4 °C overnight, followed by cryoprotection in 30% sucrose in PBS for two days, embedded in Surgipath (FSC22, Leica Biosystems), and stored at -80 °C. The frozen brains except olfactory bulbs were cryosectioned coronally at a thickness of 40 μm. The sections were stored in antifreeze solution (0.05 M phosphate buffer, 30% glycerol, 30% ethylene glycol) at -25 °C until use. Every third section from the serial sections was processed for the same staining procedures. To confirm the reproducibility of the staining, sections from two or three mice were used for each staining combination.

### 4.2 Branched HCR with IHC

#### 4.2.1 Preparation of probes and hairpin DNAs

The probes for ISH-HCR were designed to minimize off-target complementarity using a homology search by NCBI Blastn (https://blast.ncbi.nlm.nih.gov), and they were designed to have split-initiator sequences with an mRNA binding site. For *Ox1r* probes, five target sequences overlapped with exon 5-exon 6, which are absent in *Ox1r* KO mice. For *Ox2r* probes, 10 target sequences were selected from the complete cDNA sequence because mRNA expression was completely abolished in *Ox2r* KO mice. Other probes were designed according to our published protocol (Supplementary Table S3). The DNA probes were synthesized as standard desalted oligos (Integrated DNA Technologies) and purified by denaturing polyacrylamide gel electrophoresis (PAGE) using 20% polyacrylamide gels. The fluorescent hairpin DNAs were prepared by conjugation of succinimidyl ester of fluorophores (Sarafluore488, ATTO550, or ATTO647N) and synthesized with a C12 amino-linker at the 5’ end (Integrated DNA Technologies or Tsukuba Oligo Services). The DNA probes and short hairpin DNAs were purified by PAGE using 20% polyacrylamide gels. The double-stem hairpin DNAs were designed as a combination of our established hairpin DNA sets according to a previous study for branched HCR(30). The double-stem hairpins were purified by PAGE using 15% polyacrylamide gels. The detailed procedures for DNA purification were described in our previous study (29).

#### 4.2.2 Probe hybridization

ISH procedures were similar to those previously described with some modifications (29, 67) The free-floating sections were washed with PBS containing 0.2% Triton (PBST) and immersed in methanol for 10 min, followed by PBST washing for 5 min twice. After washing, the sections were prehybridized for 5 min at 37 °C in a hybridization buffer containing 10% dextran sulfate, 0.5× SSC, 0.1% Tween 20, 50 μg/ml heparin, and 1× Denhardt’s solution. The sections were treated with another hybridization solution containing a mixture of 10 nM probes and incubated overnight at 37 °C. One to three probe sets were selected from the probe sets for *Ox1r*, *Ox2r*, *Vgat*, *Vglut1*, *Vglut2,* and *Chat* mRNAs. The control experiment was performed without probes or using sections of *Ox1r-* or *Ox2r*-deficient mice. After hybridization, the sections were washed three times for 10 min in 0.5× SSC containing 0.1% Tween 20 at 37 °C. Then, the sections were bleached by an LED illuminator for 60 min in PBST to quench autofluorescence (68).

#### 4.2.3 Branched HCR amplification

For 1st HCR amplification, the DH and H0 hairpin DNAs were used to form a DNA trunk (Fig. S1, Supplementary Table S4). These hairpin DNA solutions were separately snap-cooled (heated to 95 °C for 1 min and then gradually cooled to 65 °C for 15 min and 25 °C for 40 min) before use. The sections were incubated in amplification buffer (10% dextran sulfate in 8× SSC, 0.2% Triton X-100, 100 mM MgCl_2_) for 5 min and then immersed in another amplification buffer containing 150 nM DH and 60 nM H0 hairpin DNAs for 45 min at 25 °C. The samples were washed three times with PBST at 37 °C for 15 min followed by 2nd HCR amplification. The sections were incubated in the amplification buffer described above for 5 min and then immersed in another amplification buffer containing 60 nM snap-cooled fluorescent hairpin DNA and 120 nM assist oligo DNA for 2 hours at 25 °C.

In the case of combined ISH and immunohistochemistry, the sections were blocked using 0.8% Block Ace/PBST (Dainihon-Seiyaku), which was followed by overnight incubation with one or two antibodies from rabbit anti-TH, mouse anti-Calbindin D28k, guinea pig HDC, goat 5-HT, and rabbit orexin antibodies in 0.4% Block Ace/PBST at 4 °C (Supplementary Table S5. After washing three times with PBST, sections were incubated with a secondary antibody mixture with Hoechst 33342 (1 μg/ml) for an hour at room temperature. The sections were mounted on the slide glass and coverslipped with mounting media containing antifade (1% n-propyl gallate and 10% Mowiol4-88 in PBS).

### 4.3 Histological analysis

Fluorescence photomicrographs were obtained using a Nikon Eclipse Ni microscope equipped with the A1R confocal detection system under 20×/0.75 NA objective lenses at 4 μs/pixel speed at 0.628 μm/pixel resolution using autofocus (Nikon Instruments Inc., Tokyo, Japan). Tiled images were captured and automatically stitched by NIS-Elements C software (Nikon Instruments Inc., Tokyo, Japan).

Images were analyzed using ImageJ software (version 1.50i, NIH, USA). Quantification of the fluorescent photographs was performed at the same threshold and adjustment of contrast. All of the mapping procedures of receptor-expressing cells were performed manually under blinded conditions. If more than one granule contacted each cell nucleus, the cell was judged as mRNA-positive. If more than five puncta were observed, the expression level of each cell was regarded as high expression. The distributions of mRNA-positive cells were reconstructed by drawing circles on the original photographs. The 3-dimensional images of receptor-expressing cells were reconstructed from serial sections of which positions were adjusted manually according to the published brain atlas. The area location was basically decided according to the brain atlases (69, 70), while the area boundary shown in Fig. S2 was adjusted from the expression of *Vgat*, *Vglut1*, *Vglut2,* and *Chat* mRNAs, Calbindin and TH immunoreactivity. In each brain region, whether orexin receptor-positive cells were glutamatergic or GABAergic was examined using bHCR sections for *Ox1r*, *Vgat,* and *Vglut2*, and bHCR sections for *Ox2r*, *Vgat,* and *Vglut2*. For monoaminergic neurons, orexin receptor expression was examined using bHCR sections for *Ox1r* and *Ox2r* with immunostaining of anti-TH, HDC, or 5-HT antibodies. For cholinergic neurons, orexin receptor expression was examined using bHCR sections for *Ox1r*, *Ox2r,* and *Chat*.

## 5 Data, Materials, and Software Availability

Data are included in the main and supplementary figures. All other data supporting this study are available from the corresponding authors without restriction.

## 6 Conflict of interest

The authors declare that the research was conducted in the absence of any commercial or financial relationships that could be construed as a potential conflict of interest.

## 7 Author Contributions

YT: conceptualization, methodology, and formal analysis. YT and HF: writing, funding, acquisition, and supervision.

## 8 Acknowledgements

We thank N. Ohno, H. Arai and A. Iijima for animal care. This work was supported by the Precise Measurement Technology Promotion Foundation (to Y. T.), JSPS Kakenhi (21K06414 to Y. T.), Research Grant from Uehara Memorial Foundation (to H. F.), and Toho University Grant for Research Initiative Program (TUGRIP).

## 11 Supplementary tables

**Table S1.**
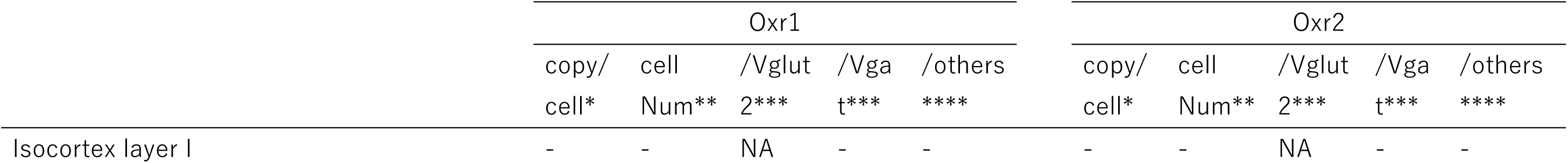
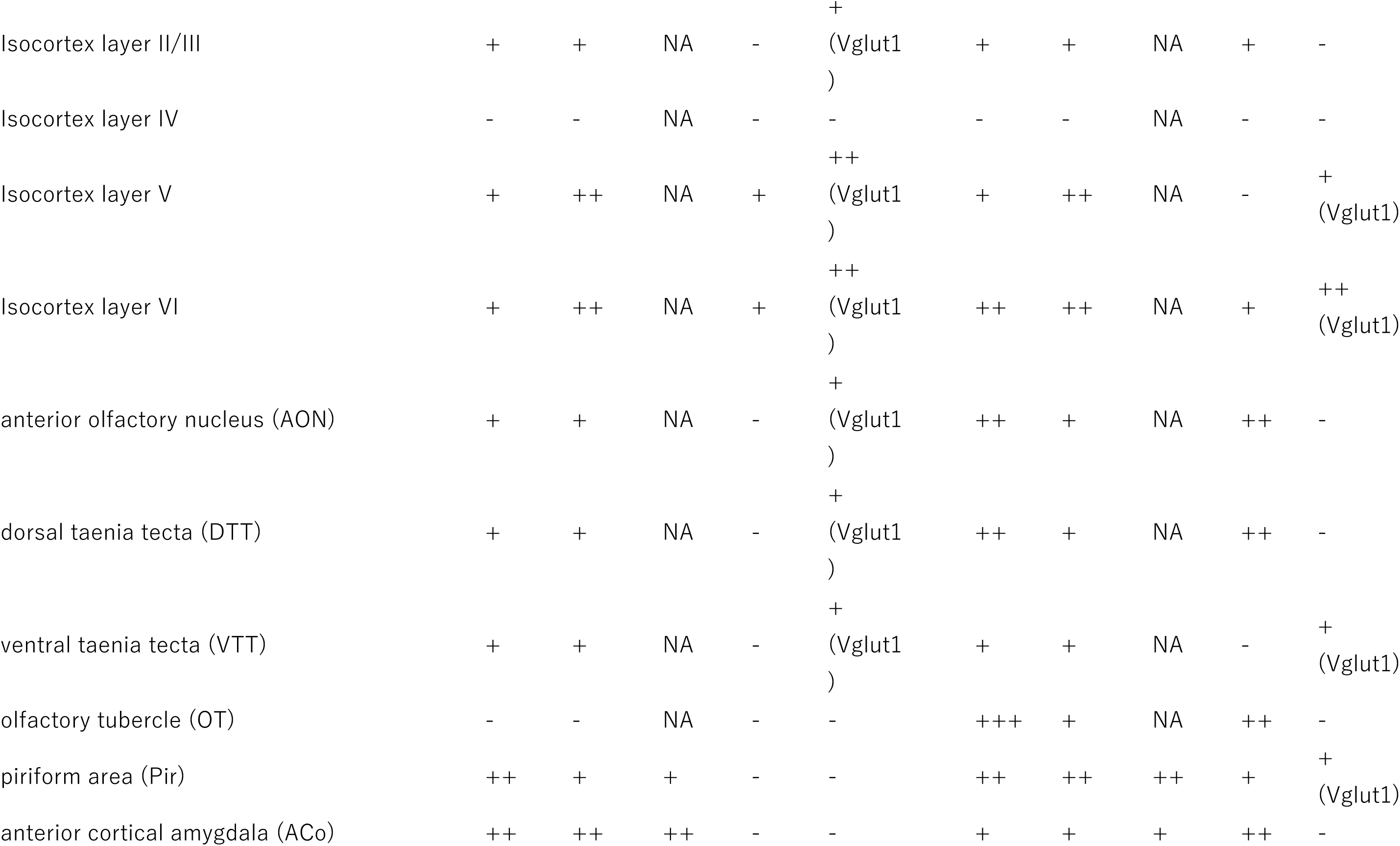
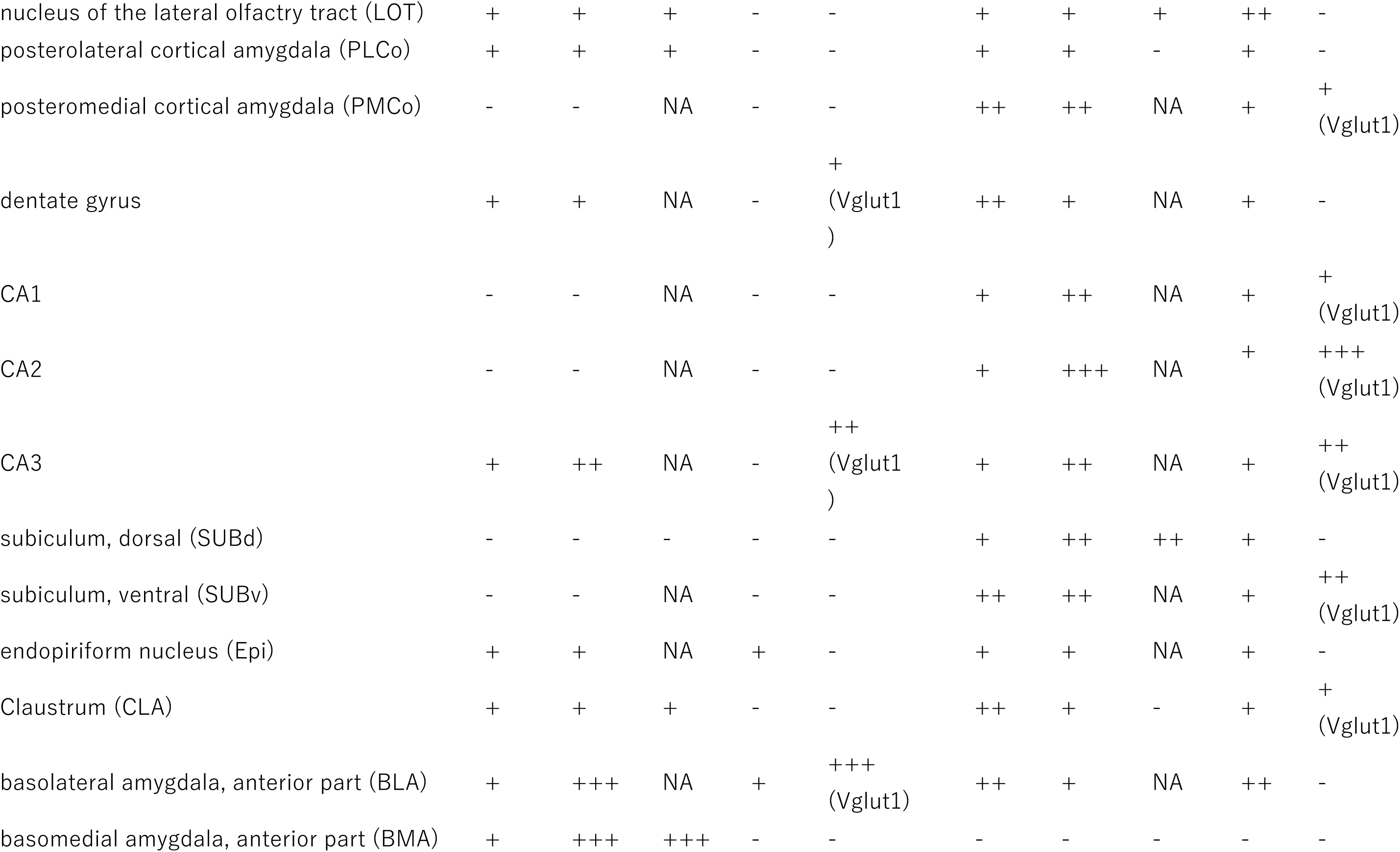
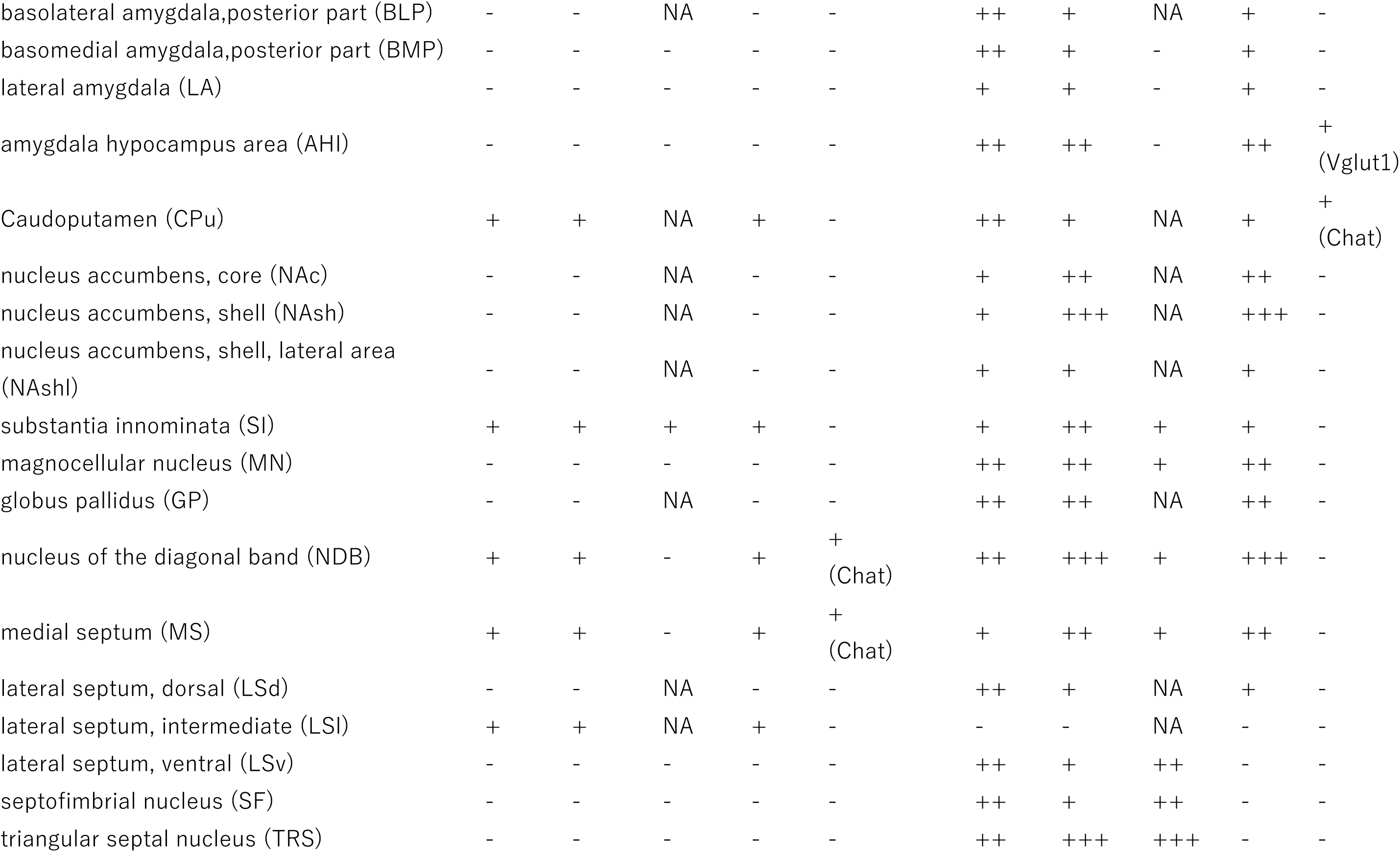
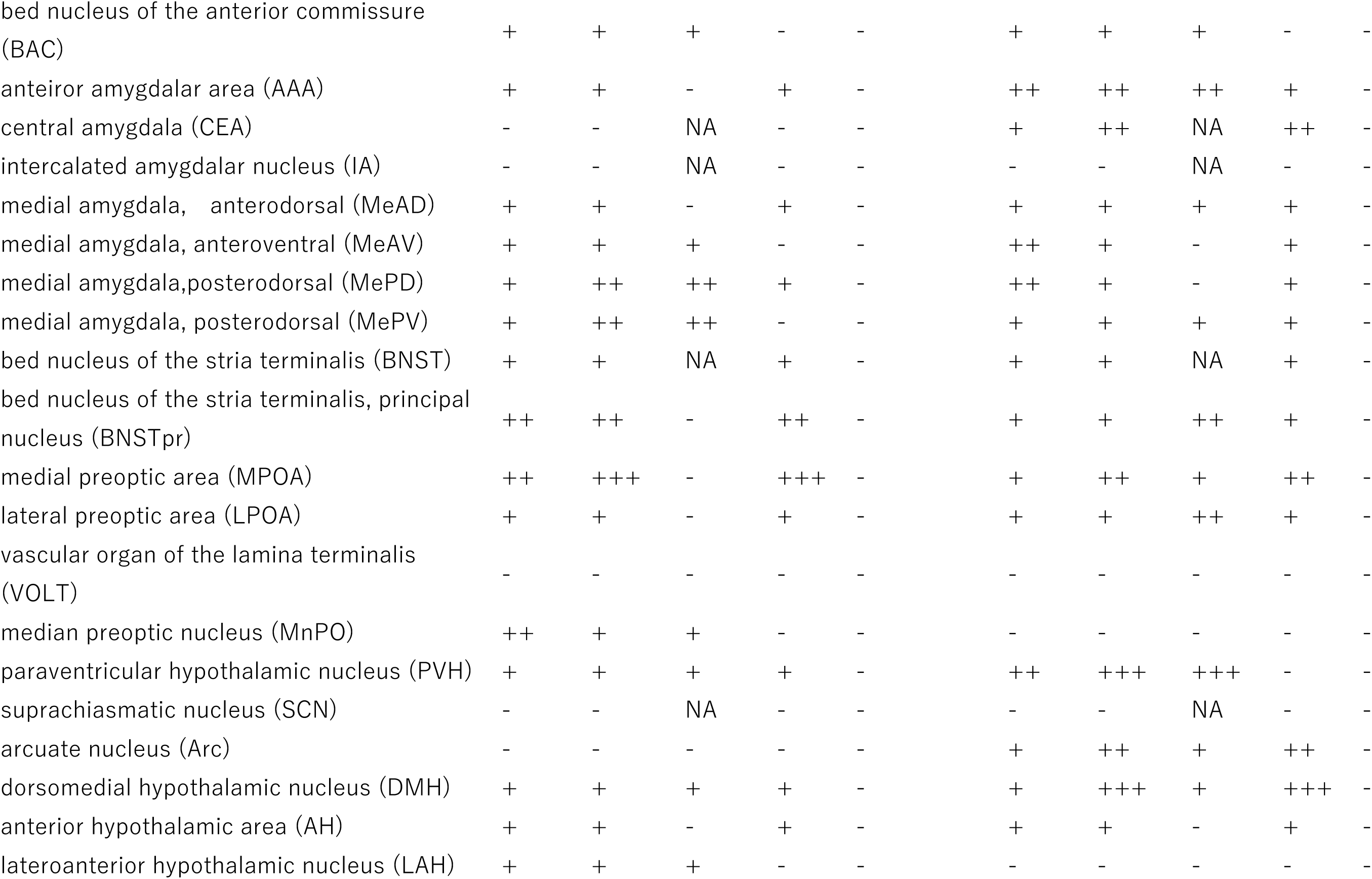
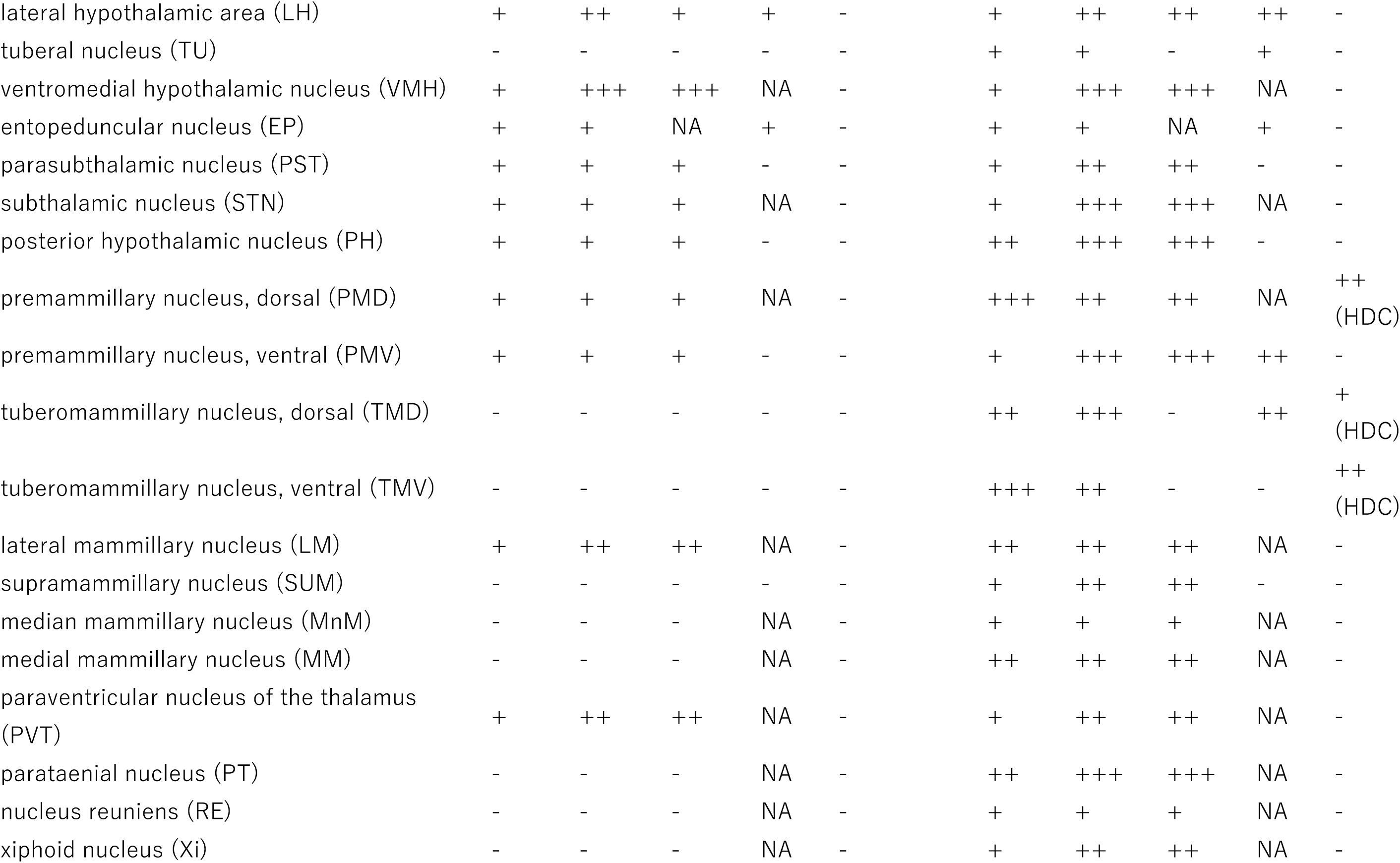
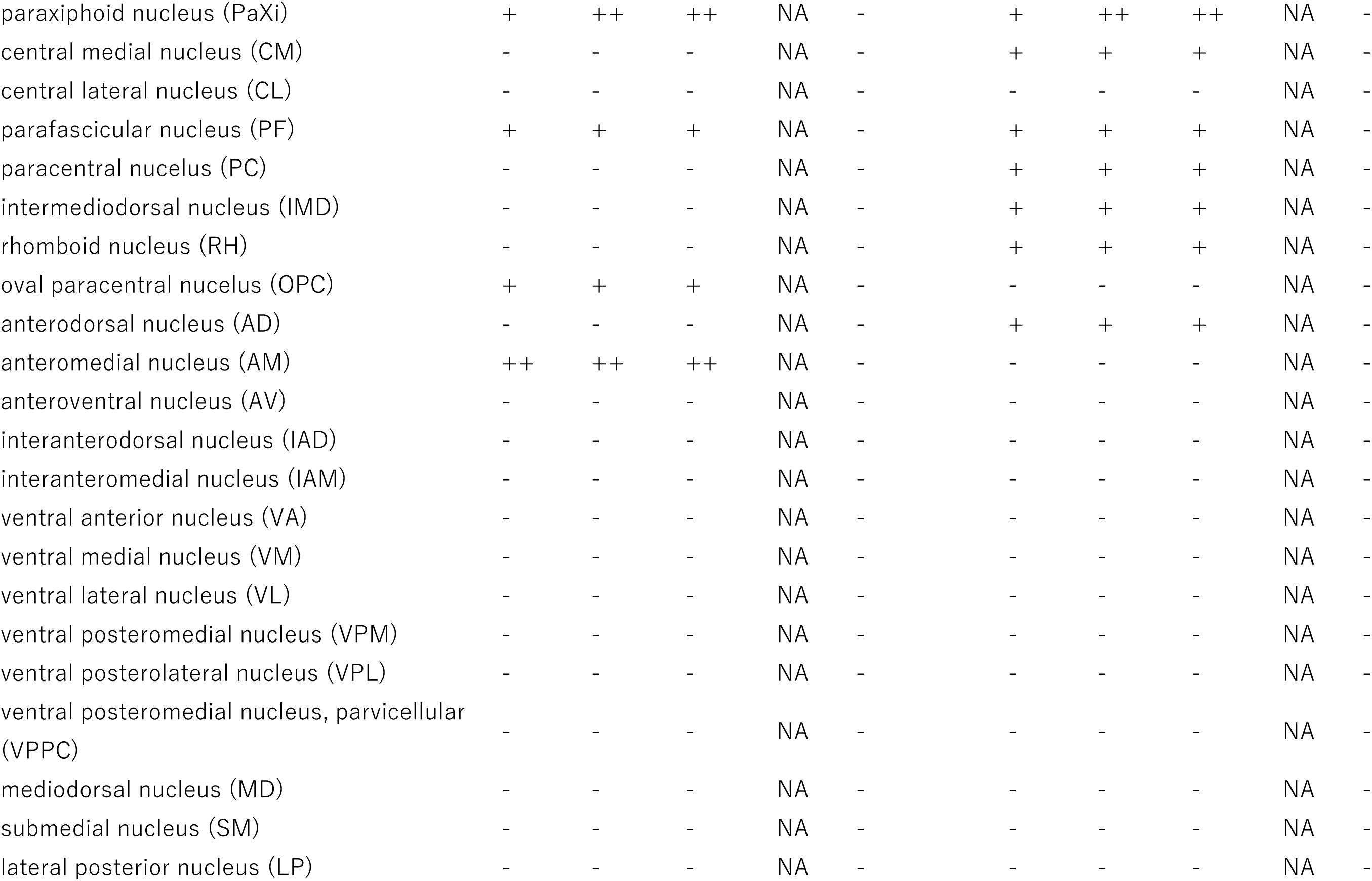
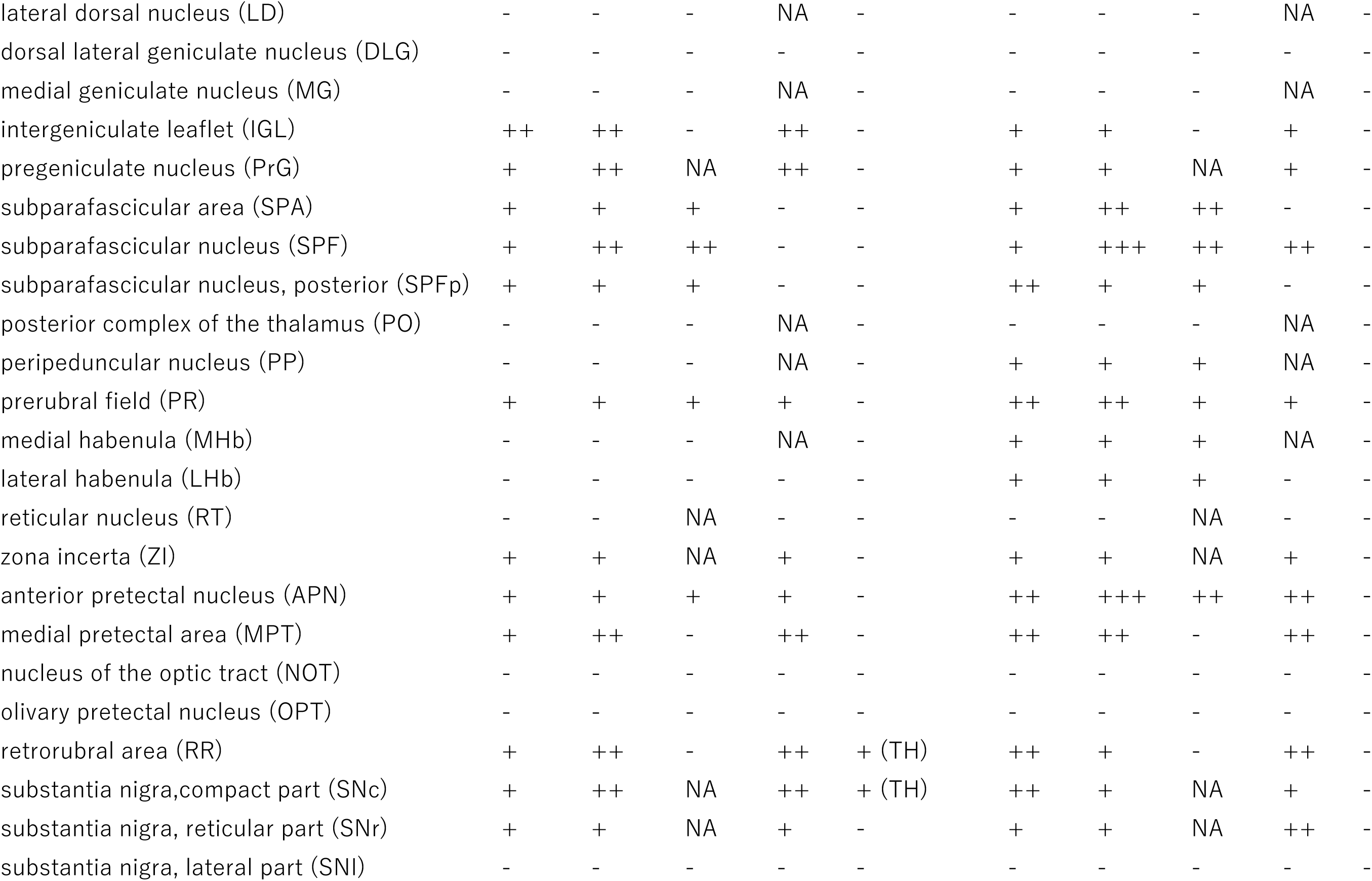
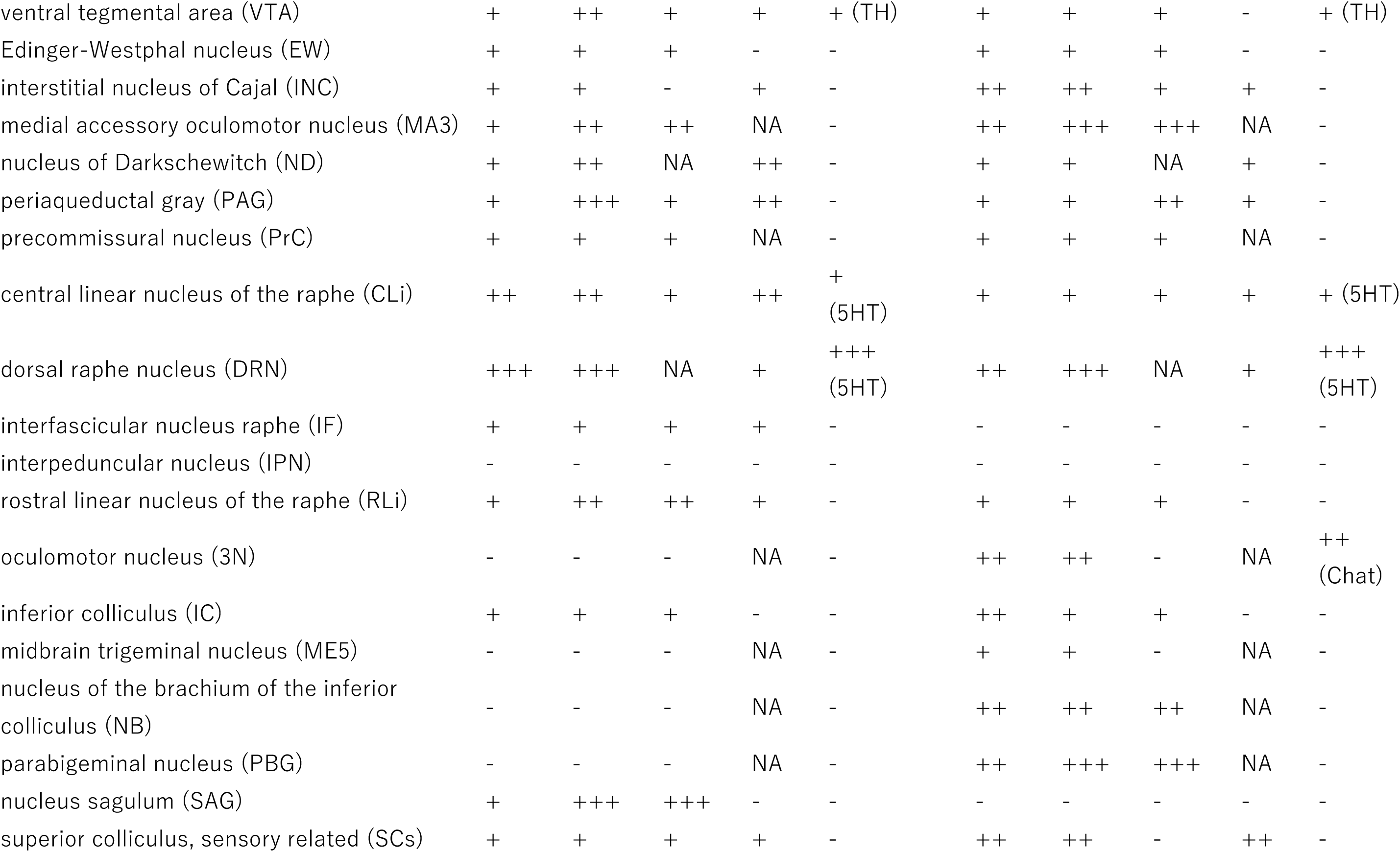
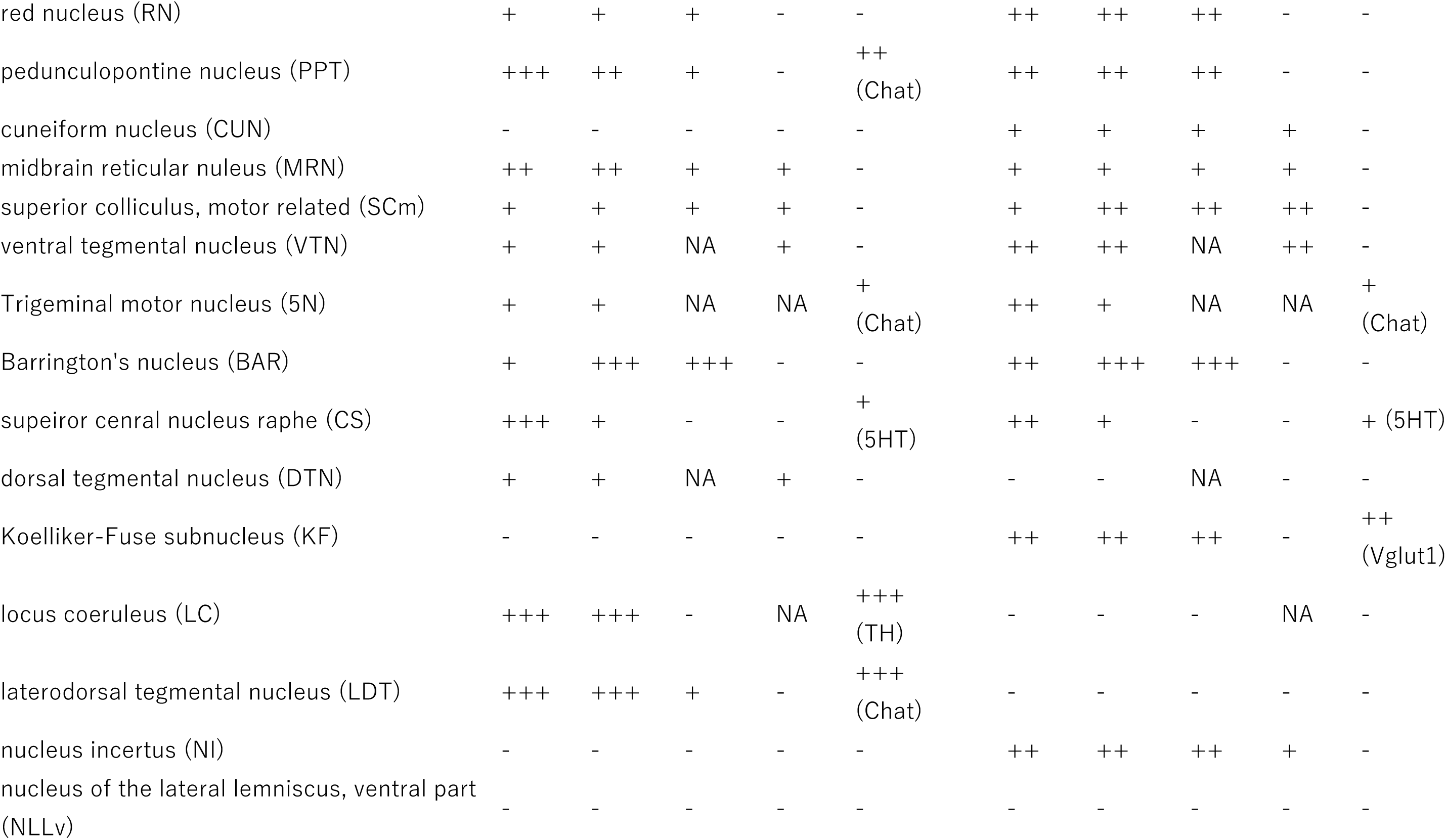
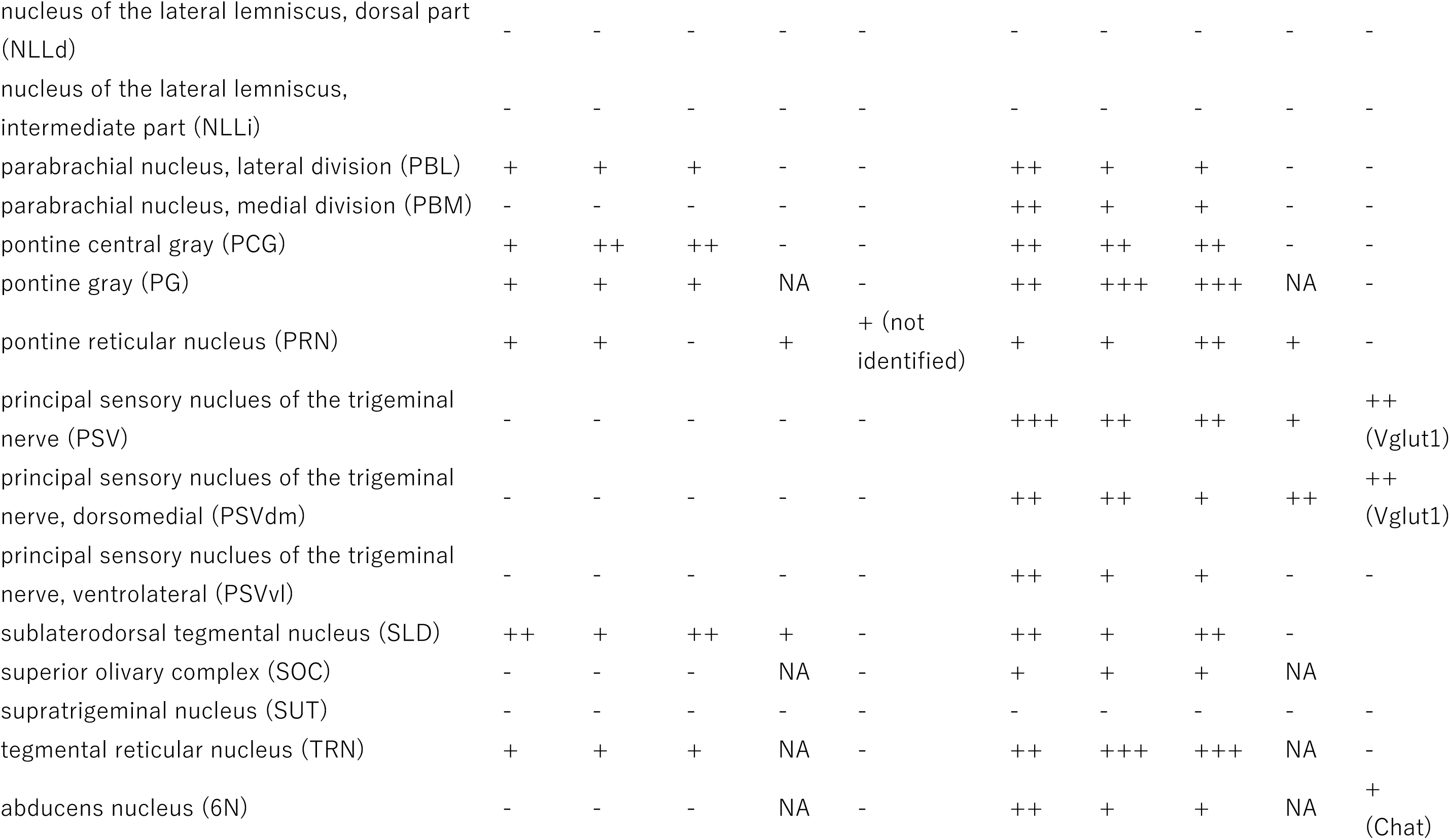
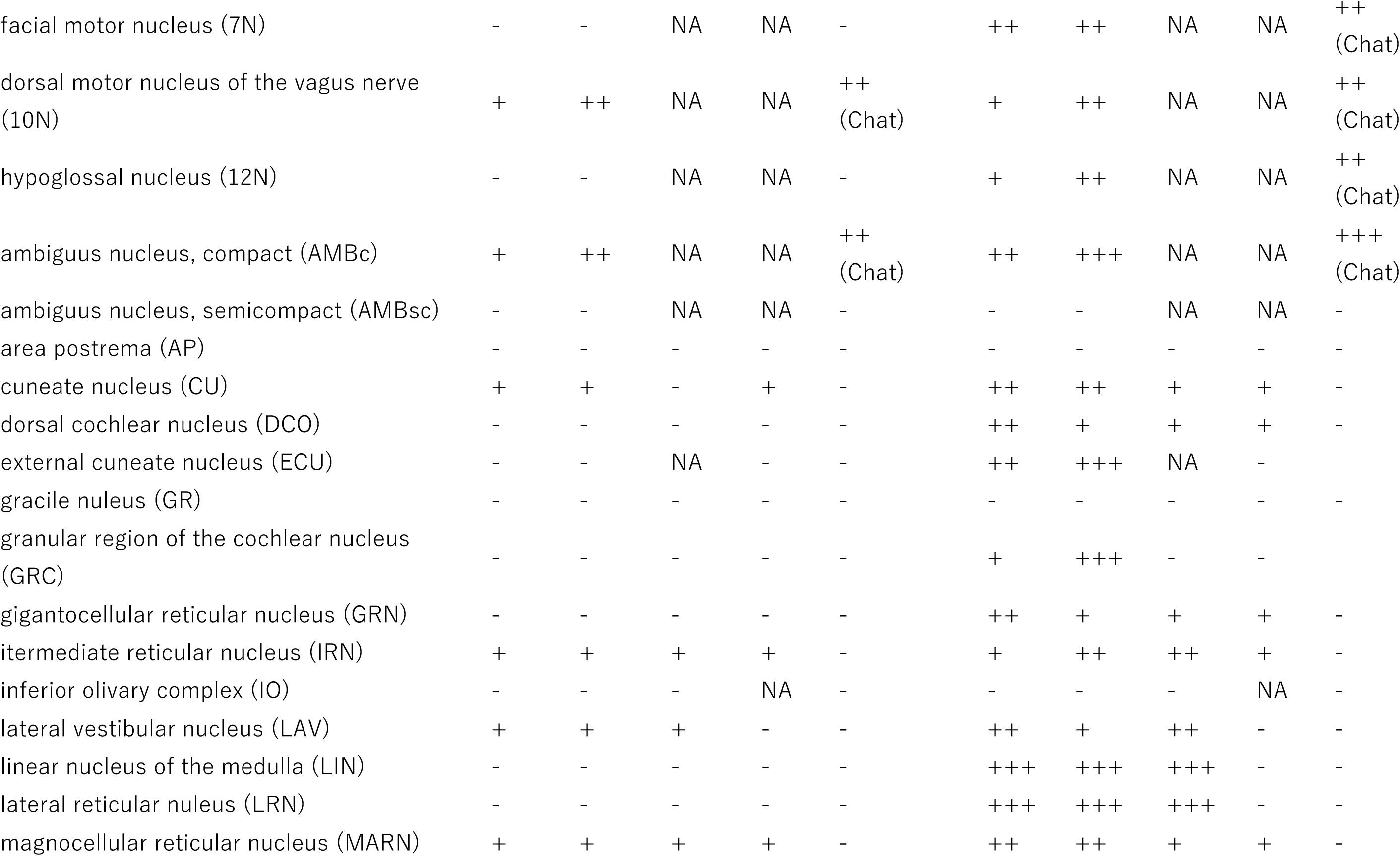
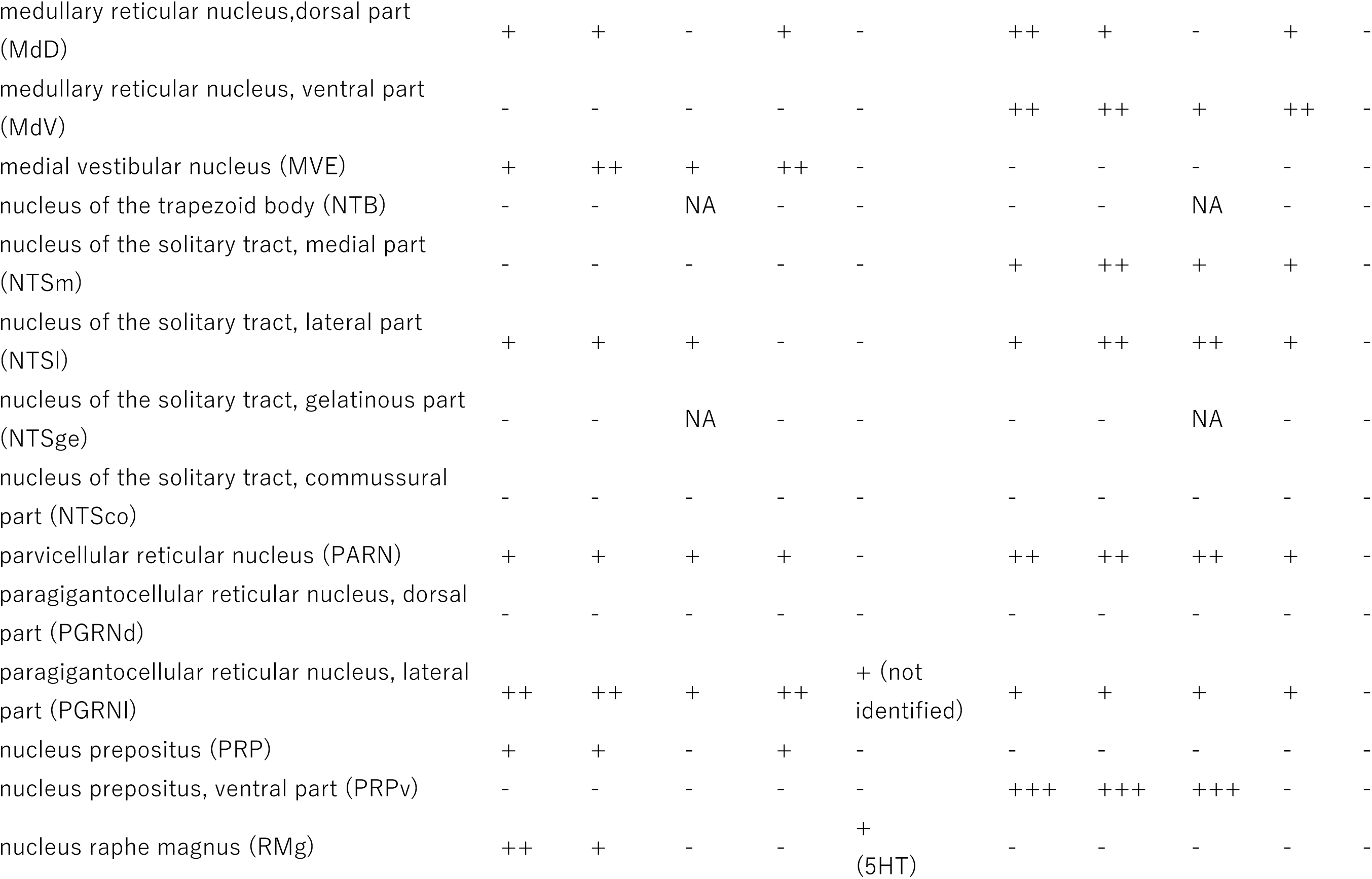
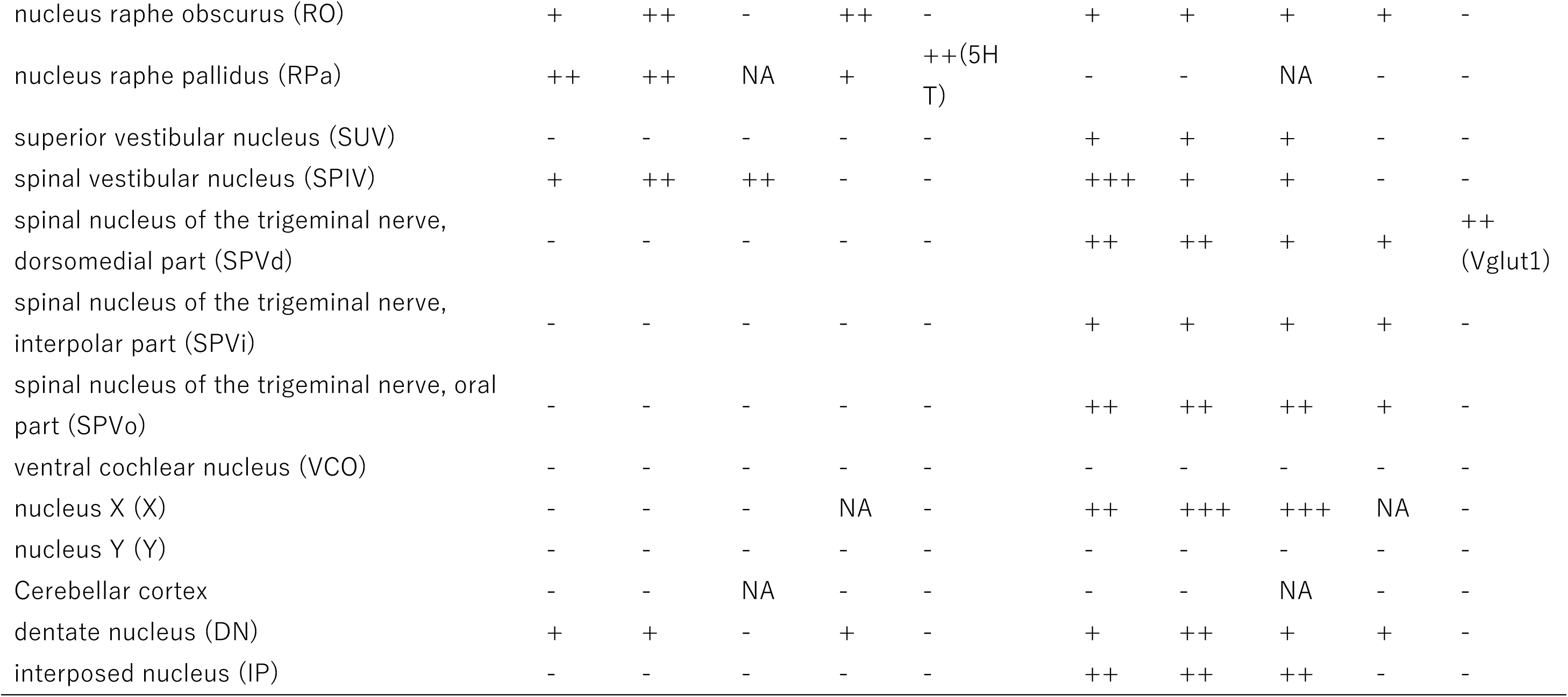
The cellular amount of *Ox1r* or *Ox2r* mRNA for certain cell types and the number of orexin receptor-expressing cells. The data for amount of *Ox1r* or *Ox2r* mRNA were obtained from three mice. Single mouse was used for the evaluation of mRNA expression in certain cell types. Sections including each area were examined at 120 μm interval. * The amount of Ox1r or Ox2r mRNA per cell was evaluated as +++, > 15 granules or granules were overlapping and uncountable ; ++, 6-15 granules; +, 2-5 granules ; -, background level. ** The number of Ox1r-positive or Ox2r-positive cells in the area was evaluated as +++, large ; ++, moderate; +, small; -, background level. *** The frequency of Ox1r- or Ox2r-positive cells among Vglut2-positive or Vgat-positive cell was evaluated as +++, >50 % ; ++, 20-50 %; +, <20 %; -, no or very rare. NA (not applicable) was indicated when Vglut2-positive or Vgat-positive cells were not observed in the area. **** The frequency of Ox1r- or Ox2r-positive cells among certain subtype of cell that was not characterized by Vglut2 or Vgat. For example, "/others" in the locus coeruleus row indicates the frequency of Ox1r- or Ox2r-positive cells among TH-positive cell. Abbreviations:5-HT, 5-hydroxytryptamine; Calb, calbindin; Chat, choline acetyltransferase; HDC, histidine decarboxylase; TH, tyrosine hydroxylase.

**Table S2.** Examined brain areas with regional marker information.

|  | Vglut1 | Vglut2 | Vgat | Calb cell | Calb fiber | TH cell | TH fiber | Chat |
| --- | --- | --- | --- | --- | --- | --- | --- | --- |
| Isocortex layer I | - | - | +++ | - | + | - | - | - |
| Isocortex layer II/III | +++ | - | + | ++ | +++ | - | - | + |
| Isocortex layer IV | +++ | - | + | + | ++ | - | - | - |
| Isocortex layer V | ++ | - | ++ | + | + | - | - | - |
| Isocortex layer VI | +++ | - | + | + | + | - | - | - |
| anterior olfactory nucleus (AON) | +++ | - | + | + | ++ | - | - | - |
| dorsal taenia tecta (DTT) | ++ | - | ++ | + | ++ | - | - | - |
| ventral taenia tecta (VTT) | ++ | - | ++ | + | ++ | - | - | - |
| olfactory tubercle (OT) | - | - | +++ | + | - | - | ++ | + |
| piriform area (Pir) | +++ | + | + | + | + | - | + | - |
| anterior cortical amygdala (ACo) | - | ++ | ++ | + | + | - | - | - |
| nucleus of the lateral olfactory tract (LOT) | ++ | ++ | + | + | + | - | - | - |
| posterolateral cortical amygdala (PLCo) | ++ | ++ | ++ | + | + | - | +/- | - |
| posteromedial cortical amygdala (PMCo) | ++ | - | ++ | + | + | - | - | - |
| dentate gyrus | +++ | - | + | + | ++ | - | - | - |
| CA1 | +++ | - | + | + | + | - | - | - |
| CA2 | +++ | - | + | - | ++/- | - | - | - |
| CA3 | +++ | - | + | - | ++/- | - | - | - |
| subiculum, dorsal (SUBd) | +++ | +++ | + | + | - | - | - | - |
| subiculum, ventral (SUBv) | +++ | - | + | + | + | - | - | - |
| endopiriform nucleus (Epi) | +++ | - | + | + | + | - | - | - |
| Clastrum (CLA) | +++ | + | + | + | - | - | - | - |
| basolateral amygdala, anterior part (BLA) | +++ | - | + | + | ++ | - | - |  |
| basomedial amygdala, anterior part (BMA) | + | ++ | ++ | + | ++ | - | - | - |
| basolateral amygdala, posterior part (BLP) | +++ | - | + | + | ++ | - | + | - |
| basomedial amygdala, posterior part (BMP) | +++ | +++ | + | + | + | - | - | - |
| lateral amygdala (LA) | +++ | +++ | + | + | + | - | - | - |
| amygdala hippocampus area (AHI) | ++ | + | ++ | + | + | - | - | - |
| Caudoputamen (CPu) | - | - | +++ | +/++ | ++ | - | ++ | + |
| nucleus accumbens, core (NAc) | - | - | +++ |  | ++ | - | ++ | + |
| nucleus accumbens, shell (NAsh) | - | - | +++ | + | - | - | ++ | + |
| nucleus accumbens, shell, lateral area (NAshl) | - | - | +++ | + | + | - | ++ | + |
| substantia innominata (SI) | - | + | +++ | - | - | - | +/- | + |
| magnocellular nucleus (MN) | - | + | +++ | + | - | - | - | + |
| globus pallidus (GP) | - | - | +++ | + | + | - | + | + |
| nucleus of the diagonal band (NDB) | - | + | +++ | - | - | - | - | ++ |
| medial septum (MS) | - | + | +++ | - | - | - | - | ++ |
| lateral septum, dorsal (LSd) | - | - | +++ | + | + | - | - | - |
| lateral septum, intermediate (LSI) | - | - | +++ | + | - | - | - | - |
| lateral septum, ventral (LSv) | - | + | +++ | ++ | + | - | - | - |
| septofimbrial nucleus (SF) | - | ++ | ++ | + | - | - | - | - |
| triangular septal nucleus (TRS) | ++ | +++ | + | ++ | ++ | - | - | - |
| bed nucleus of the anterior commissure (BAC) | - | +++ | + | - | - | - | - | - |
| anterior amygdalar area (AAA) | + | + | +++ | + | + | - | - | - |
| central amygdala (CEA) | - | - | +++ | + | + | - | +/- | - |
| intercalated amygdalar nucleus (IA) | - | - | +++ | - | - | - | - | - |
| medial amygdala, anterodorsal (MeAD) | - | + | +++ | + | + | - | - | - |
| medial amygdala, anteroventral (MeAV) | - | +++ | + | + | + | - | - | - |
| medial amygdala, posterodorsal (MePD) | - | + | +++ | + | + | - | - | - |
| medial amygdala, posterodorsal (MePV) | - | +++ | + | + | + | - | - | - |
| bed nucleus of the stria terminalis (BNST) | - | - | +++ | +/- | +/- | - | +/- | +/- |
| bed nucleus of the stria terminalis, principal nucleus (BNSTpr) | - | + | +++ | ++ | ++ | - | - |  |
| medial preoptic area (MPOA) | - | + | +++ | ++/+ | ++/+ |  | +/- | - |
| lateral preoptic area (LPOA) | - | + | +++ | - | - | - | +/- | + |
| vascular organ of the lamina terminalis (VOLT) | - | ++ | ++ | ++ | + | - | - | - |
| median preoptic nucleus (MnPO) | - | +++ | + | + | + | - | - | - |
| paraventricular hypothalamic nucleus (PVH) | - | ++ | ++ | ++ | + | + | + | - |
| suprachiasmatic nucleus (SCN) | - | - | +++ | + | + | - | - | - |
| arcuate nucleus (Arc) | - | ++ | ++ | ++ | ++ | ++ | ++ | - |
| dorsomedial hypothalamic nucleus (DMH) | - | ++ | ++ | ++ | + | - | - | - |
| anterior hypothalamic area (AH) | - | + | +++ | +/- | + | - | - | - |
| lateroanterior hypothalamic nucleus (LAH) | - | +++ | + | + | + | - | - | - |
| lateral hypothalamic area (LH) | - | ++ | ++ | + | +/- | - | + | + |
| tuberal nucleus (TU) | - | ++ | ++ | ++ | + | + | + | - |
| ventromedial hypothalamic nucleus (VMH) | - | +++ | - | + | + | - | - | - |
| entopeduncular nucleus (EP) | - | - | +++ | - | + | - | - | - |
| parasubthalamic nucleus (PST) | - | ++ | ++ | + | - | - | - | - |
| subthalamic nucleus (STN) | - | +++ | - | - | - | - | - | - |
| posterior hypothalamic nucleus (PH) | - | ++ | ++ | + | + | +/- | +/- | - |
| premamillary nucleus, dorsal (PMD) | - | +++ | - | + | - | - | - | - |
| premamillary nucleus, ventral (PMV) | - | ++ | ++ | ++ | ++ | - | - | - |
| tuberomammillary nucleus, dorsal (TMD) | - | + | +++ | + | + | + | + | - |
| tuberomammillary nucleus, ventral (TMV) | - | + | +++ | + | + | - | - | - |
| lateral mammillary nucleus (LM) | - | +++ | - | - | - | - | - | - |
| supramammillary nucleus (SUM) | - | +++ | +++ | + | + | - | - | - |
| median mammillary nucleus (MnM) | - | +++ | - | ++ | ++ | - | - | - |
| medial mammillary nucleus (MM) | - | +++ | - | ++/- | ++/- | - | - | - |
| paraventricular nucleus of the thalamus (PVT) | - | +++ | - | + | + |  | + | - |
| parataenial nucleus (PT) | - | +++ | - | + | + | - | - | - |
| nucleus reuniens (RE) | +++ | +++ | - | ++ | + | - | - | - |
| xiphoid nucleus (Xi) | - | +++ | - | + | + | - | - | - |
| paraxiphoid nucleus (PaXi) | - | +++ | - | + | + | - | - | - |
| central medial nucleus (CM) | - | +++ | - | + | + | - | - | - |
| central lateral nucleus (CL) | +++ | +++ | - | + | + | - | - | - |
| parafascicular nucleus (PF) | - | +++ | - | + | - | - | - | - |
| paracentral nucelus (PC) | - | +++ | - | + | + | - | - | - |
| intermediodorsal nucleus (IMD) | - | +++ | - | + | + | - | - | - |
| rhomboid nucleus (RH) | - | +++ | - | + | + | - | - | - |
| oval paracentral nucleus (OPC) | +++ | +++ | - | ++ | ++ | - | - | - |
| anterodorsal nucleus (AD) | +++ | +++ | - | + | - | - | - | - |
| anteromedial nucleus (AM) | - | +++ | - | - | + | - | - | - |
| anteroventral nucleus (AV) | - | +++ | - | - | + | - | - | - |
| interanterodorsal nucleus (IAD) | - | +++ | - | - | + | - | - | - |
| interanteromedial nucleus (IAM) | - | +++ | - | + | + | - | - | - |
| ventral anterior nucleus (VA) | +++ | +++ | - | ++ | + | - | - | - |
| ventral medial nucleus (VM) | - | +++ | - | ++ | + | - | - | - |
| ventral lateral nucleus (VL) | +++ | +++ | - | - | - | - | - | - |
| ventral posteromedial nucleus (VPM) | +++ | +++ | - | - | - | - | - | - |
| ventral posterolateral nucleus (VPL) | +++ | +++ | - | - | - | - | - | - |
| ventral posteromedial nucleus, parvicellular (VPPC) | - | +++ | - | + | + | - | - | - |
| mediodorsal nucleus (MD) | +++ | +++ | - | +/- | - | - | - | - |
| submedial nucleus (SM) | +++ | +++ | - | - | - | - | - | - |
| lateral posterior nucleus (LP) | - | +++ | - | + | + | - | - | - |
| lateral dorsal nucleus (LD) | +++ | +++ | - | - | - | - | - | - |
| dorsal lateral geniculate nucleus (DLG) | +++ | +++ | + | - | - | - | - | - |
| medial geniculate nucleus (MG) | +++ | +++ | - | - | + | - | - | - |
| intergeniculate leaflet (IGL) | - | + | +++ | + | + | - | - | - |
| pregeniculate nucleus (PrG) | - | - | +++ | - | + | - | - | - |
| subparafascicular area (SPA) | - | +++ | + | + | + | + | + | - |
| subparafascicular nucleus (SPF) | - | ++ | ++ | ++ | + | - | - | - |
| subparafascicular nucleus, posterior (SPFp) | - | +++ | + | + | + | - | - | - |
| posterior complex of the thalamus (PO) | +++ | +++ | - | - | - | - | - | - |
| peripeduncular nucleus (PP) | - | +++ | - | + | ++ | - | - | - |
| prerubral field (PR) | - | +++ | + | - | - | - | - | - |
| medial habenula (MHb) | +++ | +++ | - | ++ | + | - | - | +++ |
| lateral habenula (LHb) | - | +++ | + | - | - | - | - | - |
| reticular nucleus (RT) | - | - | +++ | - | - | - | - | - |
| zona incerta (ZI) | - | - | +++ | - | - | ++/- | ++/- | - |
| anterior pretectal nucleus (APN) | - | ++ | ++ | - | - | - | - | - |
| medial pretectal area (MPT) | - | + | +++ | + | + | - | - | - |
| nucleus of the optic tract (NOT) | - | +++ | + | + | + | - | - | - |
| olivary pretectal nucleus (OPT) | - | ++ | ++ | + | + | - | - | - |
| retrotrubral area (RR) | - | ++ | ++ | - | - | ++ | ++ | - |
| substantia nigra, compact part (SNc) | - | - | +++ | + | + | ++ | ++ | - |
| substantia nigra, reticular part (SNr) | - | - | +++ | - | ++ | - | ++ | - |
| substantia nigra, lateral part (SNI) | - | ++ | ++ | + | + | ++ | ++ | - |
| ventral tegmental area (VTA) | - | + | +++ | ++ | ++ | ++ | ++ | - |
| Edinger-Westphal nucleus (EW) | - | ++ | + | - | - | - | + | - |
| interstitial nucleus of Cajal (INC) | - | ++ | ++ | - | - | - | - | - |
| medial accessory oculomotor nucleus (MA3) | - | +++ | - | - | - | - | - | - |
| nucleus of Darkschewitch (ND) | - | - | +++ | - | - | - | - | - |
| periaqueductal gray (PAG) | - | +++ | + | + | + | + | + | - |
| precommissural nucleus (PrC) | - | +++ | - | - | - | - | - | - |
| central linear nucleus of the raphe (CLi) | - | ++ | ++ | + | ++ | + | ++ | - |
| dorsal raphe nucleus (DRN) | - | - | + | + | + | + | + | - |
| interfascicular nucleus raphe (IF) | - | ++ | ++ | + | + | + | ++ | - |
| interpeduncular nucleus (IPN) | - | + | +++ | - | ++/- | - | - | - |
| rostral linear nucleus of the raphe (RLi) | - | ++ | ++ | + | + | + | + | - |
| oculomotor nucleus (3N) | - | + | - | - | - | - | - | +++ |
| inferior colliculus (IC) | - | ++ | ++ | - | - | - | - | - |
| midbrain trigeminal nucleus (ME5) | +++ | +++ | - | - | - | - | - | - |
| nucleus of the brachium of the inferior colliculus (NB) | - | +++ | - | + | + | - | - | - |
| parabigeminal nucleus (PBG) | - | +++ | - | - | - | - | - | +++ |
| nucleus sagulum (SAG) | - | +++ | + | - | - | - | - | - |
| superior colliculus, sensory related (SCs) | - | + | +++ | + | - | - | - | - |
| red nucleus (RN) | - | +++ | + | - | - | - | - | - |
| pedunclopontine nucleus (PPT) | - | ++ | ++ | + | - | - | + | + |
| cuneiform nucleus (CUN) | - | ++ | ++ | - | - | - | - | - |
| midbrain reticular nucleus (MRN) | - | ++ | ++ | - | - | - | - | - |
| superior colliculus, motor related (SCm) | - | ++ | ++ | - | - | - | - | - |
| ventral tegmental nucleus (VTN) | - | - | +++ | - | + | - | - | - |
| Trigeminal motor nucleus (5N) | - | - | - | - | - | - | - | +++ |
| Barrington's nucleus (BAR) | - | +++ | + | + | - | - | + | - |
| superior central nucleus raphe (CS) | - | + | +++ | - | + | - | - | - |
| dorsal tegmental nucleus (DTN) | - | - | +++ | - | - | - | - | - |
| Koelliker-Fuse subnucleus (KF) | ++ | ++ | ++ | - | - | - | - | - |
| locus coeruleus (LC) | - | +++ | - | + | - | +++ | ++ | - |
| laterodorsal tegmental nucleus (LDT) | - | ++ | ++ | - | - | - | + | ++ |
| nucleus incertus (NI) | - | ++ | ++ | + | + | - | - | - |
| nucleus of the lateral lemniscus, ventral part (NLLv) | - | + | +++ | + | + | - | - | - |
| nucleus of the lateral lemniscus, dorsal part (NLLd) | - | + | +++ | - | - | - | - | - |
| nucleus of the lateral lemniscus, intermediate part (NLLi) | - | ++ | ++ | + | + | - | - | - |
| parabrachial nucleus, lateral division (PBL) | - | +++ | + | + | + | - | + | - |
| parabrachial nucleus, medial division (PBM) | - | +++ | + | + | + | - | - | - |
| pontine central gray (PCG) | + | + | ++ | - | - | - | - | - |
| pontine gray (PG) | +++ | +++ | - | - | - | - | - | - |
| pontine reticular nucleus (PRN) | - | ++ | ++ | - | - | - | - | - |
| principal sensory nucleus of the trigeminal nerve (PSV) | ++ | +++ | + | - | - | - | - | - |
| principal sensory nucleus of the trigeminal nerve, dorsomedial (PSVdm) | +++ | +++ | + | - | - | - | - | - |
| principal sensory nucleus of the trigeminal nerve, ventrolateral (PSVvl) | ++ | +++ | + | - | - | - | - | - |
| sublaterodorsal tegmental nucleus (SLD) | - | + | + | - | - | - | + | - |
| superior olivary complex (SOC) | - | +++ | - | - | + | - | - | - |
| supratrigeminal nucleus (SUT) | - | +++ | + | - | - | - | - | - |
| tegmental reticular nucleus (TRN) | +++ | +++ | - | - | - | - | - | - |
| abducens nucleus (6N) | +++ | +++ | - | - | - | - | - | +++ |
| facial motor nucleus (7N) | - | - | - | - | - | - | - | +++ |
| dorsal motor nucleus of the vagus nerve (10N) | - | - | - | - | - | - | - | +++ |
| hypoglossal nucleus (12N) | - | - | - | - | - | - | - | +++ |
| ambiguus nucleus, compact (AMBc) | - | - | - | - | - | - | - | +++ |
| ambiguus nucleus, semicompact (AMBsc) | - | - | - | - | - | - | - | +++ |
| area postrema (AP) | - | ++ | ++ | ++ | ++ | + | + | + |
| cuneate nucleus (CU) | + | ++ | ++ | - | - | - | - | - |
| dorsal cochlear nucleus (DCO) | ++ | + | ++ | - | - | - | - | - |
| external cuneate nucleus (ECU) | +++ | - | + | - | - | - | - | - |
| gracile nucleus (GR) | - | ++ | ++ | - | - | - | - | - |
| granular region of the cochlear nucleus (GRC) | +++ | + | + | - | - | - | - | - |
| gigantocellular reticular nucleus (GRN) | - | + | +++ | - | - | - | - | - |
| intermediate reticular nucleus (IRN) | - | ++ | ++ | + | - | - | - | - |
| inferior olivary complex (IO) | - | +++ | - | ++ | ++ | - | - | - |
| lateral vestibular nucleus (LAV) | - | +++ | + | - | + | - | - | - |
| linear nucleus of the medulla (LIN) | +++ | +++ | + | - | - | - | - | - |
| lateral reticular nucleus (LRN) | +++ | +++ | + | - | - | - | - | - |
| magnocellular reticular nucleus (MARN) | - | + | +++ | - | - | - | - | - |
| medullary reticular nucleus,dorsal part (MdD) | - | ++ | ++ | - | - | - | - | - |
| medullary reticular nucleus, ventral part (MdV) | - | ++ | ++ | - | - | - | - | - |
| medial vestibular nucleus (MVE) | - | ++ | ++ | - | ++ | - | - | - |
| nucleus of the trapezoid body (NTB) | - | - | +++ | ++ | + | - | - | - |
| nucleus of the solitary tract, medial part (NTSm) | - | + | +++ | - | - | - | - | - |
| nucleus of the solitary tract, lateral part (NTSl) | - | ++ | ++ | - | - | - | - | - |
| nucleus of the solitary tract, gelatinous part (NTSge) | - | - | +++ | - | + | - | - | - |
| nucleus of the solitary tract, commussural part (NTSco) | - | ++ | ++ | ++ | ++ | ++ | ++ | - |
| parvicellular reticular nucleus (PARN) | - | ++ | ++ | - | - | - | - | - |
| paragigantocellular reticular nucleus, dorsal part (PGRNd) | - | + | +++ | + | + | - | - | - |
| paragigantocellular reticular nucleus, lateral part (PGRNI) | - | + | +++ | - | - | + | + | - |
| nucleus prepositus (PRP) | + | + | +++ | + | + | + | - | - |
| nucleus prepositus, ventral part (PRPv) | +++ | +++ | + | - | - | - | - | - |
| nucleus raphe magnus (RMg) | - | ++ | ++ | - | - | - | - | - |
| nucleus raphe obscurus (RO) | + | + | +++ | - | - | - | - | - |
| nucleus raphe pallidus (RPa) | - | - | +++ | - | - | - | - | - |
| superior vestibular nucleus (SUV) | + | + | +++ | - | ++ | - | - | - |
| spinal vestibular nucleus (SPIV) | - | ++ | ++ | - | + | - | - | - |
| spinal nucleus of the trigeminal nerve,<br>dorsomedial part (SPVd) | + | ++ | ++ | - | - | - | - | - |
| spinal nucleus of the trigeminal nerve,<br>interpolar part (SPVi) | + | + | +++ | - | - | - | - | - |
| spinal nucleus of the trigeminal nerve, oral part<br>(SPVo) | - | ++ | ++ | - | - | - | - | - |
| ventral cochlear nucleus (VCO) | +++ | +++ | + | + | - | - | - | - |
| nucleus X (X) | +++ | +++ | - | - | + | - | - | - |
| nucleus Y (Y) | - | ++ | ++ | + | - | - | - | - |
| Cerebellar cortex | +++ | - | + | +++ | +++ | - | - | - |
| dentate nucleus (DN) | - | ++ | ++ | - | +++ | - | - | - |
| interposed nucleus (IP) | - | ++ | ++ | - | +++ | - | - | - |

**Table S3.**
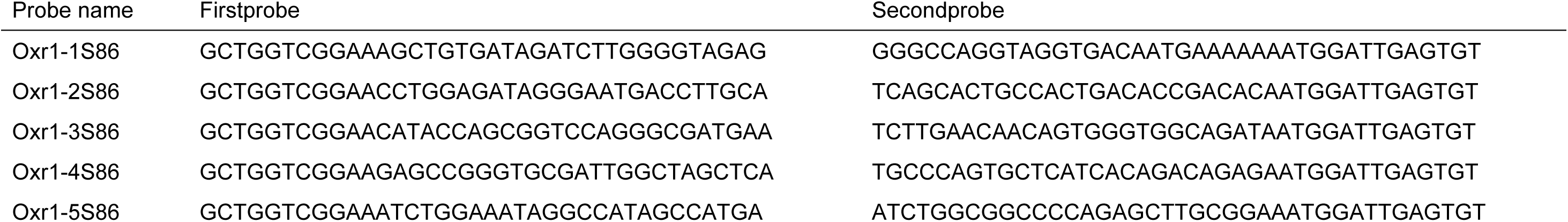
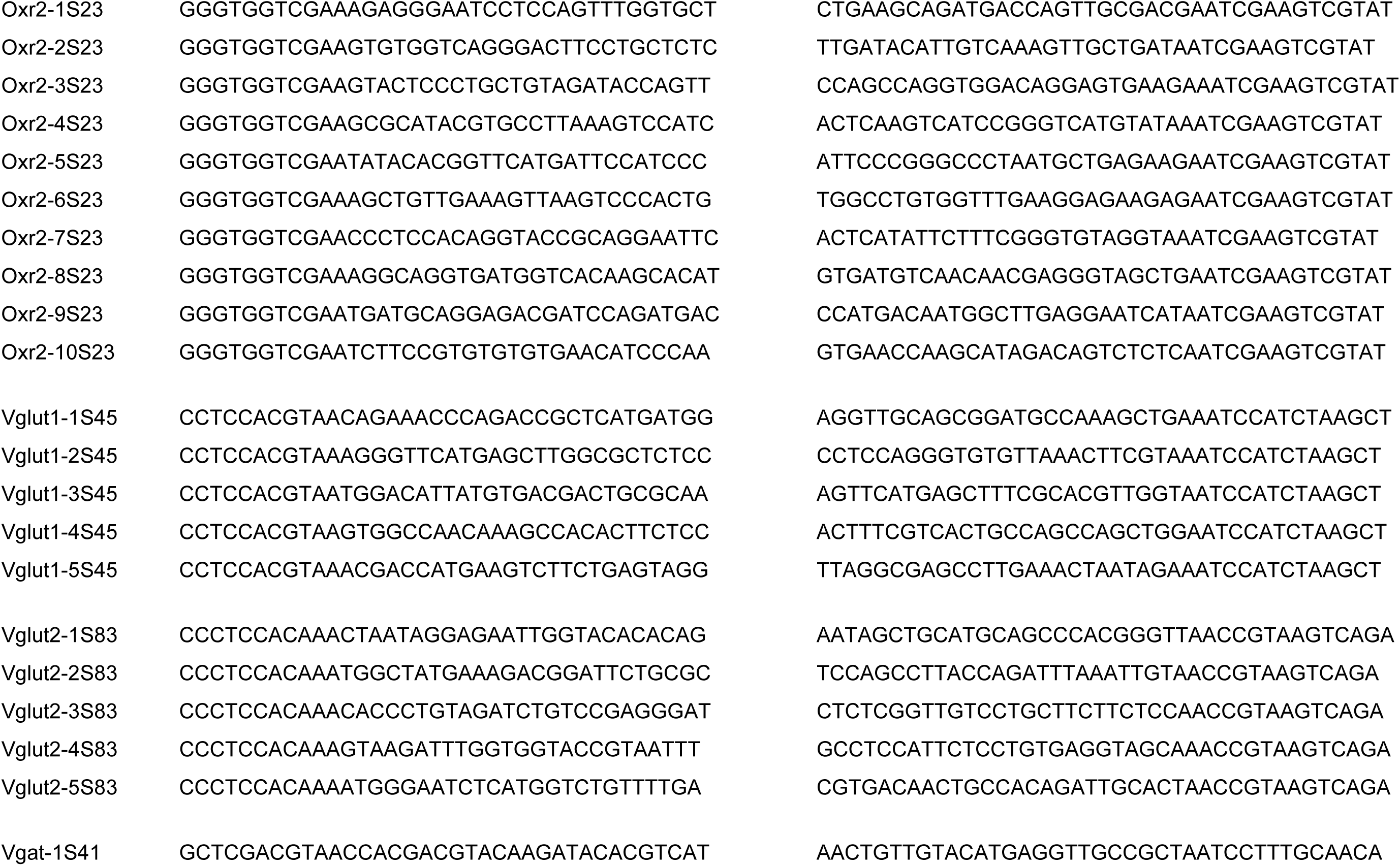
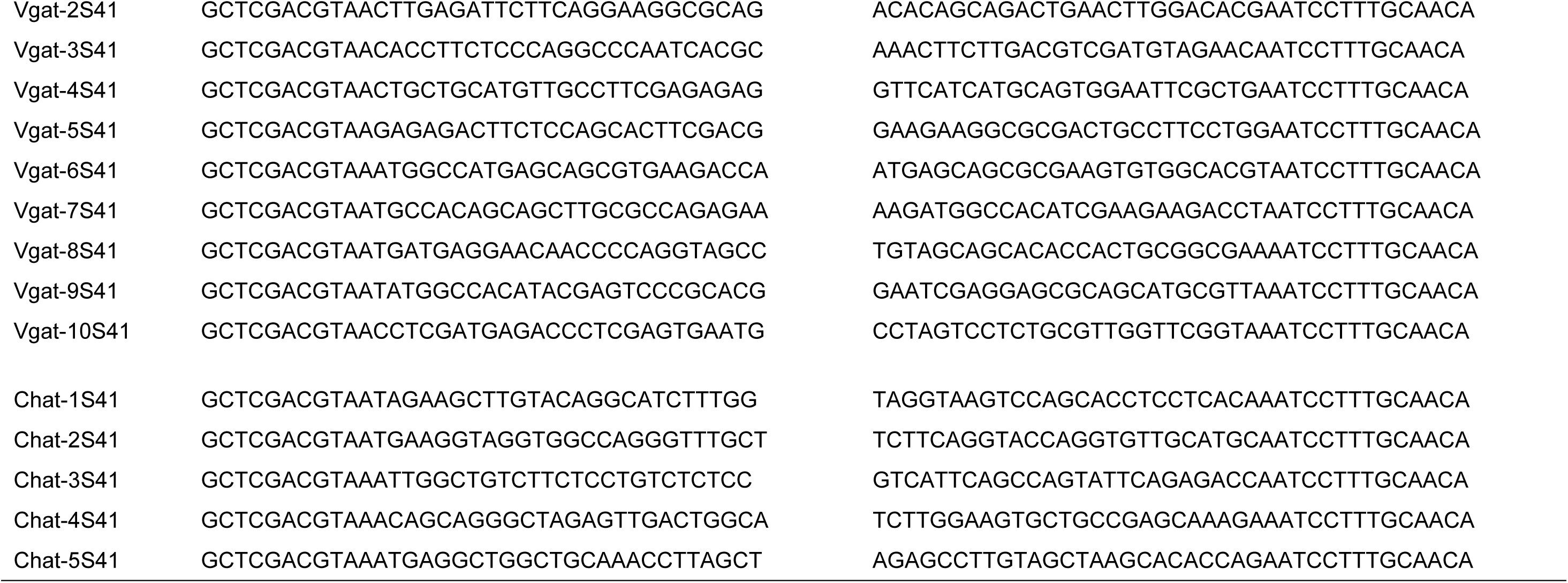
Split-initiator probe sequences.

**Table S4.**
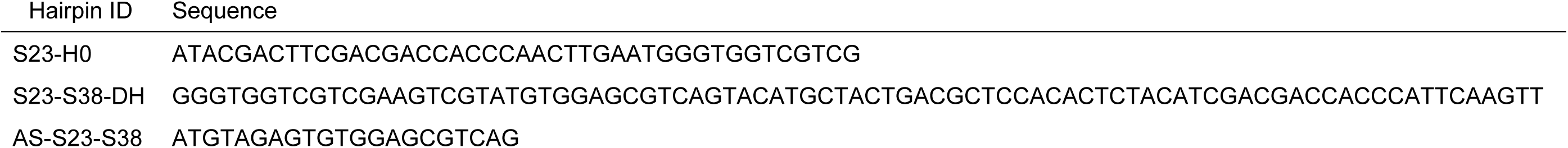
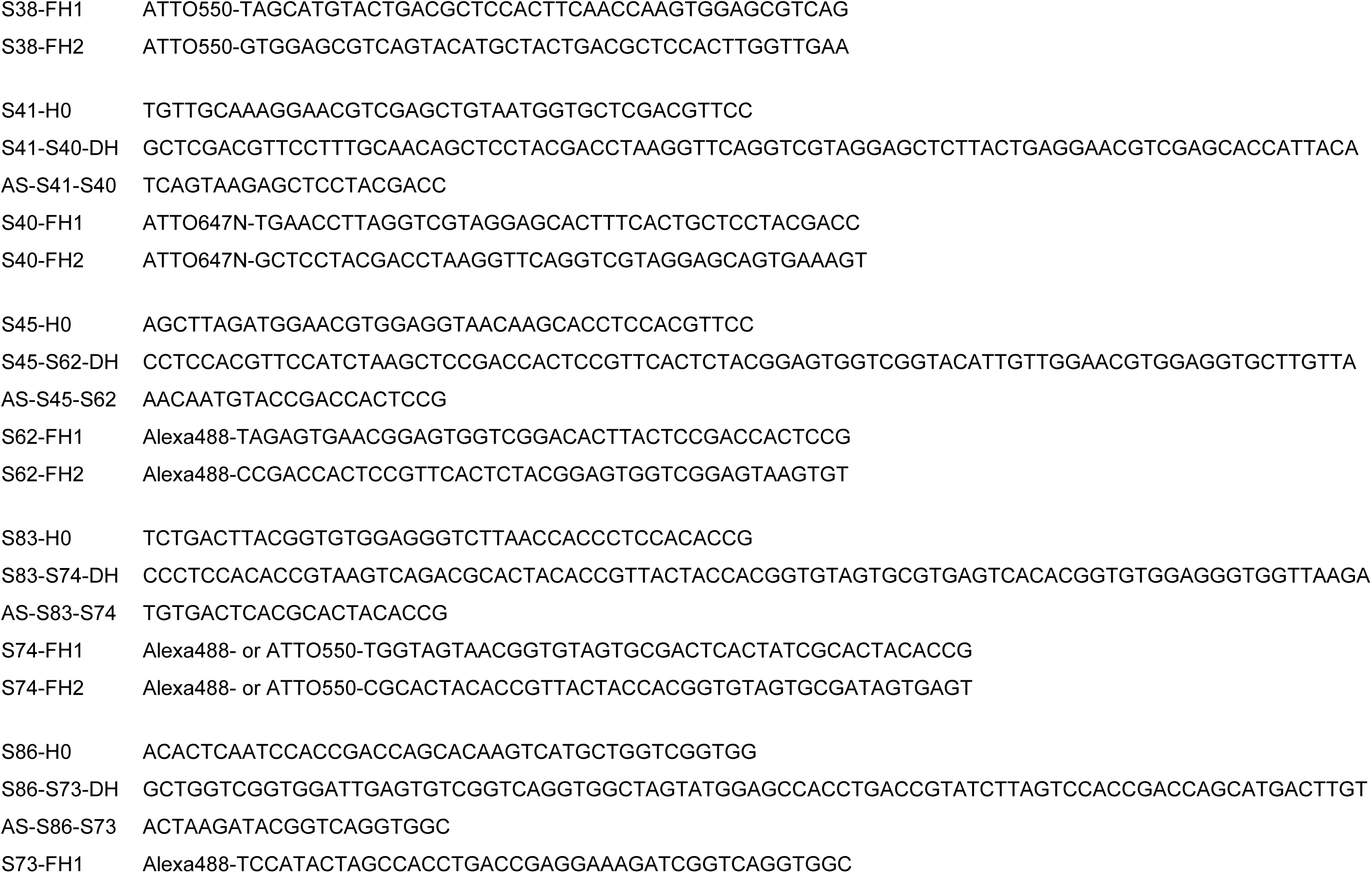
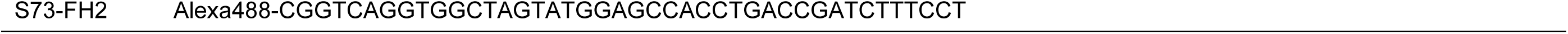
Hairpin sequence and conjugated dye.

**Table S5.** Antibody list.

Antibody list
| Antigen | Host | Source | Cat ID | Concentration |
| --- | --- | --- | --- | --- |
| 5HT (serotonin) | Goat | Immunostar | 20079 | 1:2000 |
| Calbindin D28k | Mouse | Sigma | C9848 | 1:2000 |
| Histidine decarboxylase | Guniea pig | American research product | 03-16046 | 1:2000 |
| Orexin A | Rabbit | Millipore | AB3704 | 1:2000 |
| Tyrosine hydroxylase | Rabbit | Millipore | AB152 | 1:4000 |

## 12 Supplementary figure legends

**Fig. S1.**
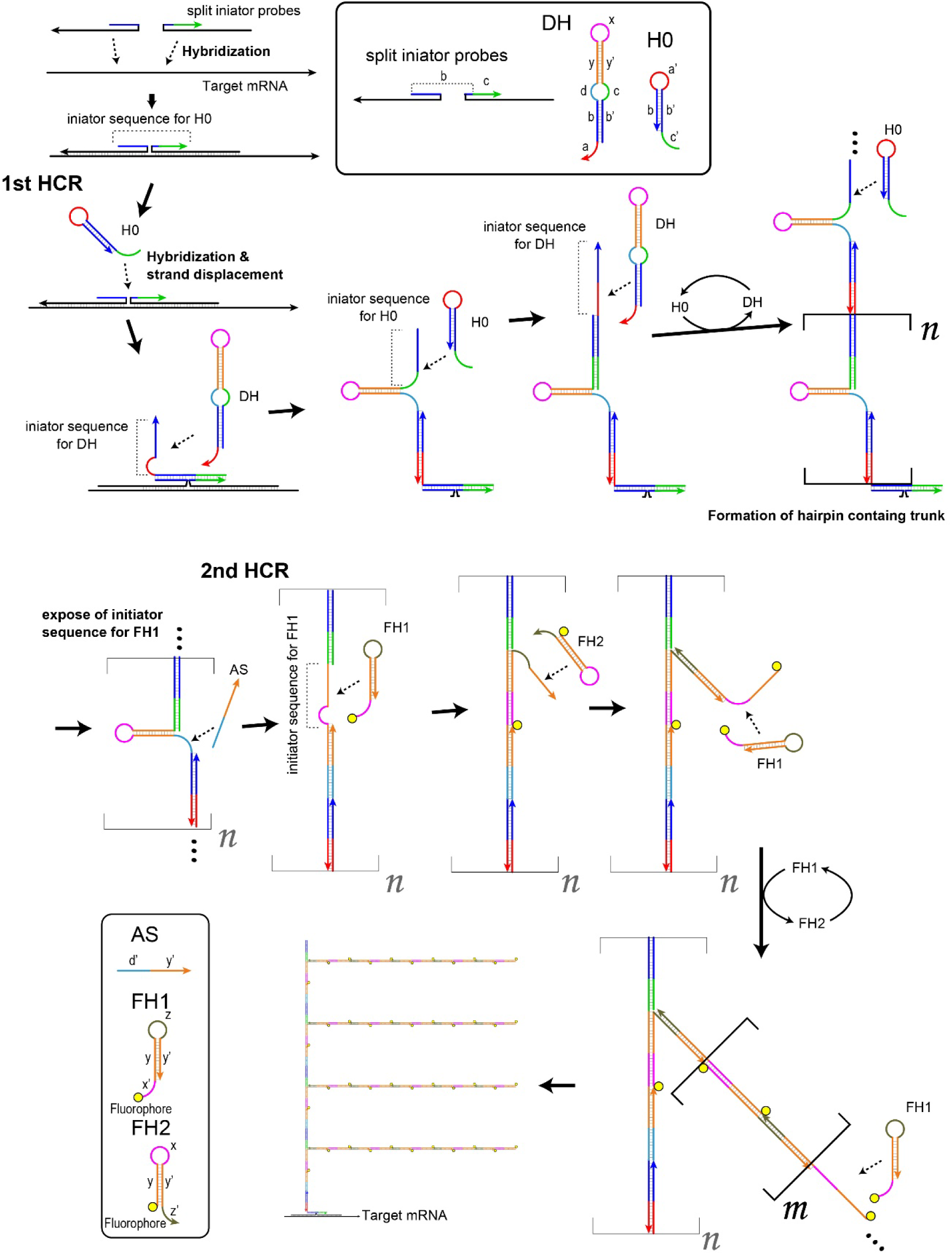
Principle of branched *in situ* hybridization chain reaction (HCR) Split-initiator probes hybridize with the target mRNA to form an initiator sequence (bc) for hairpin DNA H0. In the 1st HCR, H0 hybridizes with the pair of split-initiator probes, followed by exposure of initiator sequence (a’b’) for double stem-hairpin DH. DH hybridizes with the 3’ end of H0, and the initiator sequence for H0 is exposed from DH. As a consequence of HCR, the DNA trunk with the repeated hairpin-containing domain is formed on the target mRNA. In the 2nd HCR, the assist oligo (AS) hybridizes with the hairpin-containing domain to expose the initiator sequence (xy’) of fluorescent hairpin DNA FH1. FH1 hybridizes with the single-strand region of the trunk to form branched DNA with the initiator sequence (zy) of FH2. HCR of FH1 and FH2 occur in multiple regions of the DNA trunk, enabling higher polymerization of fluorescent hairpins per single target site than non-branched HCR.

**Fig. S2.**
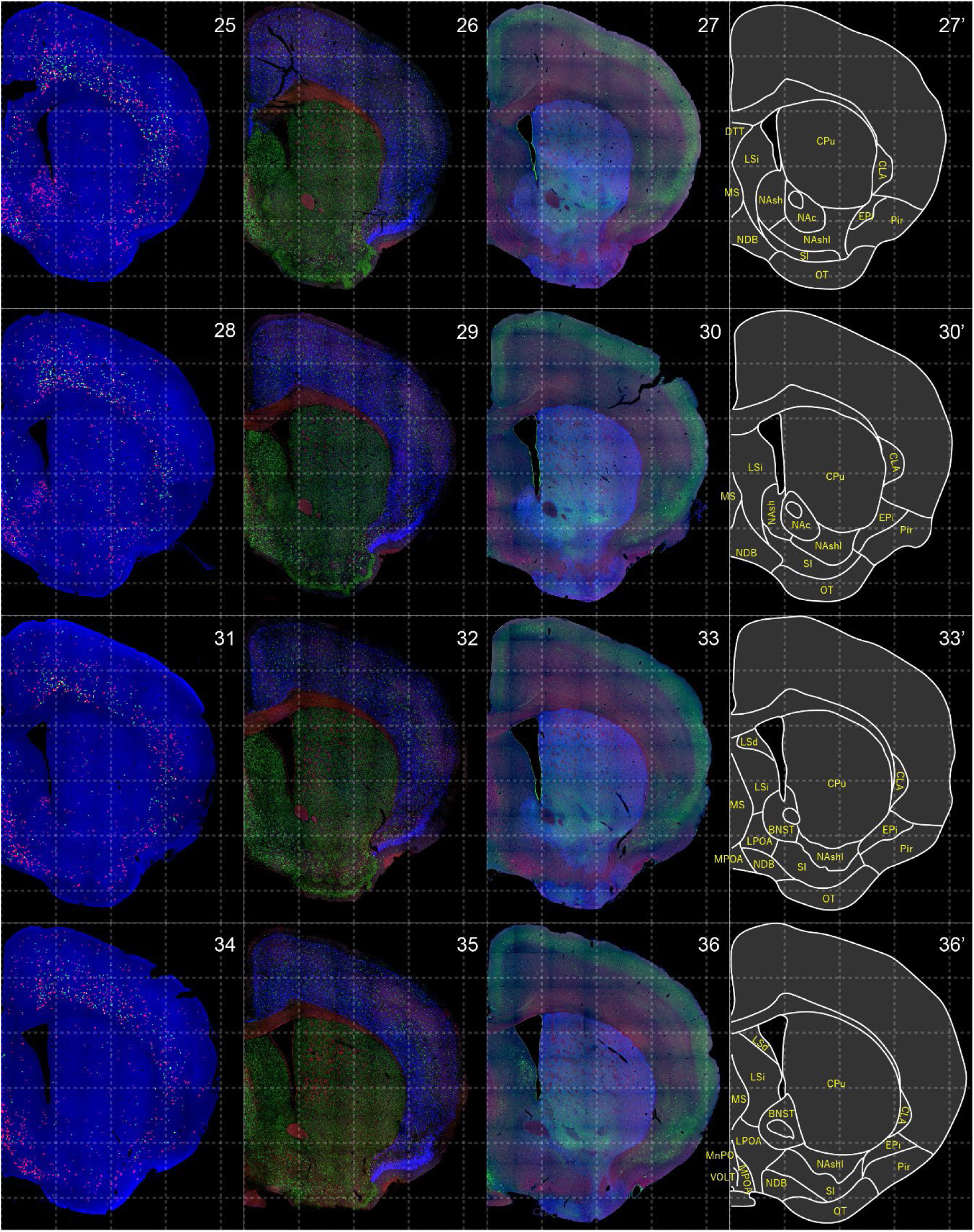
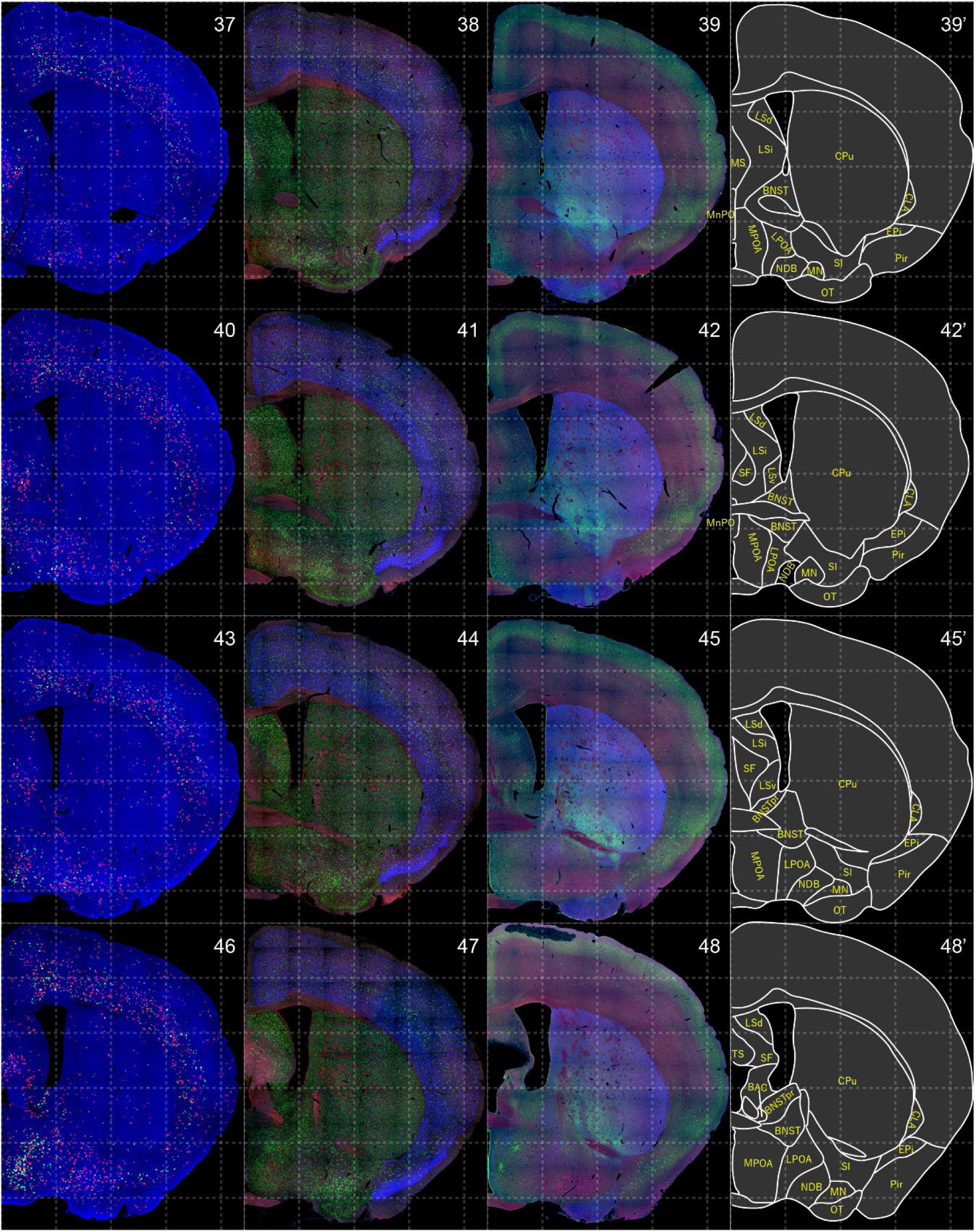
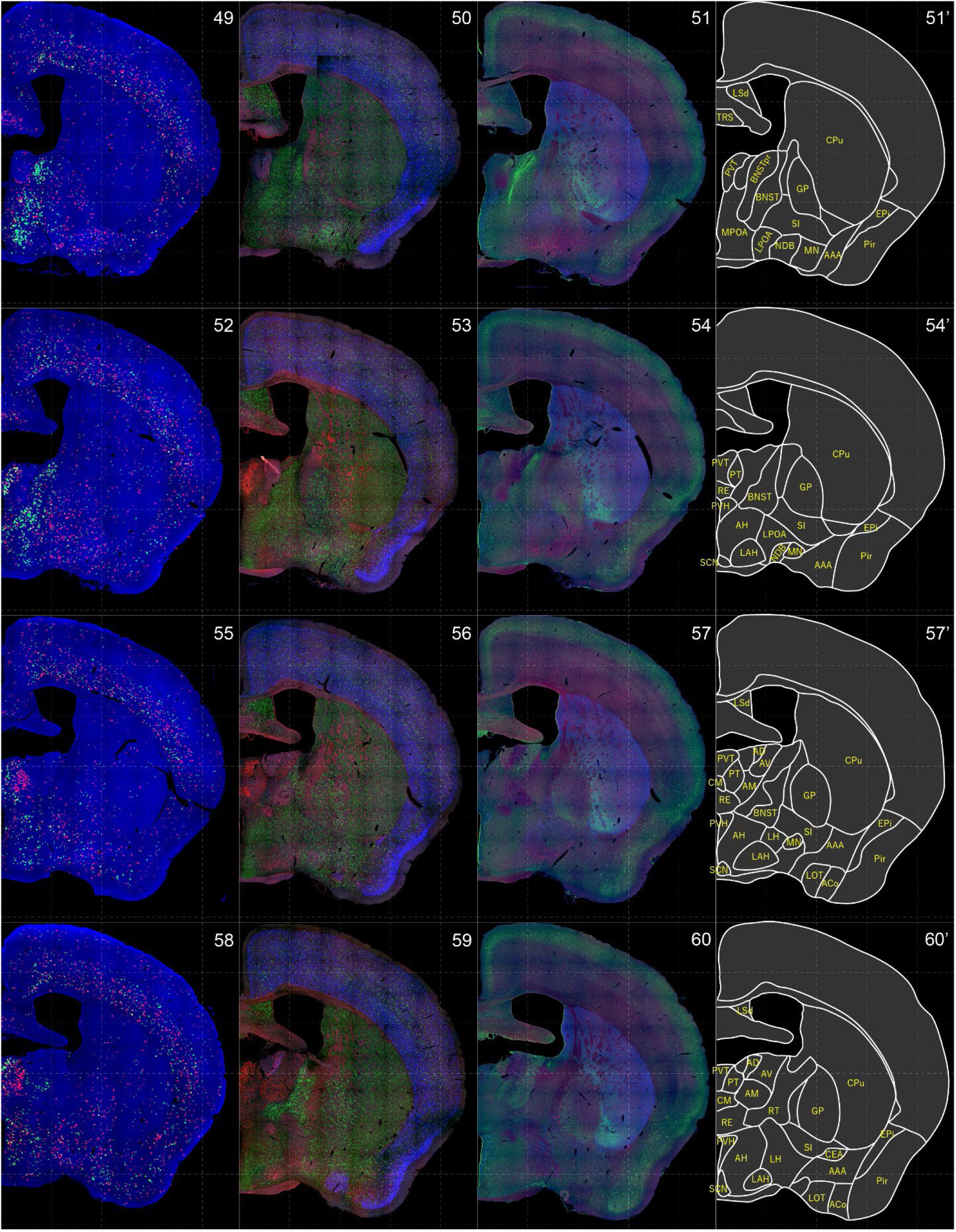
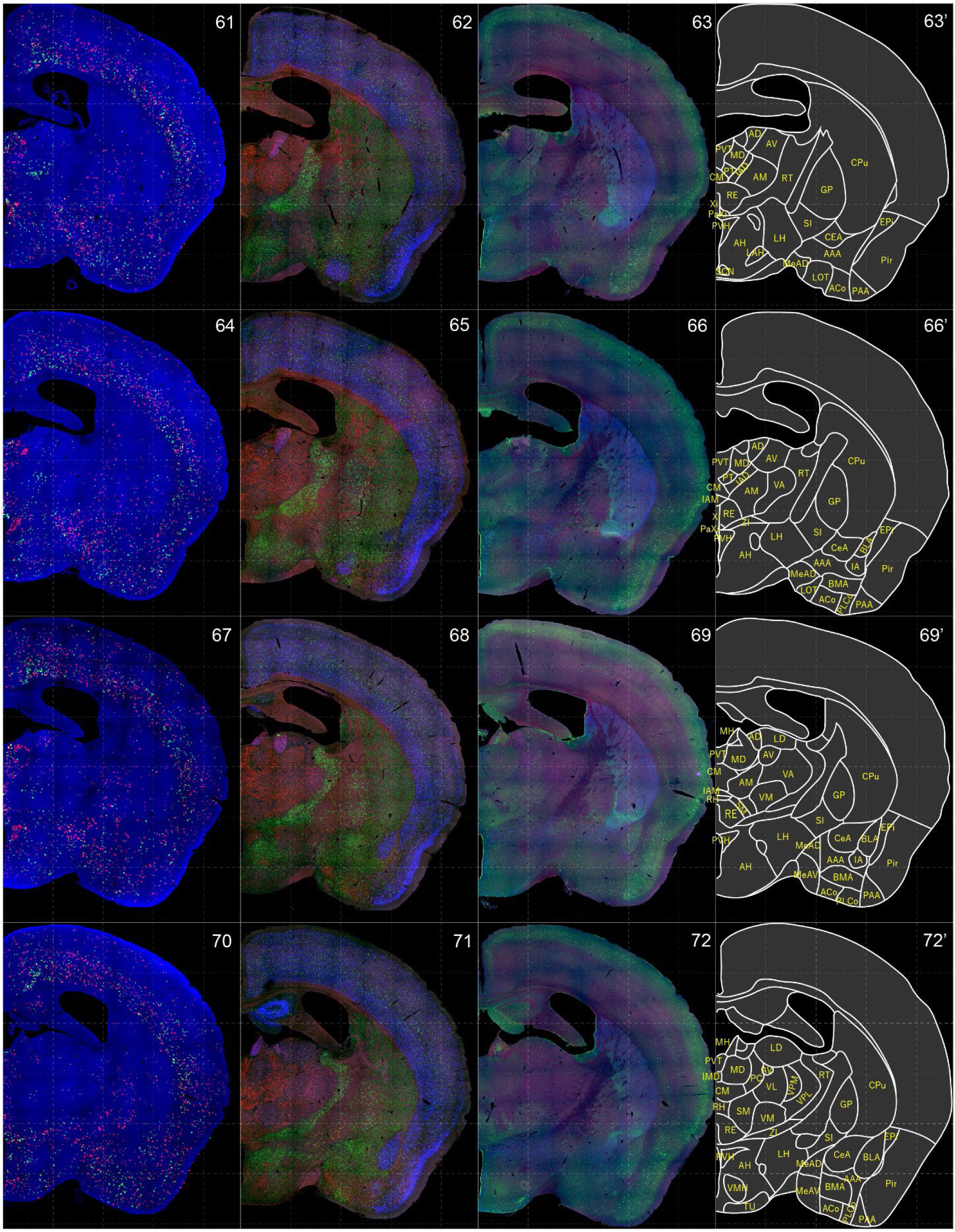
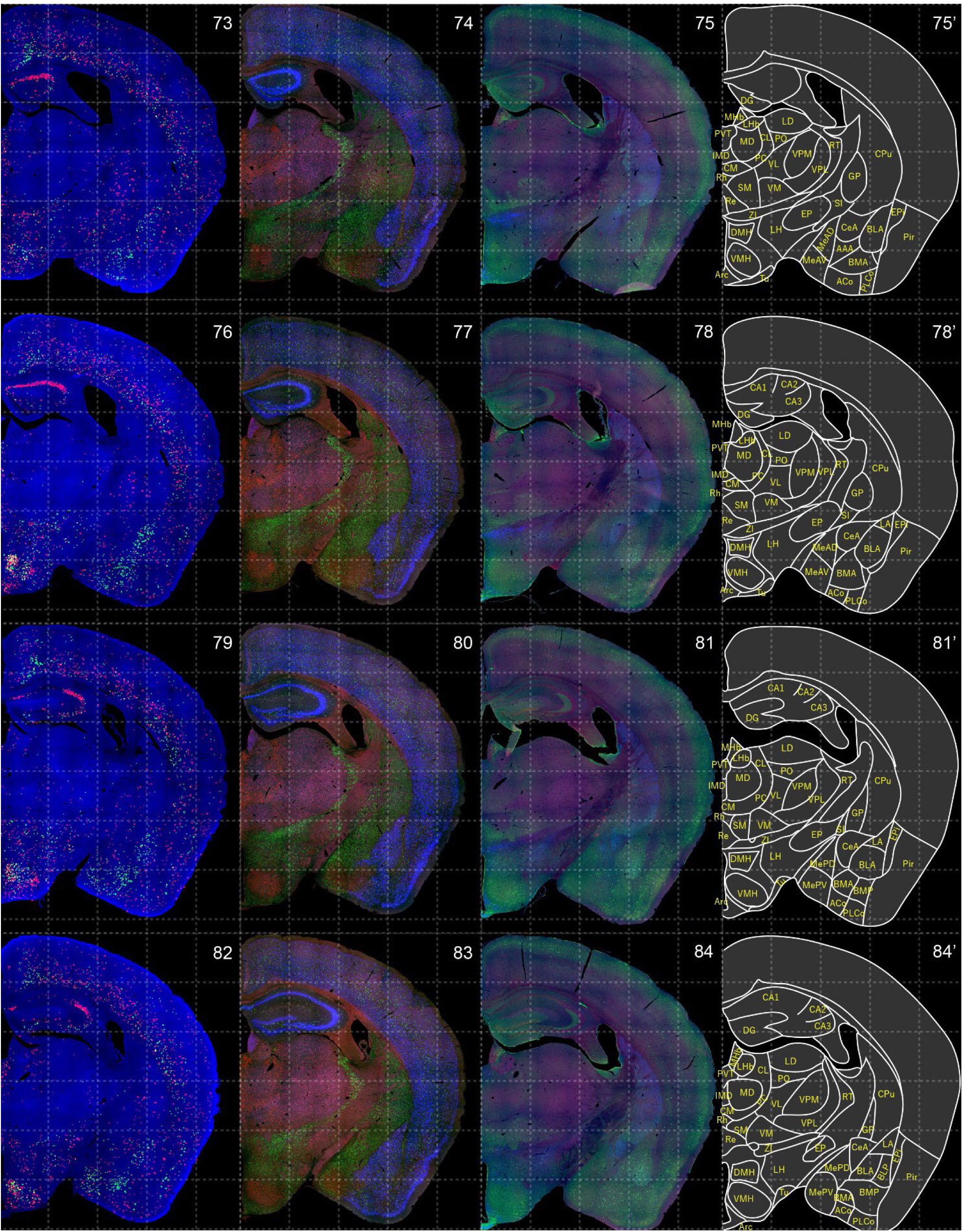
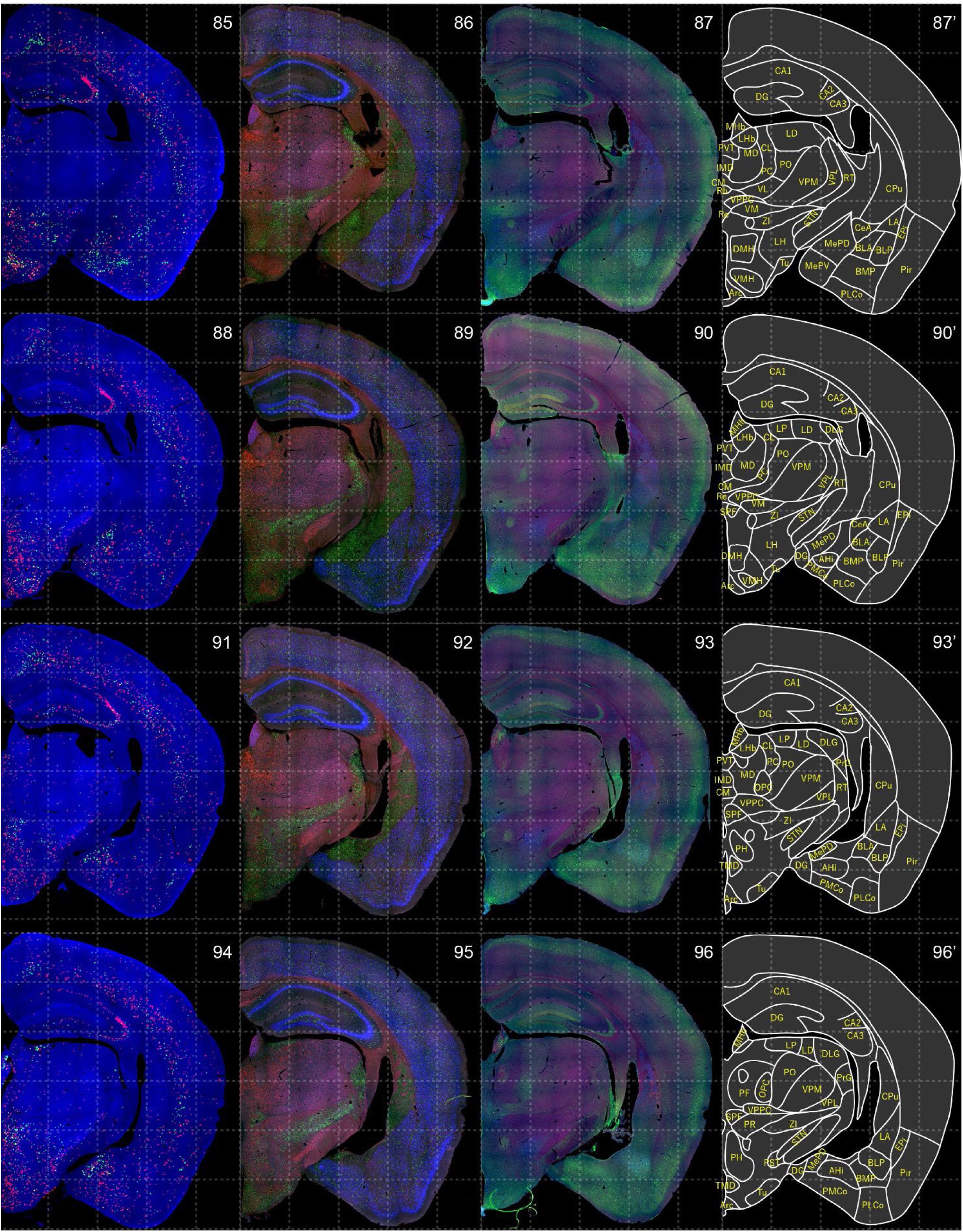
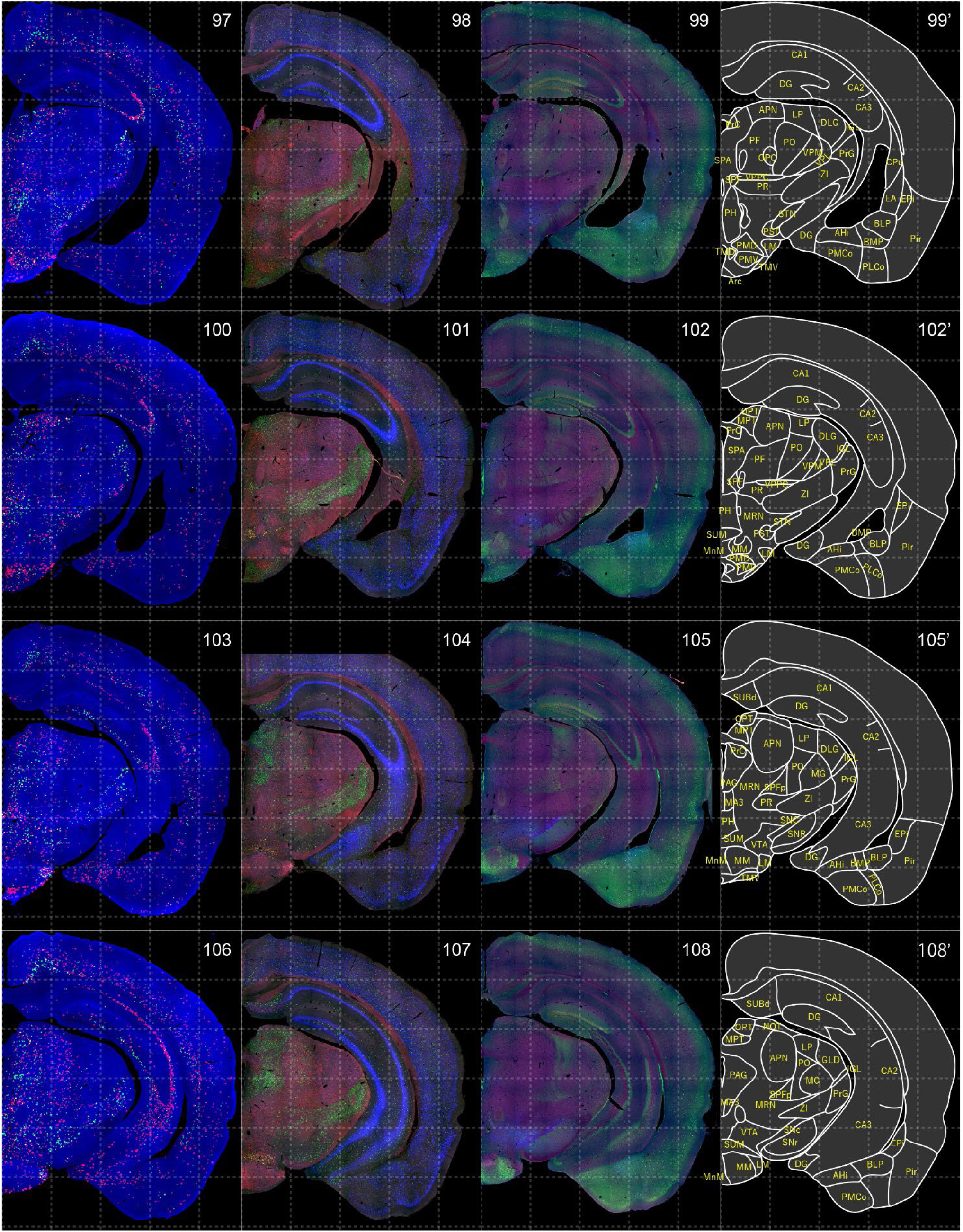
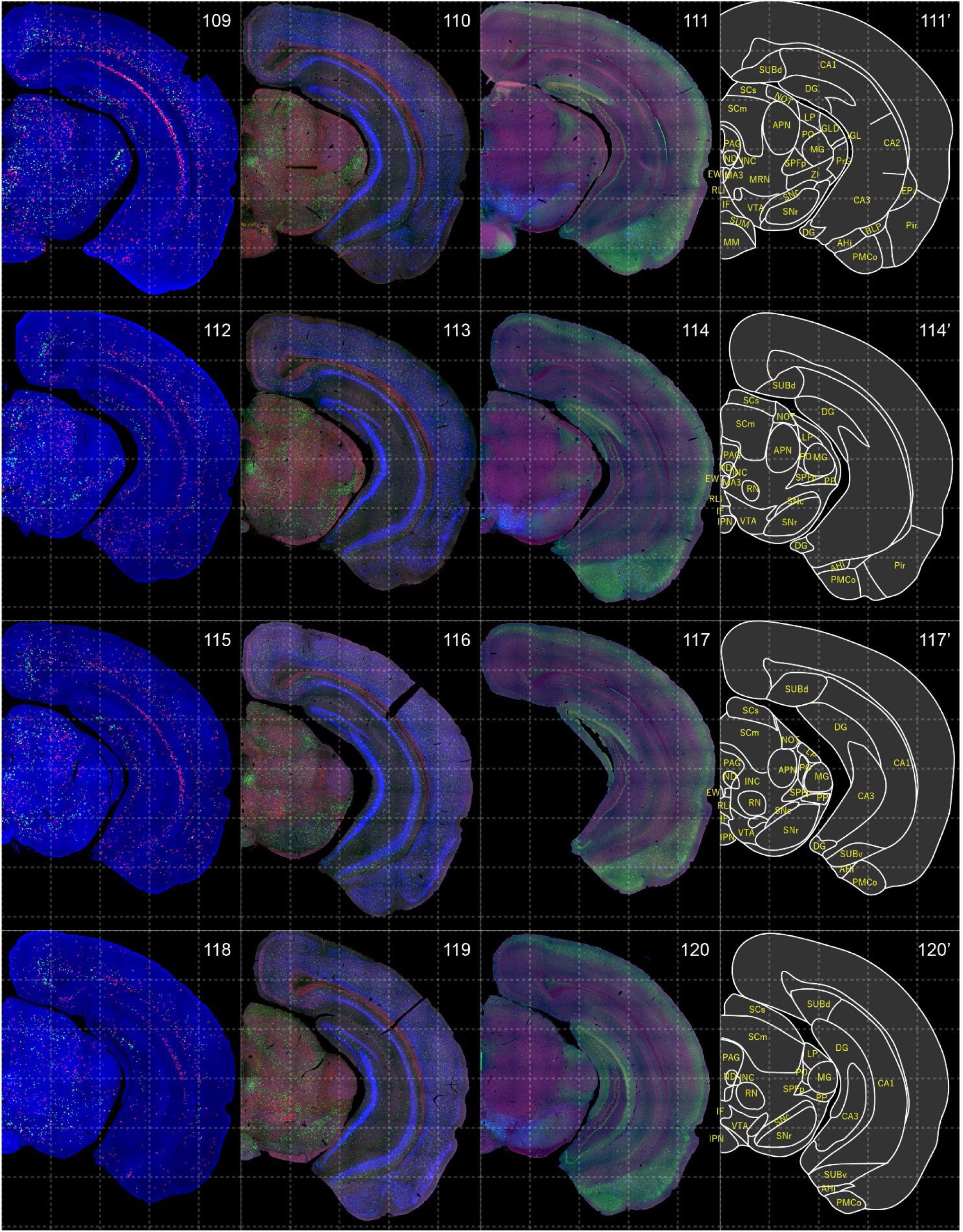
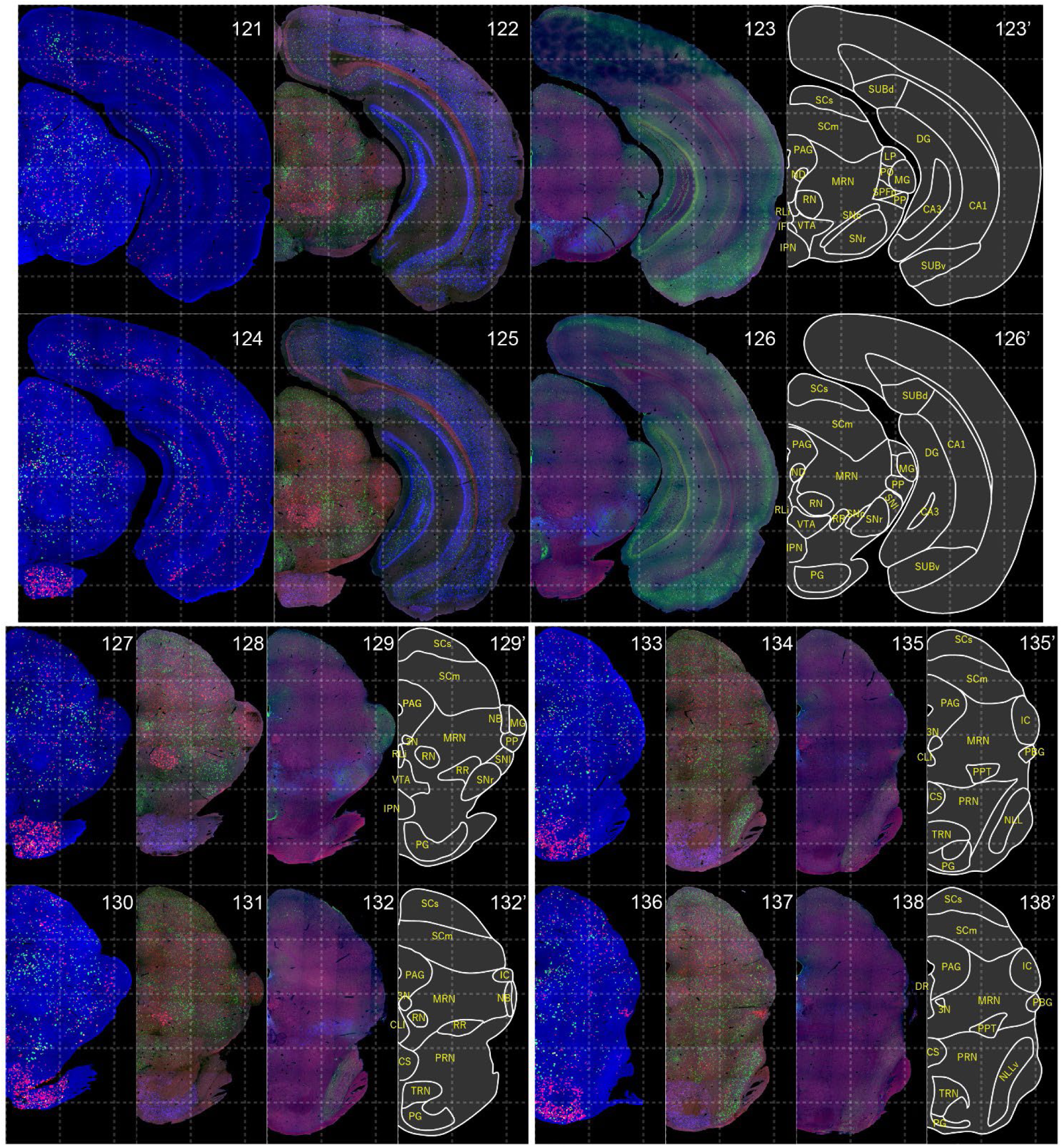
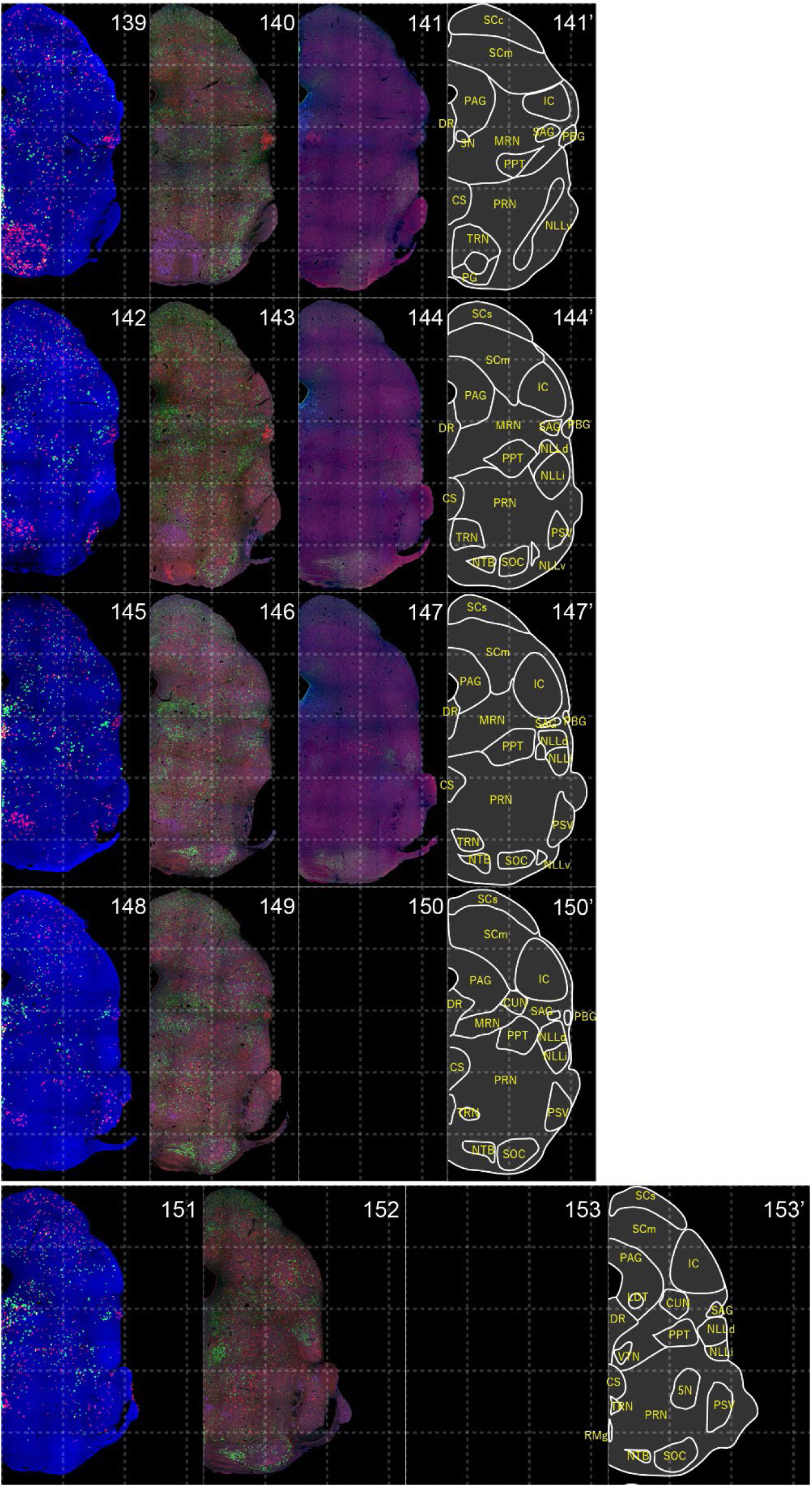
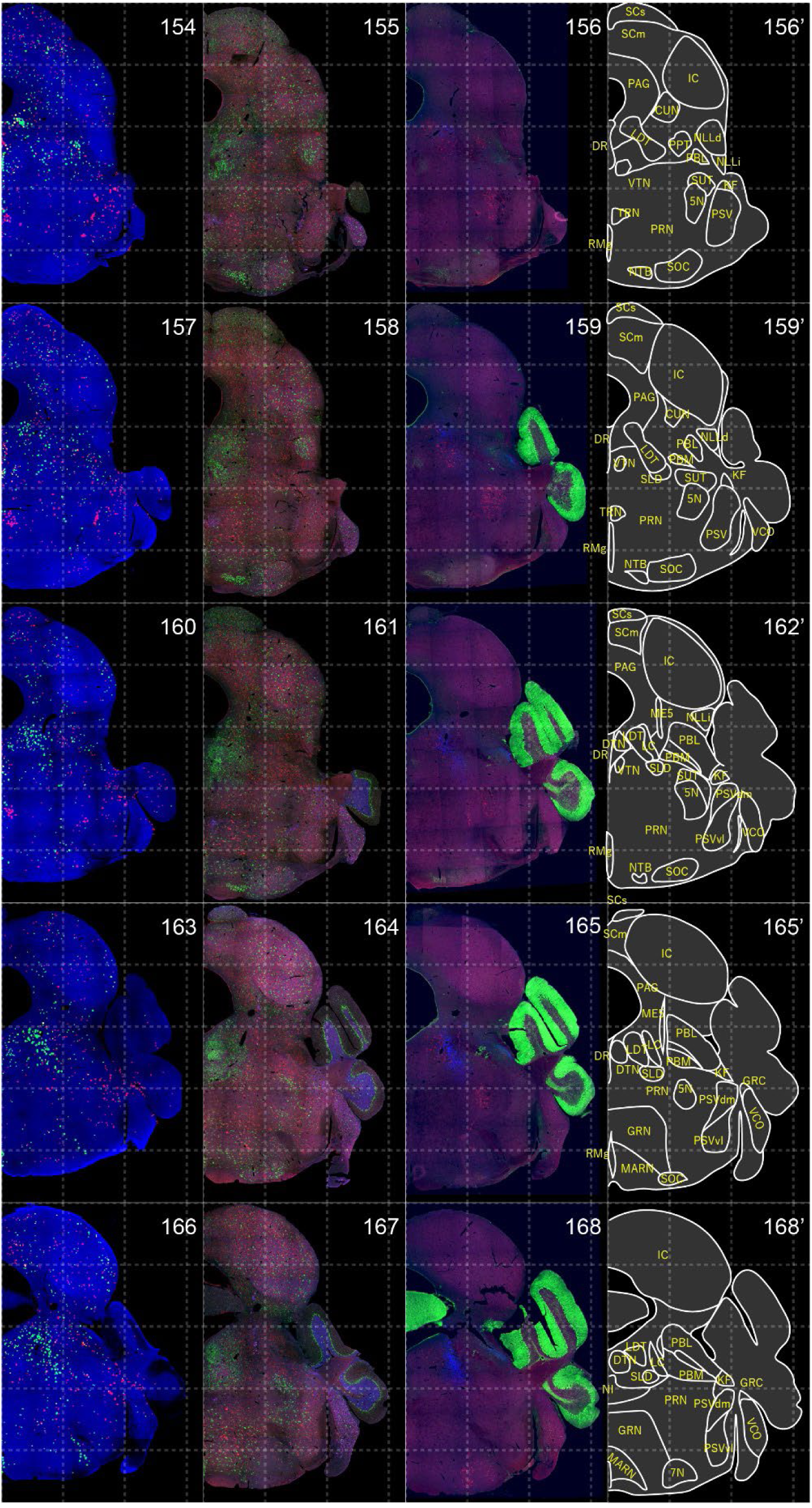
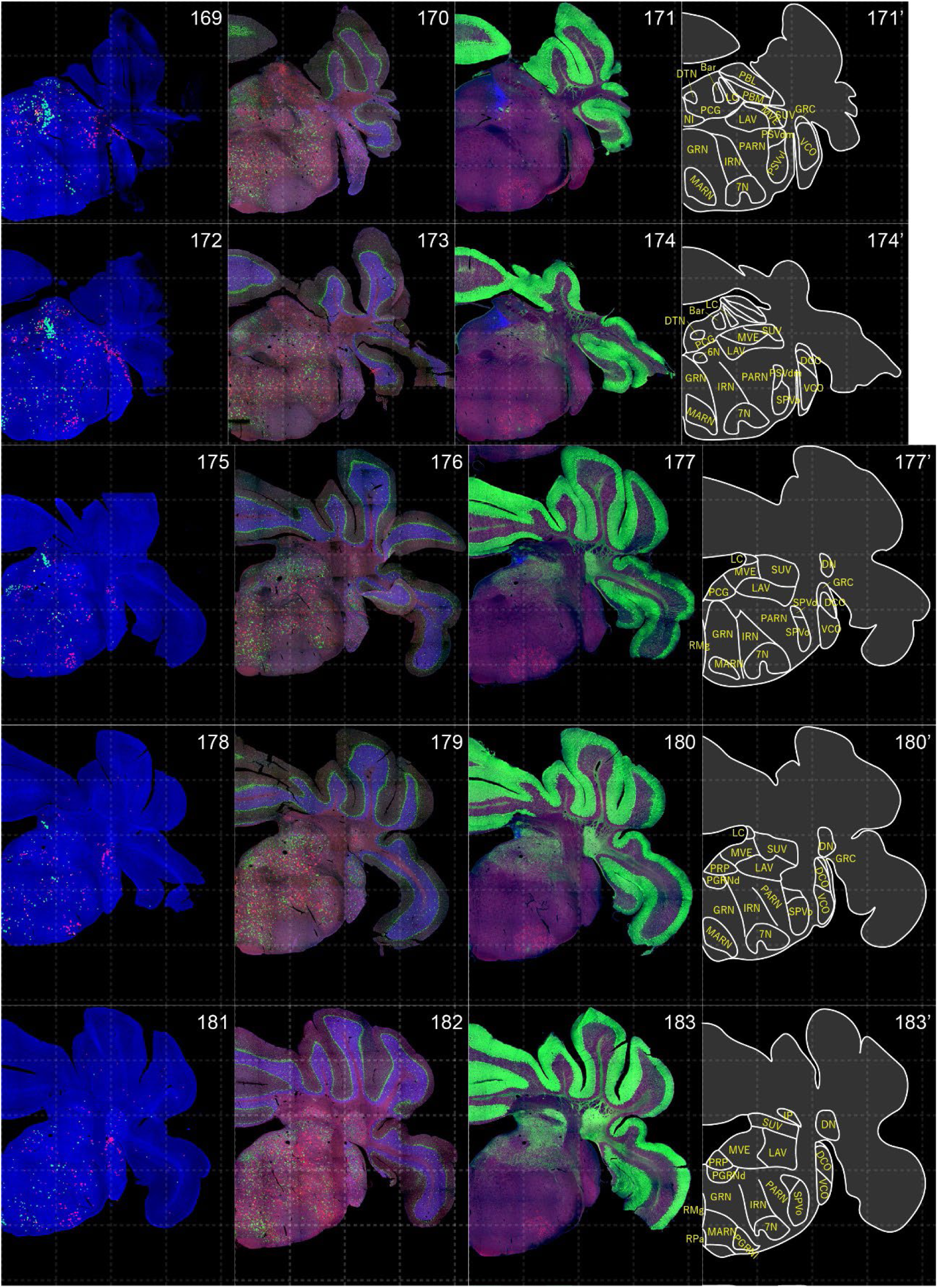
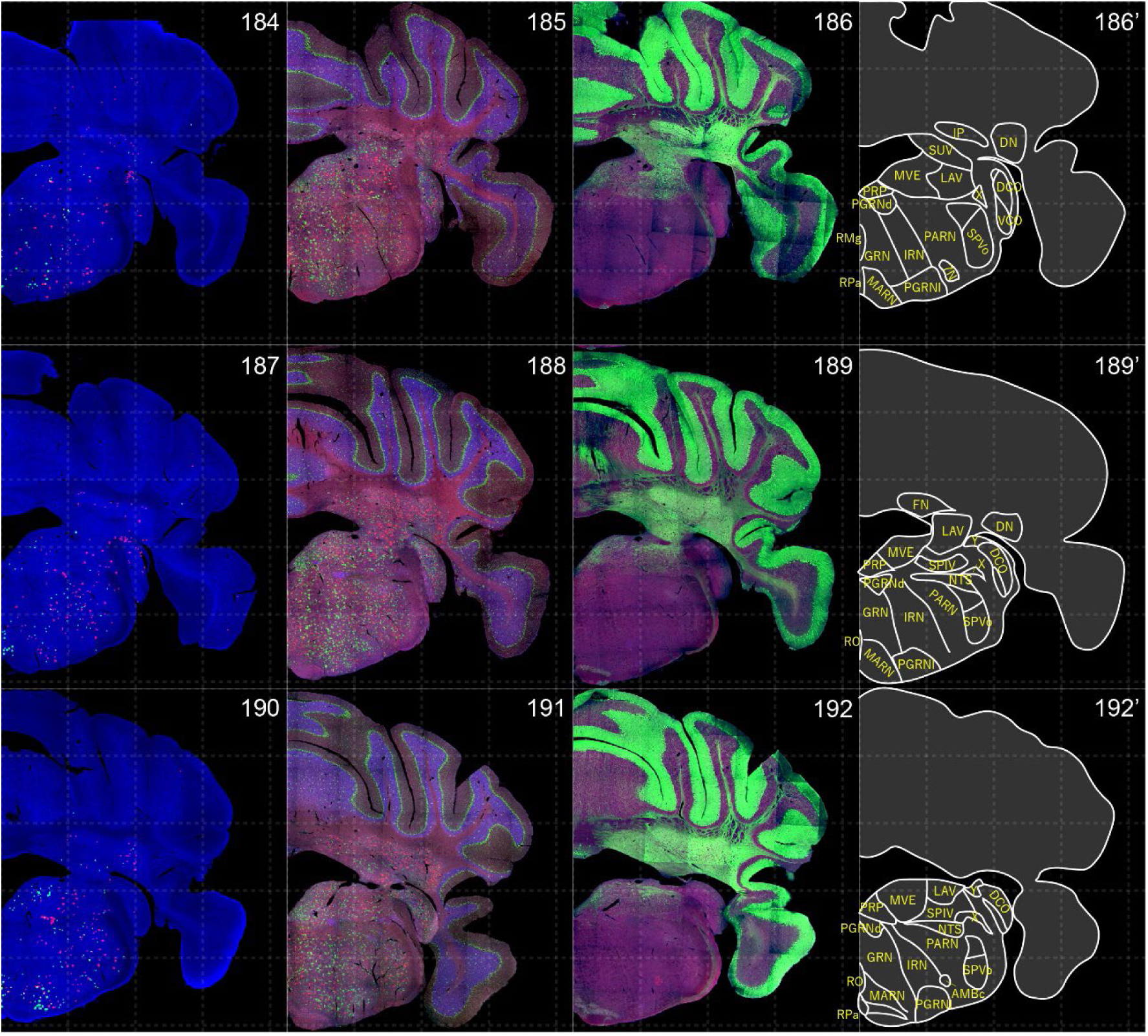
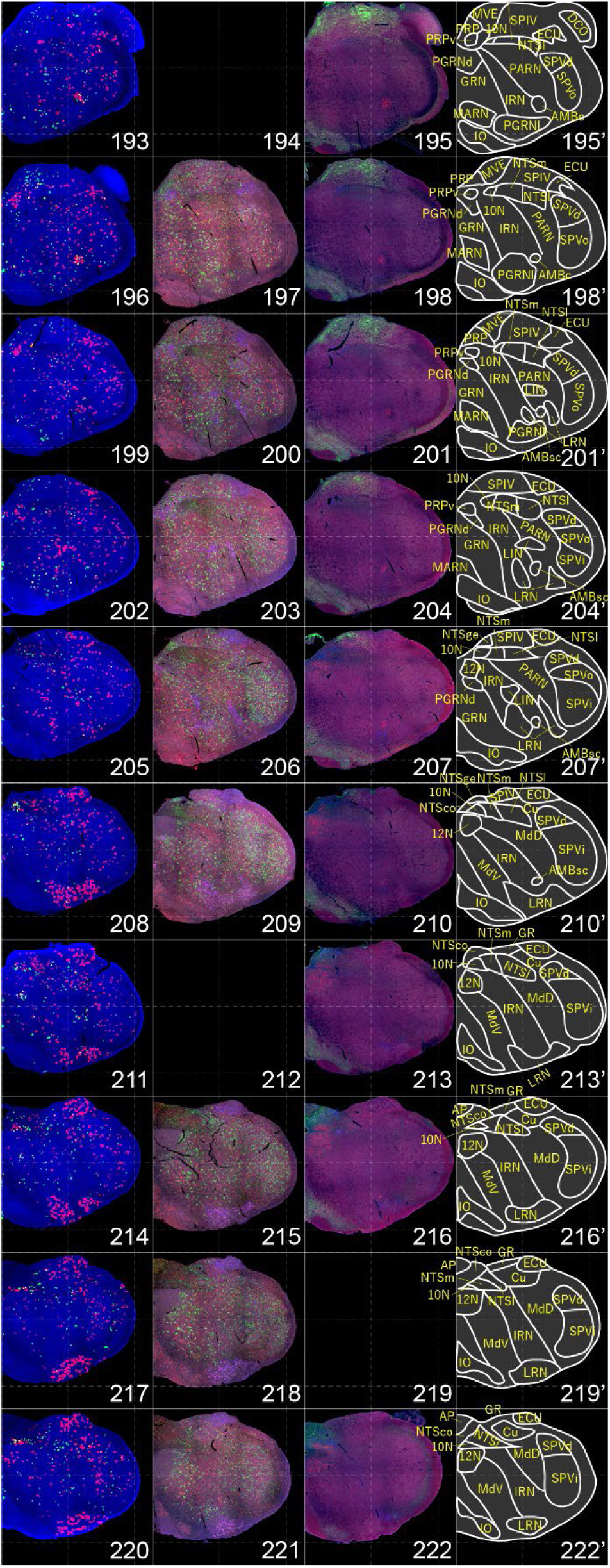
Distribution of orexin receptor-expressing cells and regional markers. Numbers show the anterior-posterior position of the complete serial sections with 40 μm thickness from a single mouse brain. Dashed-line 1mm grids overlapped. (A) Reconstructed distribution of the orexin receptor-expressing cells. Green and red dots indicate *Ox1r-* and *Ox2r*-expressing cells, respectively. (B) Photographs of *Vgat* (green), *Vglut1* (blue), and *Vglut2* (red) mRNA staining. (C) Photographs of calbindin (green) and TH (blue) immunostaining and *Chat* (red) mRNA staining. (D) Atlas of brain regions examined.

**Fig. S3.**
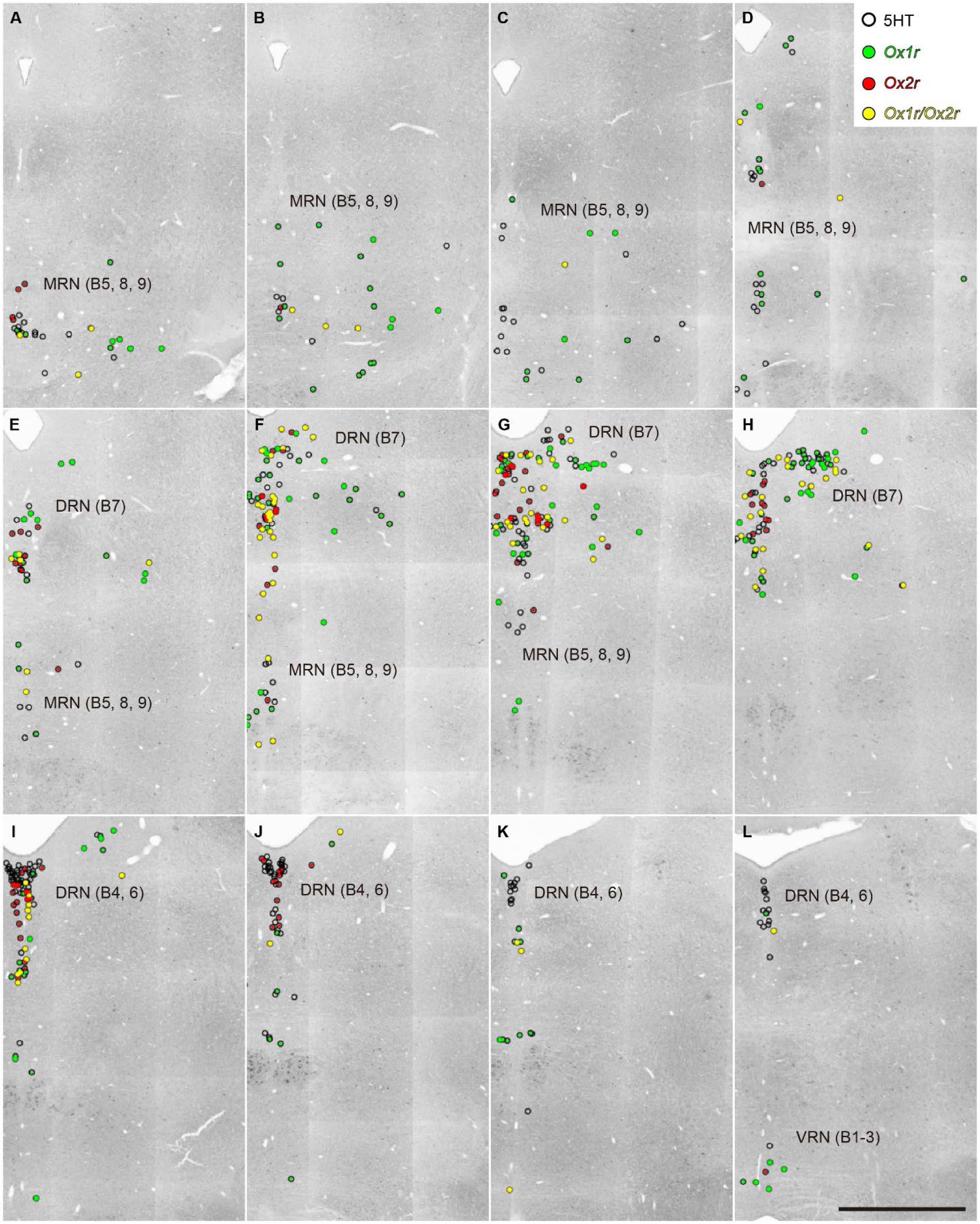
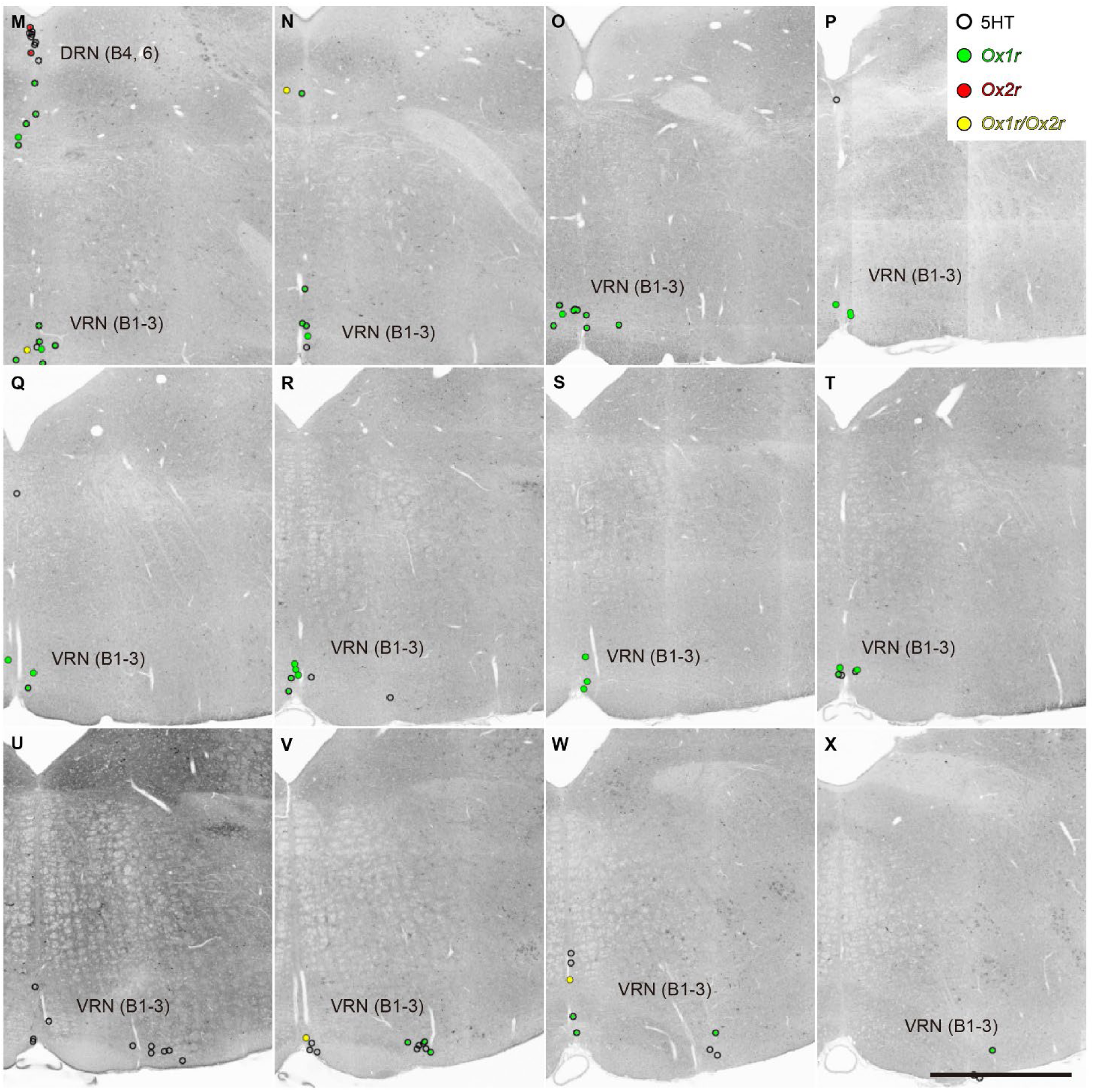
Distribution of orexin receptor-expressing serotonergic neurons. White, green, red, and yellow circles indicate receptor-negative, *Ox1r*-positive, *Ox2r*-positive, and both *Ox1r* and *Ox2r*-positive serotonergic neurons, respectively. Panels are arranged in anterior-posterior order. Scale bar: 500 μm.

**Fig. S4.**
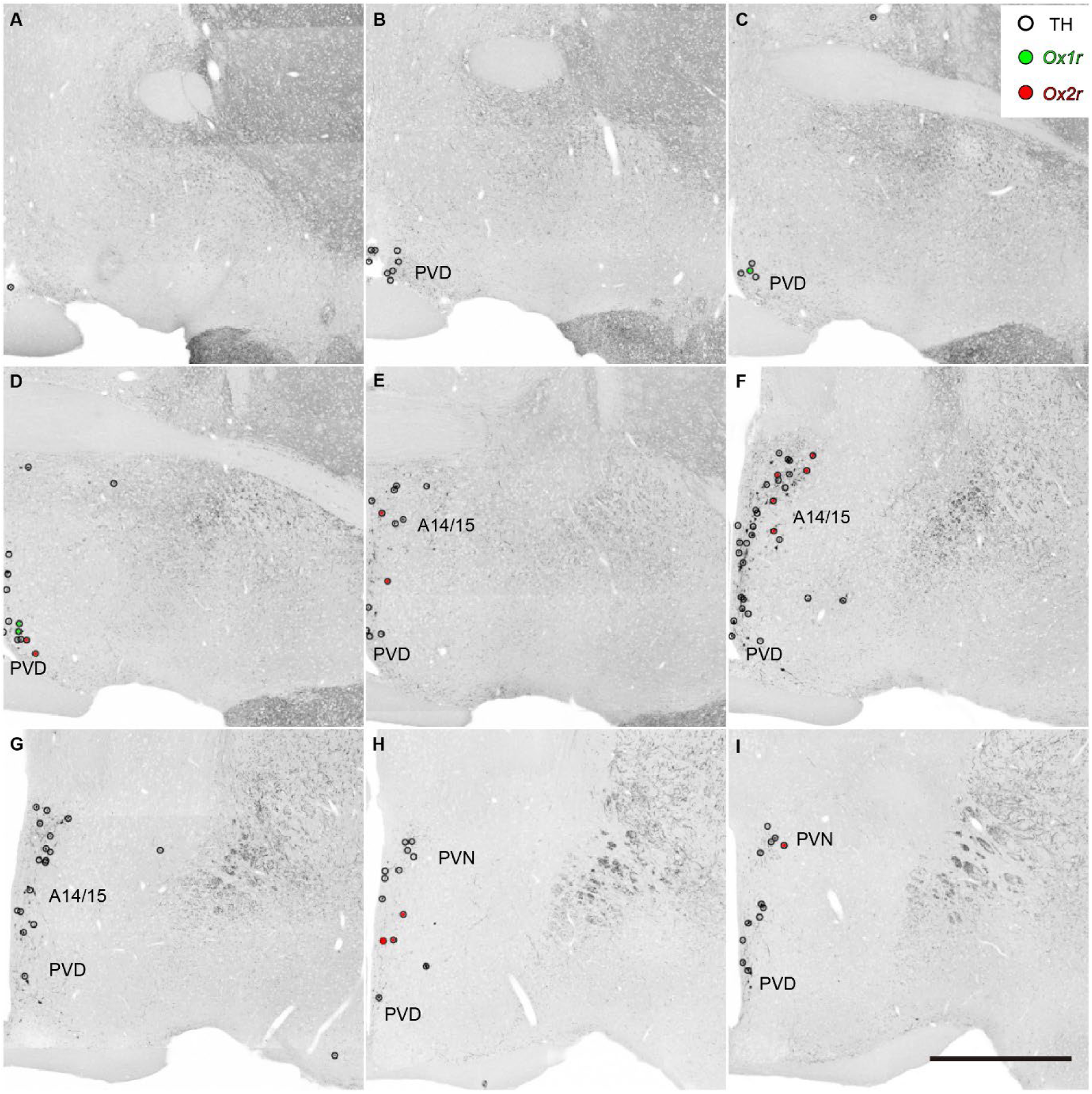
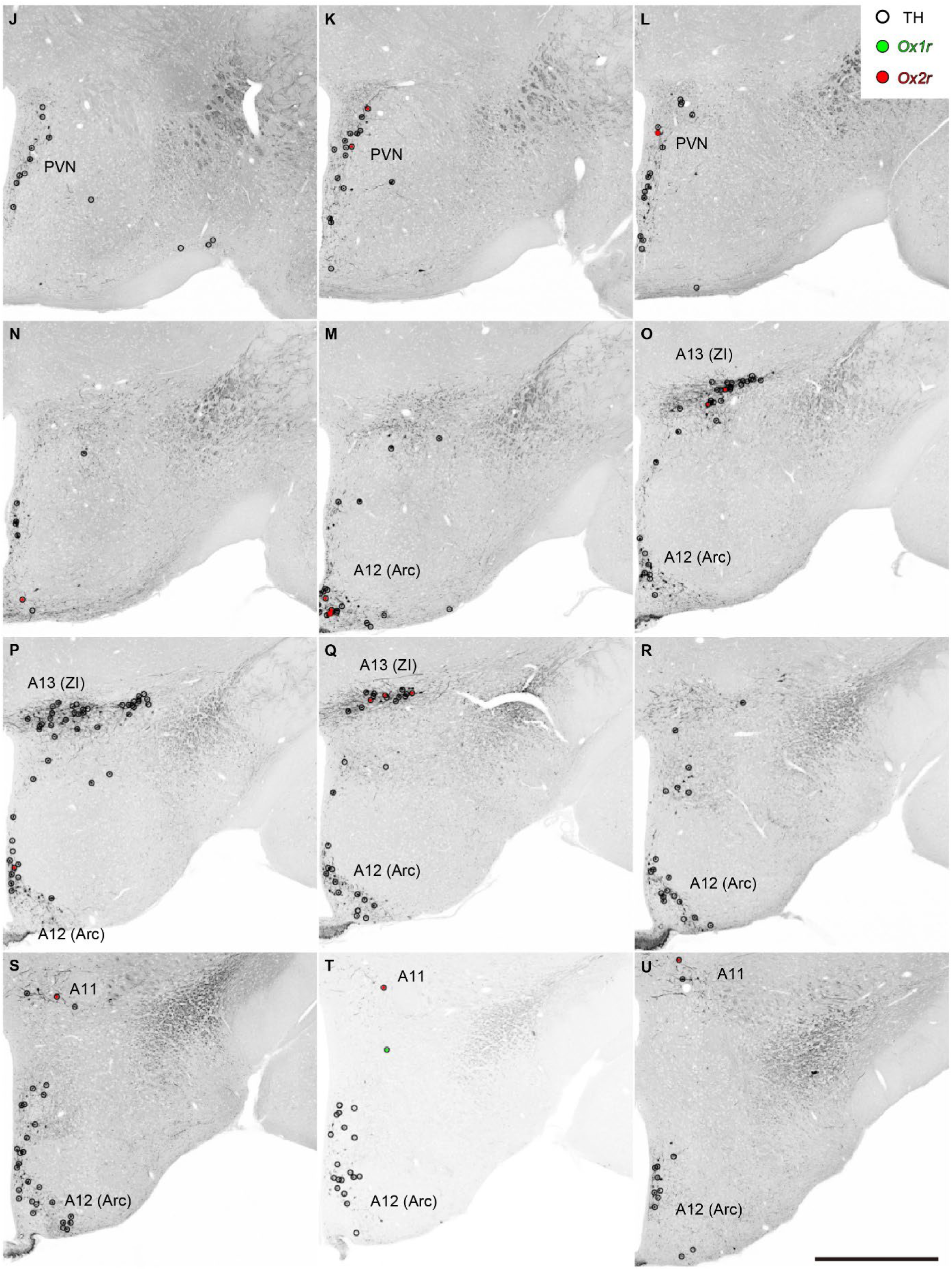
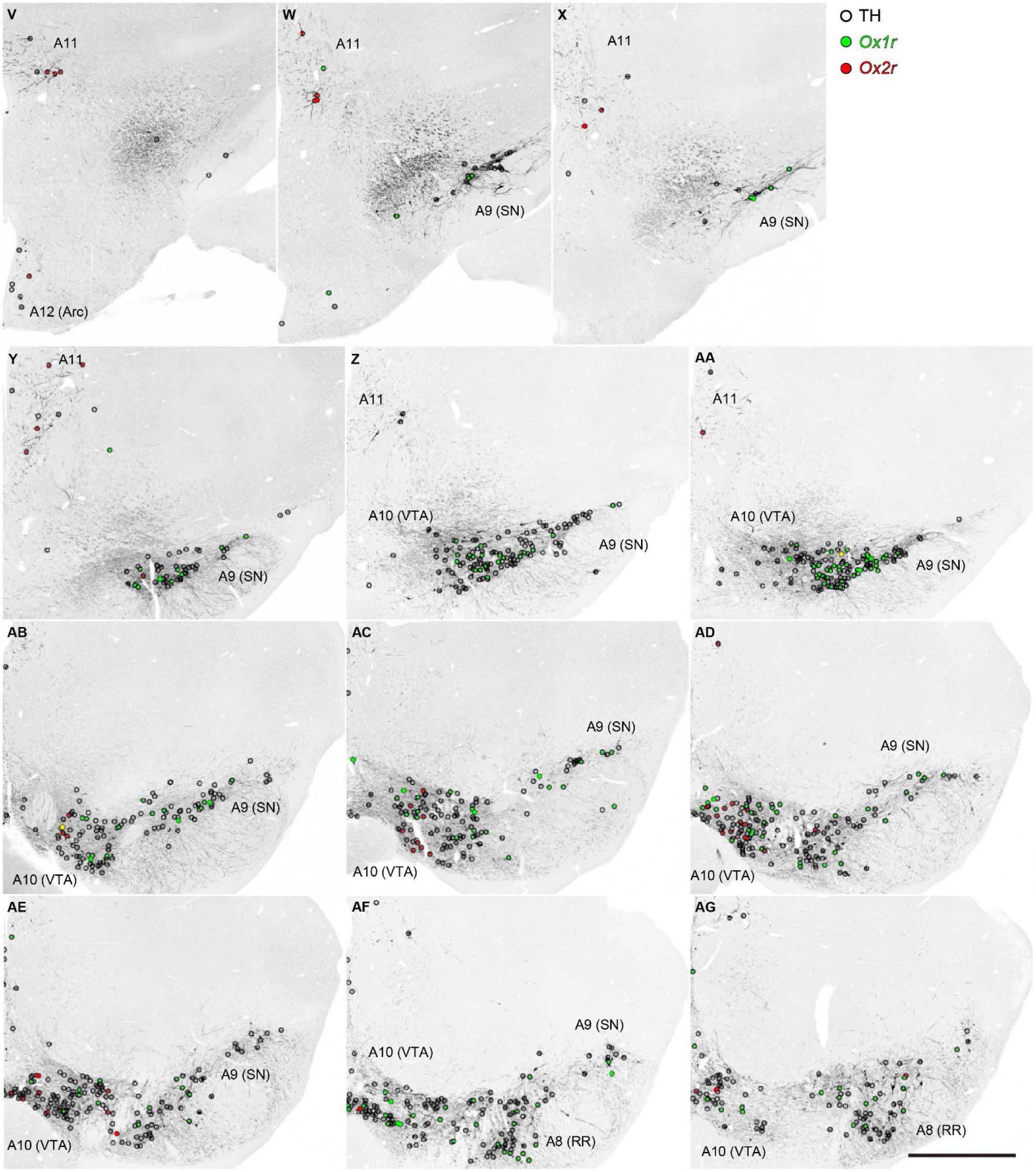
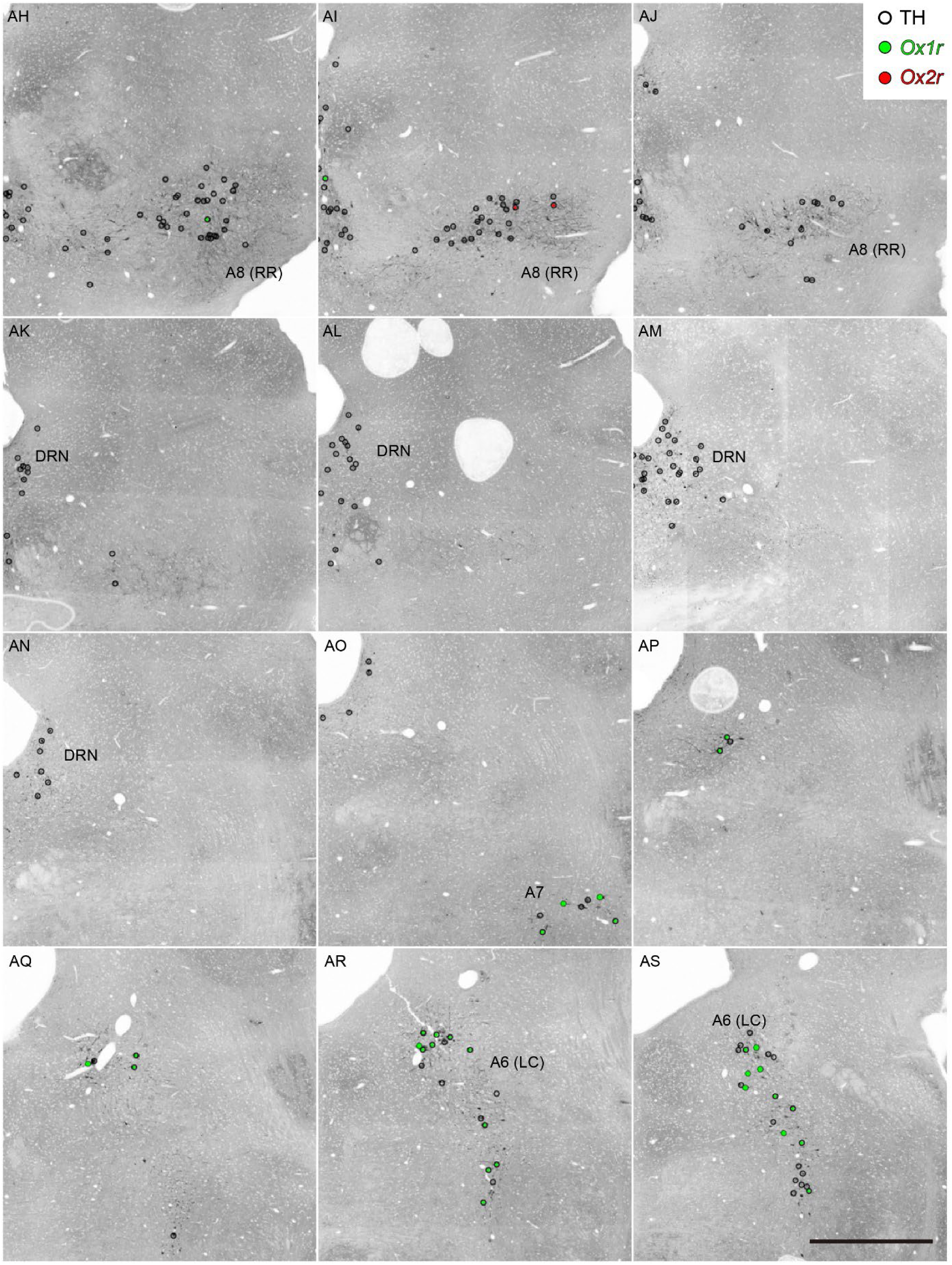
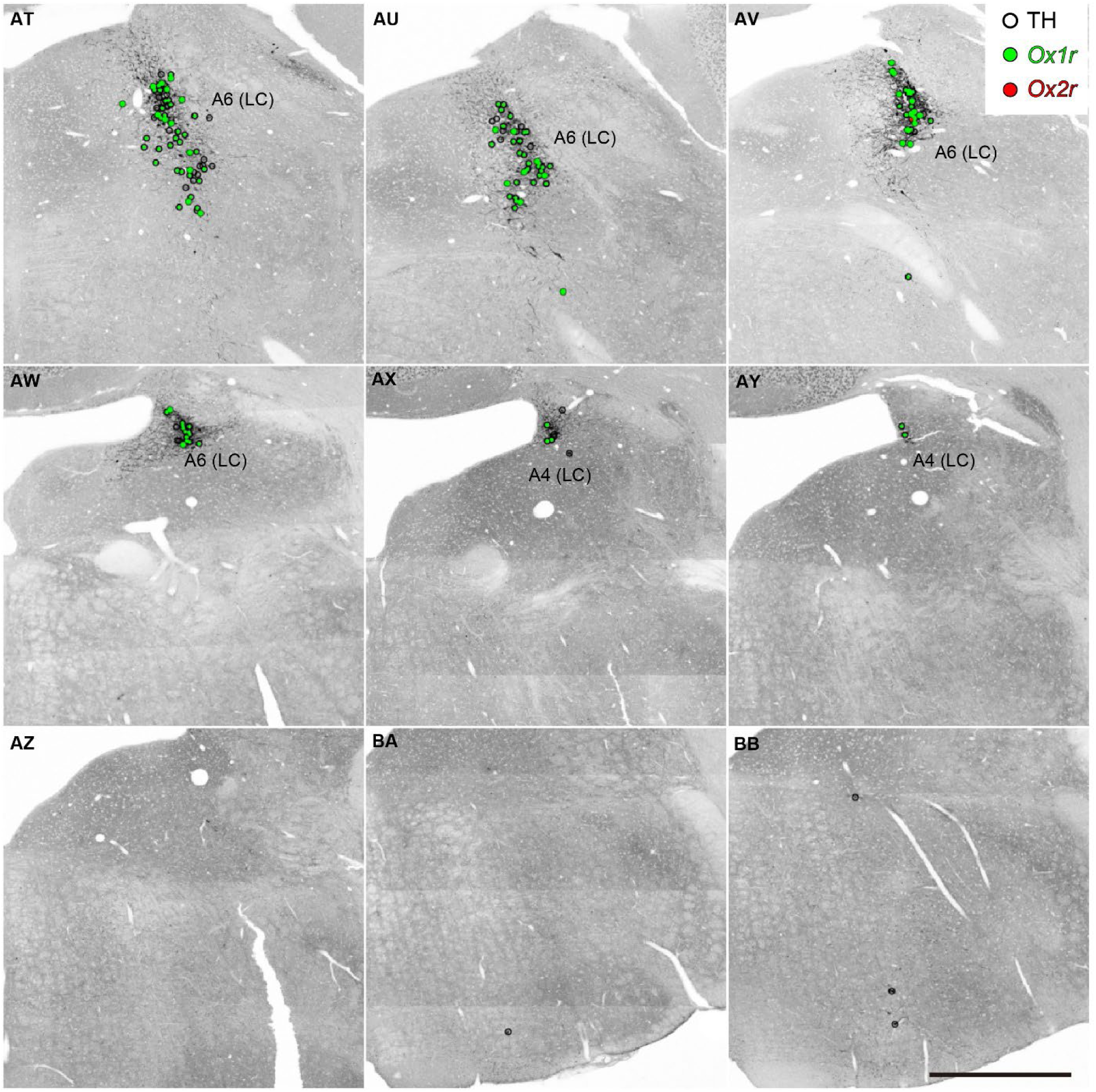
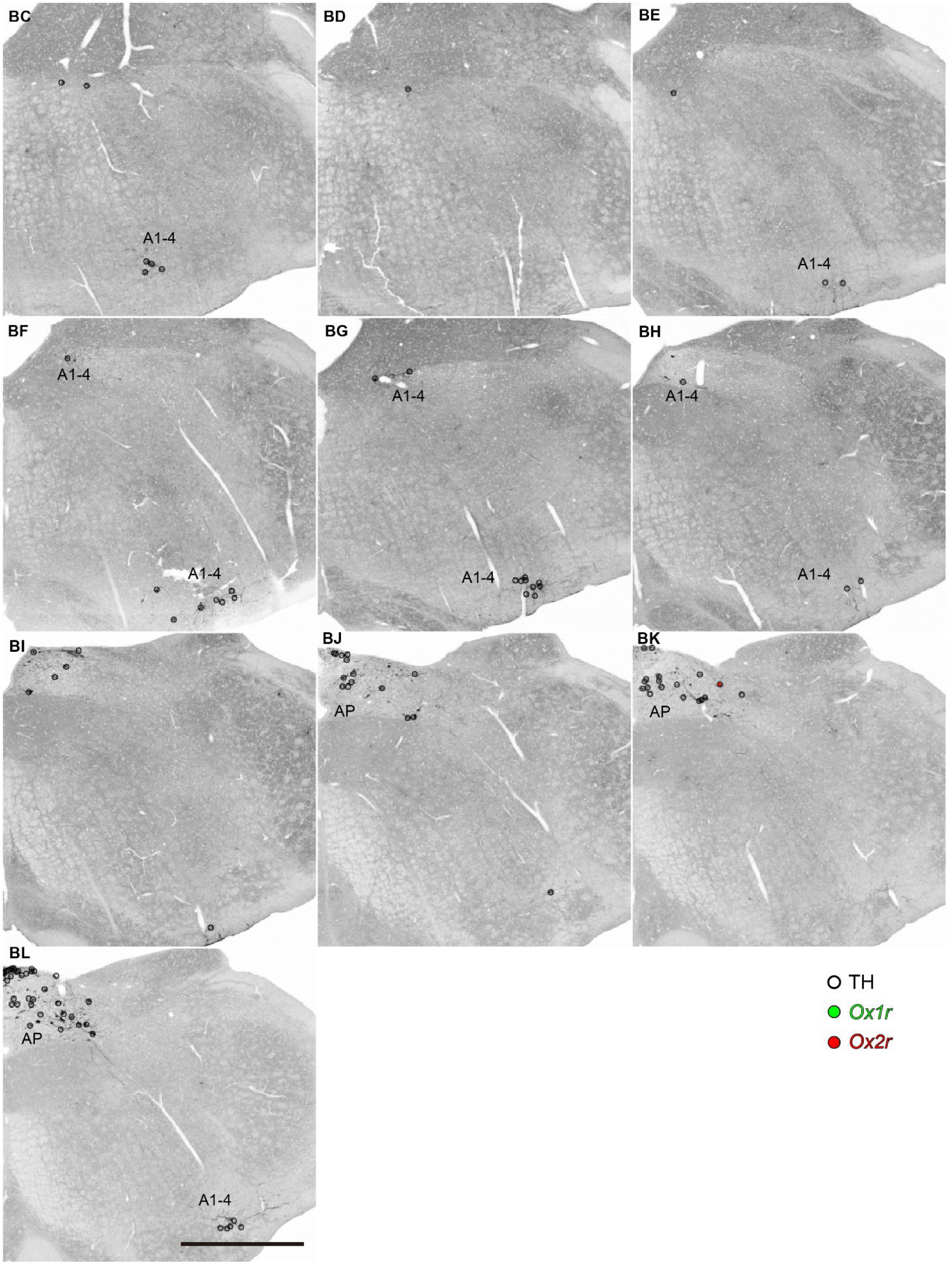
Distribution of orexin receptor-expressing dopaminergic/adrenergic neurons. White, green and red circles indicate receptor-negative, *Ox1r*-positive, and *Ox2r*-positive dopaminergic/adrenergic neurons, respectively. Panels are arranged in anterior-posterior order. Scale bar: 500 μm.

**Fig. S5.**
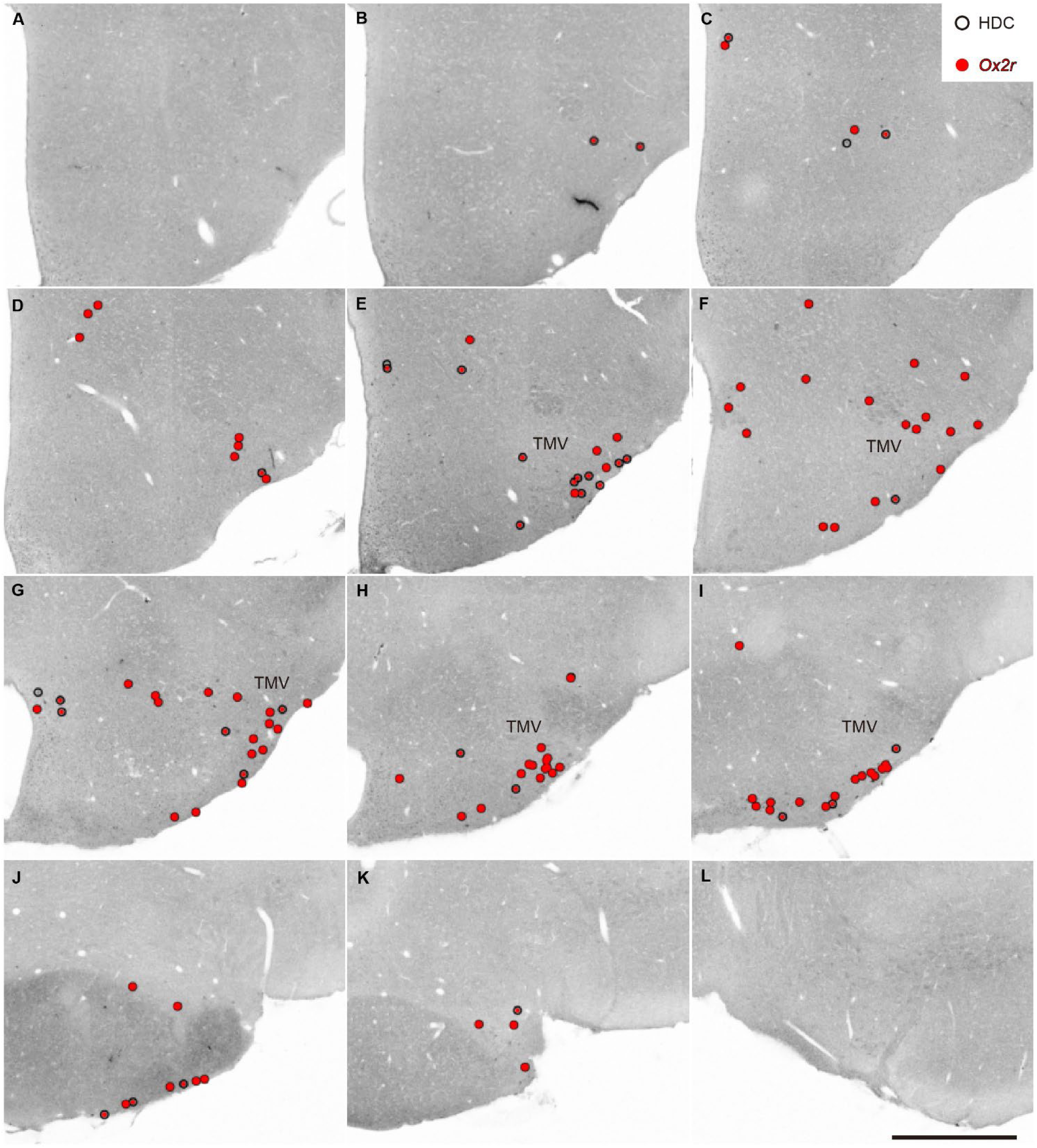
Distribution of *Ox2r*-expressing histaminergic neurons. White and red circles indicate *Ox2r*-negative and *Ox2r*-positive histaminergic neurons, respectively. Panels are arranged in anterior-posterior order. Scale bar: 500 μm.

**Fig. S6.**
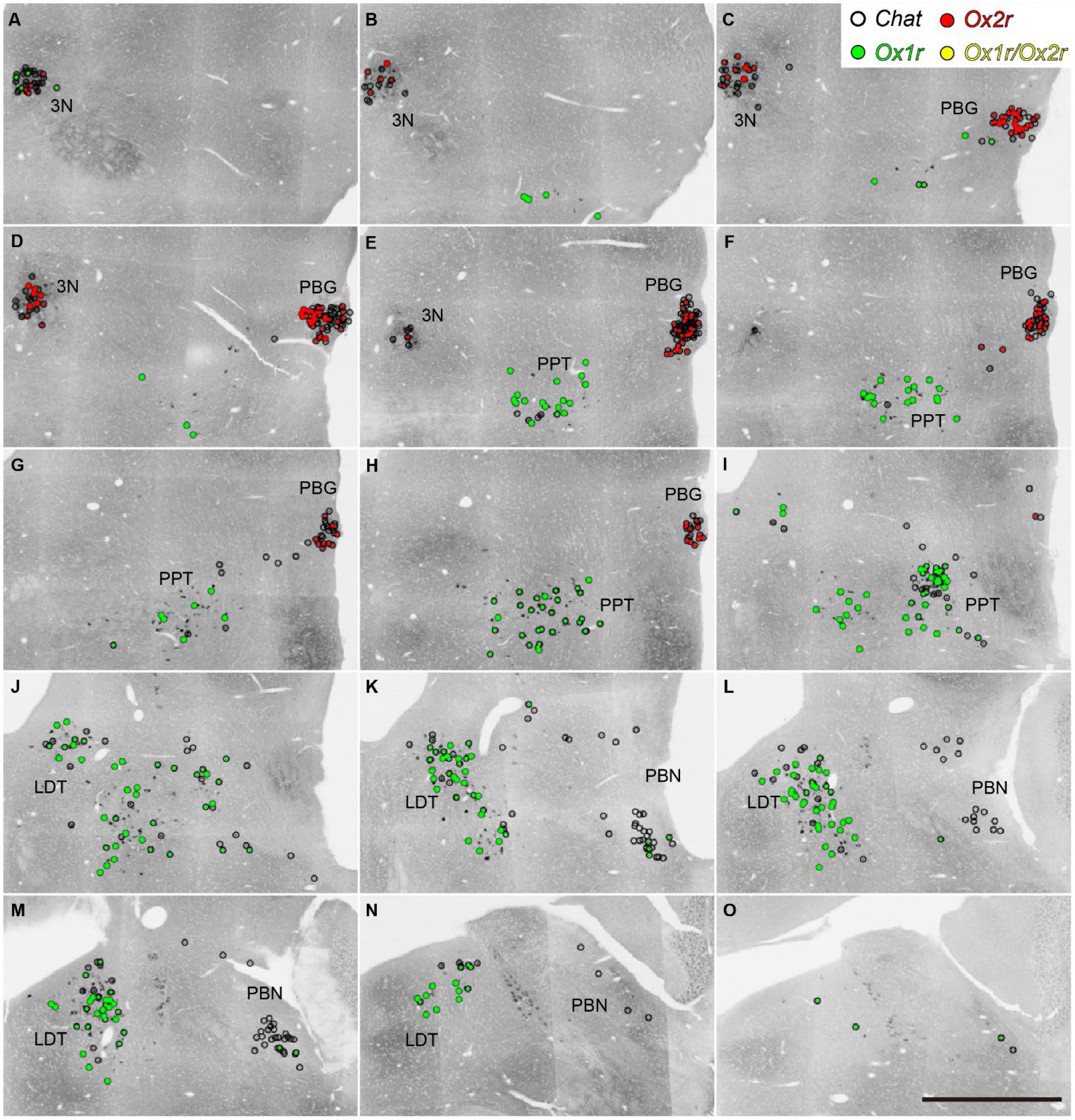
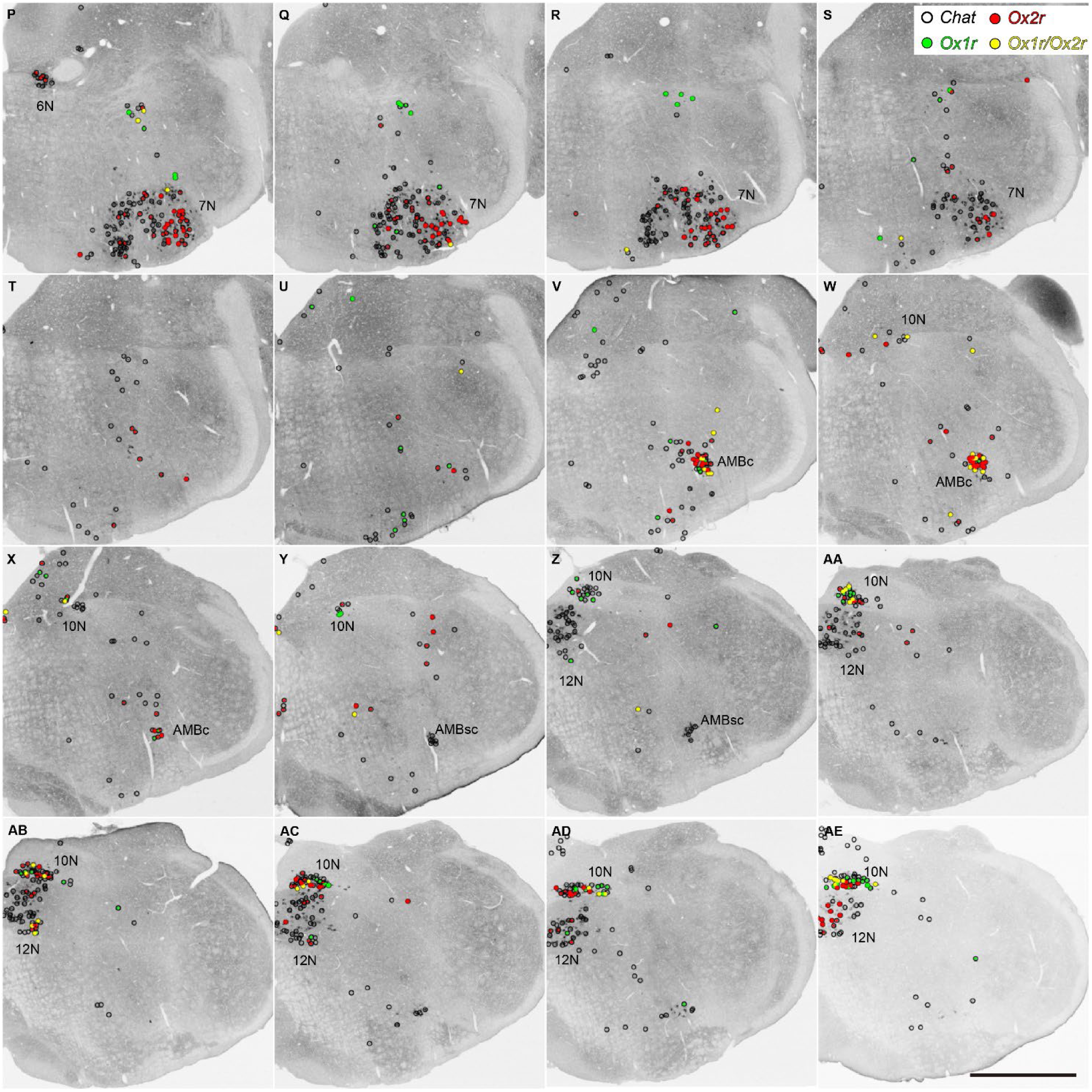
Distribution of orexin receptor-expressing cholinergic neurons in the brainstem. White, green, red, and yellow circles indicate receptor-negative, *Ox1r*-positive, *Ox2r*-positive, and both *Ox1r-* and *Ox2r-*positive cholinergic neurons, respectively. Panels are arranged in anterior-posterior order. Scale bar: 500 μm

## Notes

### Competing Interest Statement

The authors have declared no competing interest.

